# The WormFood CURE: Screening for bioactive metabolites that antagonize the *Caenorhabditis elegans* Ras signaling pathway

**DOI:** 10.64898/2026.06.03.729853

**Authors:** Emily Washeleski, Emily Morrin, Ryleigh Parsons, Christian Holmstrom, Megan E. Guyer, Amanda Bekkala, Paul D. Goetsch

## Abstract

Course-based Undergraduate Research Experiences (CUREs) provide an accessible, scalable platform for scientific discovery. Here, we present the WormFood CURE, which mines environmental bacterial isolates for bioactive secondary metabolites using *Caenorhabditis elegans* phenotype suppression as a functional readout. Utilizing the multivulva (Muv) phenotype, our pilot cohort interrogated 41 wild bacterial isolates for suppression of Ras/MAPK signaling. We identified one *Bacillus safensis* isolate BAC-08 and one *Bacillus altitudinis* isolate BAC-44 that significantly inhibited ectopic vulval precursor cell (VPC) induction in Muv strains when fed as a live food source. BAC-08 and BAC-44 also significantly affected wild-type nematode development and growth. Metabolic pathway reconstruction from annotated genome assemblies did not support nutritional deficiency as the potential mechanism; instead, we observed that methanol-soluble intracellular extracts from BAC-44 were sufficient to inhibit pseudovulvae growth. We concluded that the observed Muv suppression is likely driven by a secondary metabolite effect. Comparative genomic analysis further identified unique biosynthetic gene clusters (BGCs) present in both BAC-08 and BAC-44 isolates compared to the other isolated *Bacillus* species. Altogether, our study demonstrates that the WormFood CURE model successfully identifies novel bacterial-genetic interactions, providing a scalable platform for discovery of new natural microbial products that modulate conserved eukaryotic signaling pathways.

## Introduction

Microbial natural products, or secondary metabolites, remain a major source of new bioactive small molecules with therapeutic potential. Natural products and their derivatives account for approximately one-third of the ~1,900 new chemical entities approved through the year 2019, including 107 anticancer and 90 antibacterial drugs [1]. Roughly 16% of all FDA-approved natural products originated from bacteria [2], which likely underrepresents true bacterial chemical diversity given the reservoir of uncharacterized microbial taxa [3]. Unfortunately, the discovery rate of genuinely new chemical entities has decreased, driven in part by the high frequency of known-compound rediscovery [4]. Although high-throughput chemical library screening techniques and computational prediction approaches promise to accelerate the discovery of new compounds [5], establishing effective methods to identify bioactive compounds represented within the immense diversity of microbial species remains a challenge.

We propose a novel strategy to mine microbial natural products by leveraging the predatory-prey relationship between the bacterivorous model organism *Caenorhabditis elegans* and their microbial food source. *C. elegans* is a well-established model for host-microbe interaction investigations at both a molecular and systems level [6–9]. *C. elegans* natively inhabit soil pockets rich in decaying organic matter [10], which harbor highly diverse microbial communities [11–13]. Variation of bacterial food sources isolated from *C. elegans* native habitats revealed fitness benefits and costs beyond what can be explained solely by nutritional content, suggesting that microbial secondary metabolites affect nematode physiology even through feeding [13].

Unlike mammalian systems, *C. elegans* permits high-throughput screens of microbial isolates because both the target animal population and their live bacterial diet are more easily scalable [8]. As a proof-of-principle, we targeted the Ras/MAPK pathway, a major focus in drug discovery as the pathway is frequently dysregulated in cancer [14]. The *C. elegans* Ras/MAPK pathway is functionally orthologous to the mammalian pathway and overactivation causes a distinct multivulva (Muv) phenotype characterized by hyperinduction of vulval precursor cells (VPCs) that cause ectopic pseudovulva growth [15,16]. Classical genetic screens in *C. elegans* identified that overactivation of the Ras/MAPK pathway occurs through gain-of-function of the *let-60/Ras* gene [17,18] or through loss-of-function of two separate synthetic multivulva (SynMuv) class genes [19,20]. Pharmacological studies have targeted *C. elegans* the Ras/MAPK function and demonstrated chemical suppression of the Muv phenotype using the plant-derived natural product harmine, synthesized compounds, and traditional herb extracts [14,21–23]. Therefore, we hypothesize that if an isolated bacterial strain produces a bioactive secondary metabolite capable of suppressing Ras/MAPK signaling, then feeding the live bacterial isolate to *C. elegans* Muv strains will elicit an observable suppression of the phenotype.

Here, we present the identification of two wild bacterial isolates, one *Bacillus safensis* isolate BAC-08 and one *Bacillus altitudinis* isolate BAC-44, that when fed to Muv strains suppressed pseudovulva induction and growth. We identified our wild bacterial isolates through a pilot Course-based Undergraduate Research Experience (CURE) called WormFood. While similar established CUREs like SEA-PHAGES and Tiny Earth focus on viral and antibiotic discovery from soil [24,25], our WormFood CURE used *in vivo* functional genetics to explore how environmental bacterial isolates modulate *C. elegans* phenotypes. Follow-up experiments in the research lab revealed that two *Bacillus* isolates reproducibly suppressed the phenotype in *lin-52(bn131); lin-8(n2731)* (SynMuv) [26] and *let-60(n1046)* (Muv) [17] genetic backgrounds. While these wild bacterial isolates caused a slight developmental delay and stunted wild-type hermaphrodite size, our data suggest that these phenotypic effects were not a result of nutritional deficiency. Instead, intracellular metabolite extraction revealed that BAC-44 produces a methanol-soluble secondary metabolite that suppressed pseudovulvae growth. Altogether, our results demonstrate that the WormFood CURE approach provides an accessible and generalizable pipeline for the discovery of bioactive microbial natural products that can modulate conserved eukaryotic signaling pathways.

## Results

### Establishing the pilot WormFood CURE screen for bacterial natural products using *Caenorhabditis elegans*

Varying the *C. elegans* bacterial diet has helped define the organism’s nutritional requirements, elucidate host-pathogen interactions, and clarify the nematode’s ecological niche, but has never been used as a method to mine for bioactive secondary metabolites [6,13,27]. To begin our study, we established a pre-screening method to identify wild bacterial isolates of interest within a two-month Course-based Undergraduate Research Experience (CURE) called WormFood. Our course goals included isolating wild bacterial strains from student-supplied environmental samples, identifying each bacterial strain’s genus using 16S sequencing, and testing the phenotypic effects following feeding of each bacterial strain to various *C. elegans* genetic backgrounds. During the WormFood CURE activities, we tested bacterial toxicity to wild-type *C. elegans* and suppression of the multivulva (Muv) phenotype. Altogether, the WormFood CURE enabled quick evaluation of a diverse panel of random bacterial isolates while simultaneously engaging undergraduate students in a meaningful hypothesis-driven research project.

The CURE students isolated 41 individual bacterial strains from several environmental sources, including snow and ice, soil and compost, and other non-human and non-animal sample sources (Fig. 1a, Supplementary Table S1). Students logged colony macromorphology while we performed Gram staining of each isolate (Supplementary Fig. 1, Supplementary Table S1). Using 16S rRNA gene sequencing, students identified the genus of 37 of 41 isolates. We identified 14 different genera, with Gram-negative *Pseudomonas* (n = 10) and Gram-positive *Bacillus* (n = 9) emerging as the predominant groups (Fig. 1a). Two isolates remained unidentified because their 16S rRNA results did not match our gram staining results (Supplementary Fig. 1). Two additional isolates that failed the initial 16S rRNA gene sequencing run were later identified as *Bacillus*. Isolates were assigned unique identifiers based on their genus and the order of collection. Most of the sequencing results showed high 16S identity (>96%) to known reference sequences, except for BAC-39 (86.33%) (Fig. 1a). Altogether, our CURE activities yielded a phylogenetically diverse wild bacterial library spanning 14 genera for downstream phenotypic screens.

**Fig. 1.**
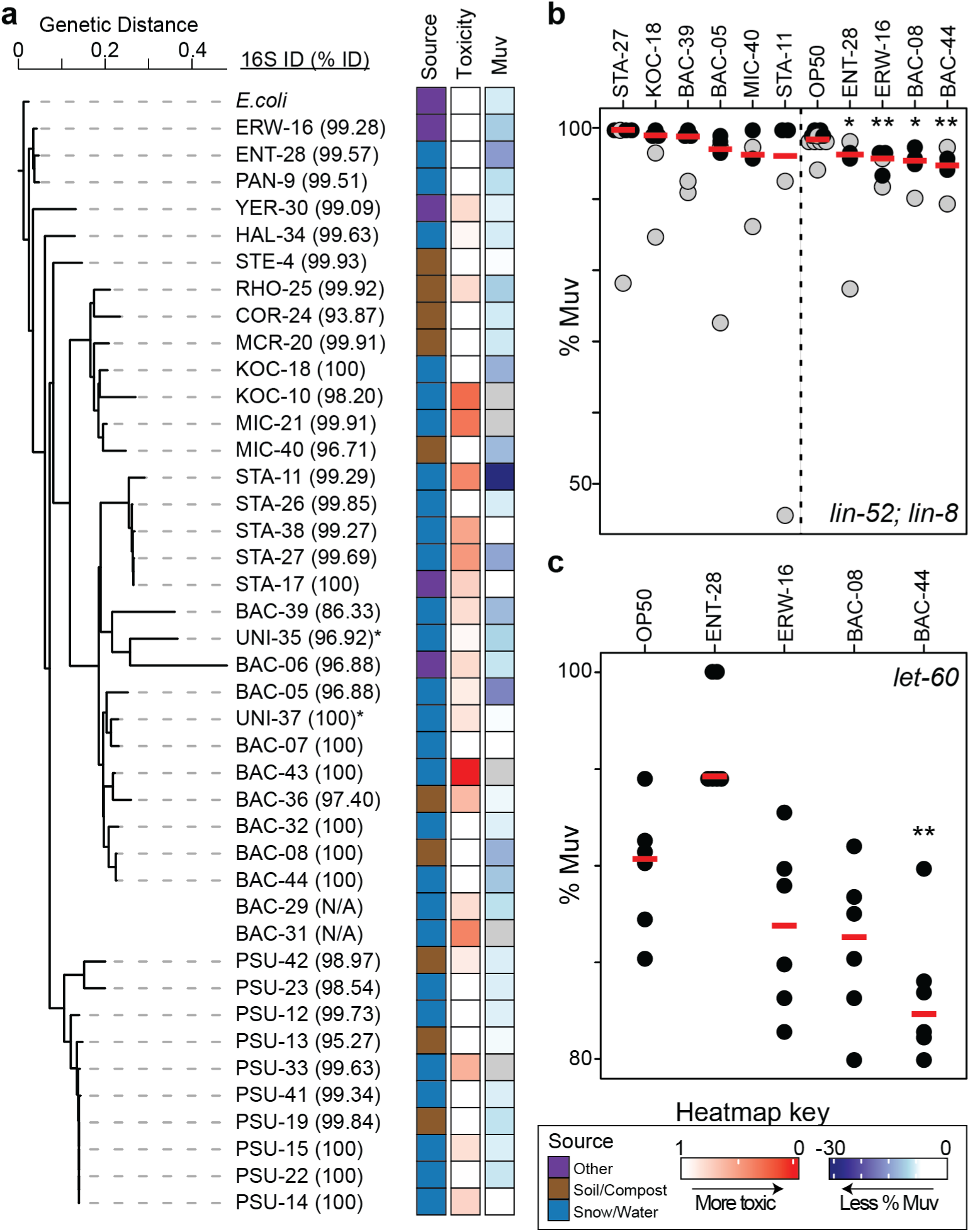
Phylogenetic diversity and bioactivity screens of bacterial isolates. (a) 16S rRNA gene phylogenetic tree of bacterial isolates with *E. coli* OP50 as the outgroup. The scale bar represents genetic distance (nucleotide substitutions per site). 16S identity (% ID) to closest reference sequence is indicated for each isolate. Asterisks (*) indicate identification did not match gram staining result. N/A indicates 16S PCR was not completed. Heatmap columns indicate isolation source (purple = other; brown = soil/compost; blue = snow/water), toxicity to wild-type *C. elegans* (red = more toxic), and effect of multivulval (Muv) suppression in *lin-52; lin-8* mutants fed the indicated isolate (blue = less % Muv observed). (b) Percentage of Muv worms observed in *lin-52; lin-8* mutants fed the indicated isolate. Dashed vertical line separates strains with no effect (left) from candidates showing reduced Muv induction (right). (c) Percentage of Muv worms observed in *let-60* mutants fed the indicated isolate. For (b-c), red horizontal bars indicate the median and the individual data points for each biological replicate, with grey points observed in the CURE and black points observed in the lab. Statistical significance was determined by a Wilcoxon-Mann-Whitney test with * p < 0.05 and ** p < 0.01 compared to OP50 control.

We first evaluated whether each bacterial isolate served as a viable food source for *C. elegans*. Our experimental goal was to screen out potential pathogens, inedible bacteria, isolates producing metabolites that repel or kill the nematodes, or isolates that cause detrimental physiological impacts on wild-type *C. elegans* [13,28,29]. We defined “toxic” as a general term for bacterial isolates that reduced the number of progeny reaching maturity from a timed egg-laying experiment, as compared to OP50 control. The WormFood CURE students observed that 5 of 41 wild isolates (12%) caused greater than 50% reduction in the number of adults compared to the OP50 control (Fig. 1a, Supplementary Fig. 2a). In the research lab, we performed a more careful follow-up using synchronization by bleaching on the 5 hits, BAC-43, KOC-10, MIC-21, BAC-31, and STA-11. We also re-evaluated PSU-33, BAC-36, BAC-29, and RHO-25 to test whether the initial CURE results were artifactual due to OP50 cross-contamination. We observed that larvae actively avoided isolates BAC-43, KOC-10, MIC-21 and RHO-25. Additionally, worms fed isolates BAC-31, STA-11, and BAC-36 exhibited severe developmental delay and growth suppression, while PSU-33 had no observable effect. Altogether, our toxicity assessments identified that BAC-43, KOC-10, and BAC-29 supported few worms to adulthood, and thus were removed from the WormFood CURE screen panel.

For our WormFood CURE screen, we performed a timed egg-laying experiment feeding using the *lin-52(bn131); lin-8(n2731)* SynMuv strain [26] on each wild bacterial isolate, asking students to count the number of wild-type and Muv adults following a 4-day incubation at 20°C (Fig. 1a, Supplementary Fig. 2b). Paired loss-of-function of the *lin-8* SynMuv A class and the *lin-52* SynMuv B class genes causes the ectopic upregulation of the LIN-3 Ras/EGF signaling peptide in the hyp7 hypodermal syncytium during L3 larvae development [26,30]. Ectopic LIN-3 expression stochastically induces the 3 tertiary vulval precursor cells (VPCs) to adopt a vulval cell fate through activation of the Ras/MAPK pathway, resulting in the Muv phenotype hallmark of 1-3 ventral protrusions called pseudovulva. Students assessed their own isolate and on additional random isolate, providing two biological replicates per isolate. The class observed apparent suppression of the *lin-52; lin-8* SynMuv phenotype in 10 out of 37 tested wild isolates (Fig. 1b, Supplementary Fig. 2b), including STA-11, BAC-05, ENT-28, STA-27, KOC-18, BAC-08, BAC-39, MIC-40, BAC-44, and ERW-16. Subsequent lab re-evaluation eliminated STA-11, BAC-05, STA-27, KOC-18, BAC-39 and MIC-40 (Fig. 1b), primarily because developmental delay masked the Muv phenotype or the initial student counts were inaccurate. Importantly, the ENT-28, ERW-16, BAC-08, and BAC-44 Muv suppression effects were replicated in the lab (Fig. 1b), suggesting that the bacterial isolates inhibit the Ras/EGF pathway and aberrant VPC induction.

We corroborated our observations and sought to identify the potential mode of action by testing additional SynMuv and Muv genetic backgrounds. We first tested the four isolates against the gain-of-function mutant *let-60(n1046)* Muv strain [17]. We observed that BAC-44 uniquely suppressed the *let-60* Muv phenotype (Fig. 1c), which places its mode of suppression at or downstream of Ras. We also tested the downstream Ets-domain repressor mutant *lin-1(n431)* strain [15]. Interestingly, we observed that only BAC-08 significantly suppressed the *lin-1* Muv phenotype (Supplementary Fig. 3), suggesting that BAC-08’s mode of action may operate within the downstream transcriptional program activated by the Ras/MAPK signaling pathway. Finally, none of the isolates suppressed the highly penetrant Muv phenotype in the SynMuv mutants *lin-15AB(n309)* and *lin-8(n111); lin-9(n112)* (Supplementary Fig. 3) [16,31], suggesting that the bacterial suppression activity is insufficient to counteract ectopic LIN-3 expression in these backgrounds. Altogether, the outcomes of the pilot WormFood CURE demonstrate that the feeding of wild bacterial isolates to Muv and SynMuv strains is a viable experimental approach for identifying bacterial isolates that may harbor chemical suppressors of the Ras/MAPK pathway.

### Bacillus isolates-of-interest stunt growth and moderately affect C. elegans larval development

One concern from our WormFood CURE results was the possibility that suppression of the Muv phenotype may stem from indirect physiological factors rather than the action of a bioactive secondary metabolite. We observed generally that worms fed *Bacillus* isolates were smaller and larval development was delayed. We measured this effect in the lab using wild-type N2 strain fed OP50, ERW-16, BAC-08, and BAC-44. We observed that worms fed ERW-16 were significantly longer but with no change in width, whereas worms fed BAC-08 or BAC-44 were significantly shorter and thinner (Fig. 2a-c). Importantly, the reduction in body size on BAC-08 and BAC-44 was strictly proportional. We observed no significant change in the width-to-length ratio compared to OP50 (0.0652 ± 0.0008, n = 60) for either BAC-08 (0.0641 ± 0.0010, n = 42) or BAC-44 (0.0663 ± 0.0009, n = 52), values represent mean ± standard error of the mean (SEM). Since feeding defects can lead to shorter body lengths [32], our findings raised concerns that the *Bacillus* isolates were nutritionally deficient and thus potentially confounding our Muv suppression screen.

**Fig. 2.**
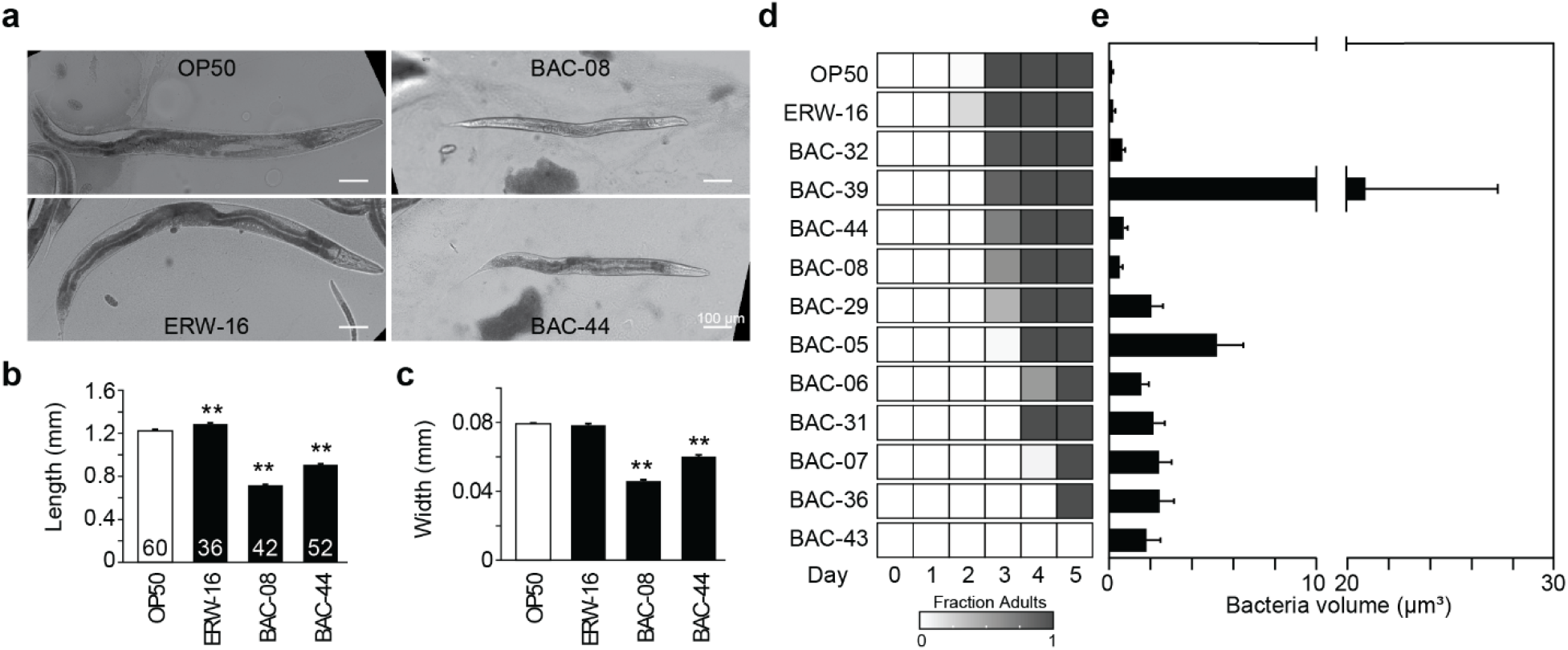
Impact of wild bacterial isolates on *C. elegans* growth and development. (a) Representative DIC micrographs of live adult hermaphrodites fed *E. coli* OP50, ERW-16, BAC-08, or BAC-44. Scale bar = 100 µm. (b-c) Bar graphs of (b) body length and (c) body width measurements of wild-type adult hermaphrodites fed the 4 diets. Total number of individuals measured (n) for each condition is shown within each length bar. Error bars represent standard error of the mean (SEM), and statistical significance was determined by a student’s T-test with ** p < 0.01 compared to OP50. d) Heatmap of larval development rate when fed each wild bacterial isolate depicted as the proportion of the population reaching adulthood (gradient from white = 0 to dark grey = 1) following a day 0 bleach synchronization. e) Bar graph of bacterial cell volume µm^3^ based on gram stain experiments. Data represent the mean volume calculated from n = 20 individual bacterial cells per strain, with error bars indicating the standard deviation.

Chronic food restriction also causes delay in larval development [33], so we evaluated the rough timing of larval development of all isolated *Bacillus* strains and ERW-16 by counting the number of adults on a plate each day following a day 0 bleach synchronization. On OP50, we observed that the entire population reached adulthood by the third day (Fig. 2d). Interestingly, ~20% of the population fed ERW-16 reached adulthood by the second day. We observed little difference in larval development timing compared to OP50 in worms fed BAC-32 and BAC-39. BAC-44 and BAC-08 caused a mild developmental delay with ~60-70% worms reaching adulthood by the third day. The rest of the *Bacillus* strains caused a more pronounced developmental delay, from ~40% of the population fed BAC-29 reached adulthood by the third day, to BAC-05, BAC-06, and BAC-31 delaying development by one day, to BAC-07 and BAC-36 delaying development by two days. Worms fed BAC-43 never reached adulthood, replicating our earlier toxicity results. Altogether, these results demonstrate that the BAC-08 and BAC-44 effects on larval development were not as pronounced compared to other *Bacillus* isolates in our panel.

One hypothesis for *Bacillus* isolates specifically being detrimental for *C. elegans* health is that the size of the gram-positive bacteria are too big for the *C. elegans* pharyngeal pump to break down effectively, thus slowing their feeding behavior [34]. We tested this hypothesis by correlating the bacterial cell volume from our WormFood CURE gram stain results (Supplementary Fig. 1) to our developmental delay results. Interestingly, we observed no correlation between bacterial volume and the severity of developmental delay. The most drastic example is that the largest *Bacillus* isolate, BAC-39, had little effect on larval development. Altogether, our results suggest that developmental delay is not linked to the size of the bacterial cells.

Finally, we evaluated whether the *Bacillus* strains were genetically capable of producing the appropriate nutritional components [35]. We sequenced, assembled, and annotated ERW-16 and each *Bacillus* isolate’s genome and assessed completeness of the biosynthetic pathways for essential amino acids and nutrients using KEGGdecoder and KEGGmapper (Fig. 3, Supplementary Fig. 4). Full genome sequencing of our isolates also allowed us to identify our strains to the species level (Supplementary Table S2), revealing that BAC-08 as a *Bacillus safensis* strain and BAC-44 as a *Bacillus altitudinis* strain. Overall, the biosynthetic pathway profiles for all strains did not reveal genetic deficiency, as the completeness for most of the pathways in BAC-08 and BAC-44 were at or exceeded the value observed in other isolates that don’t inhibit development. The one exception was BAC-08 and BAC-44 lacked a complete vitamin B7 biotin biosynthetic pathway. However, we do not expect that biotin deprivation caused the observed physiological effects because of nutrient carry-over from the rich LB culture medium used to grow each isolate [36]. Altogether, the metabolite pathways that may contribute to growth delay [35], including the heme, thiamine (B1), and cobalamin (B12) pathways were functionally complete or comparable to OP50 (Fig. 3, Supplementary Fig. 4). Altogether, we concluded that the worm growth and larval development defects observed in worms fed BAC-08 and BAC-44 are not likely caused by nutritional deficiency.

**Fig. 3.**
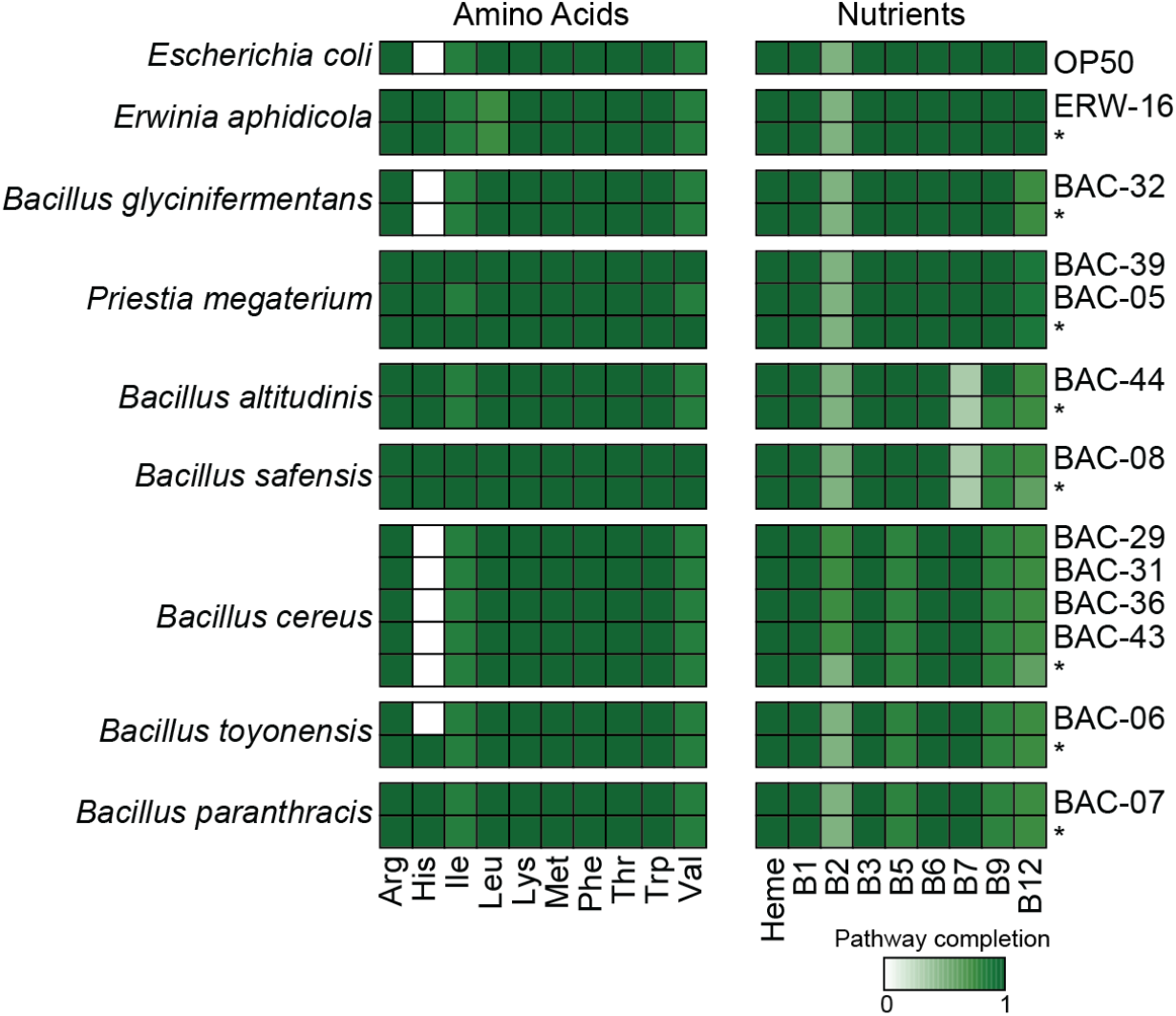
Metabolic potential of wild bacterial isolates for essential amino acids and nutrients. Heatmap analysis showing the predicted biosynthetic pathway completion (gradient from white = 0 to dark green = 1) for essential amino acids (left) and nutrients (right) identified from KEGGmapper and KEGGdecoder analysis of the genome assembly of each wild bacterial isolate. An asterisk (*) indicate analysis of the reference genome for the indicated bacterial species identified by GTDB-Tk.

### Bioactive secondary metabolites from wild bacterial isolates suppress pseudovulva induction and growth

Our preliminary screen identified that our wild isolates BAC-08 and BAC-44 reproducibly suppressed the Muv phenotype in several different SynMuv and Muv genetic backgrounds (Fig. 1b-c, Supplementary Fig. 3). Muv suppression in the *let-60/Ras* gain-of-function mutant specifically suggested that the mode of action occurs cell-autonomously within the VPCs, potentially antagonizing the Ras/MAPK signal cascade or its downstream transcriptional program. However, the initial binary scoring of the Muv phenotype lacked sufficient sensitivity to distinguish weak inhibition of the pathway. To achieve a more sensitive evaluation of Muv suppression, we measured the vulval precursor cell (VPC) induction score following wild isolate feeding [37]. The VPC induction score counts the total number of 6 possible VPCs (P3.p-P8.p) that adopt a primary or secondary vulval fate in an individual. A wild-type animal with 3 induced VPCs (P5.p-P7.p) receives a score of 3. Representative images of *lin-52; lin-8* with two pseudovulvae at the P3.p and P4.p positions demonstrate a VPC induction score of 5 and *let-60* with three psuedovulvae at the P3.p, P4.p, and P8.p positions demonstrate a VPC induction score of 6 (Fig. 4a).The VPC induction score metric allows for a more sensitive readout of suppression of Ras/MAPK overactivation in the population by capturing how many ectopic cell fates were suppressed by the wild bacterial isolate diet.

**Fig. 4.**
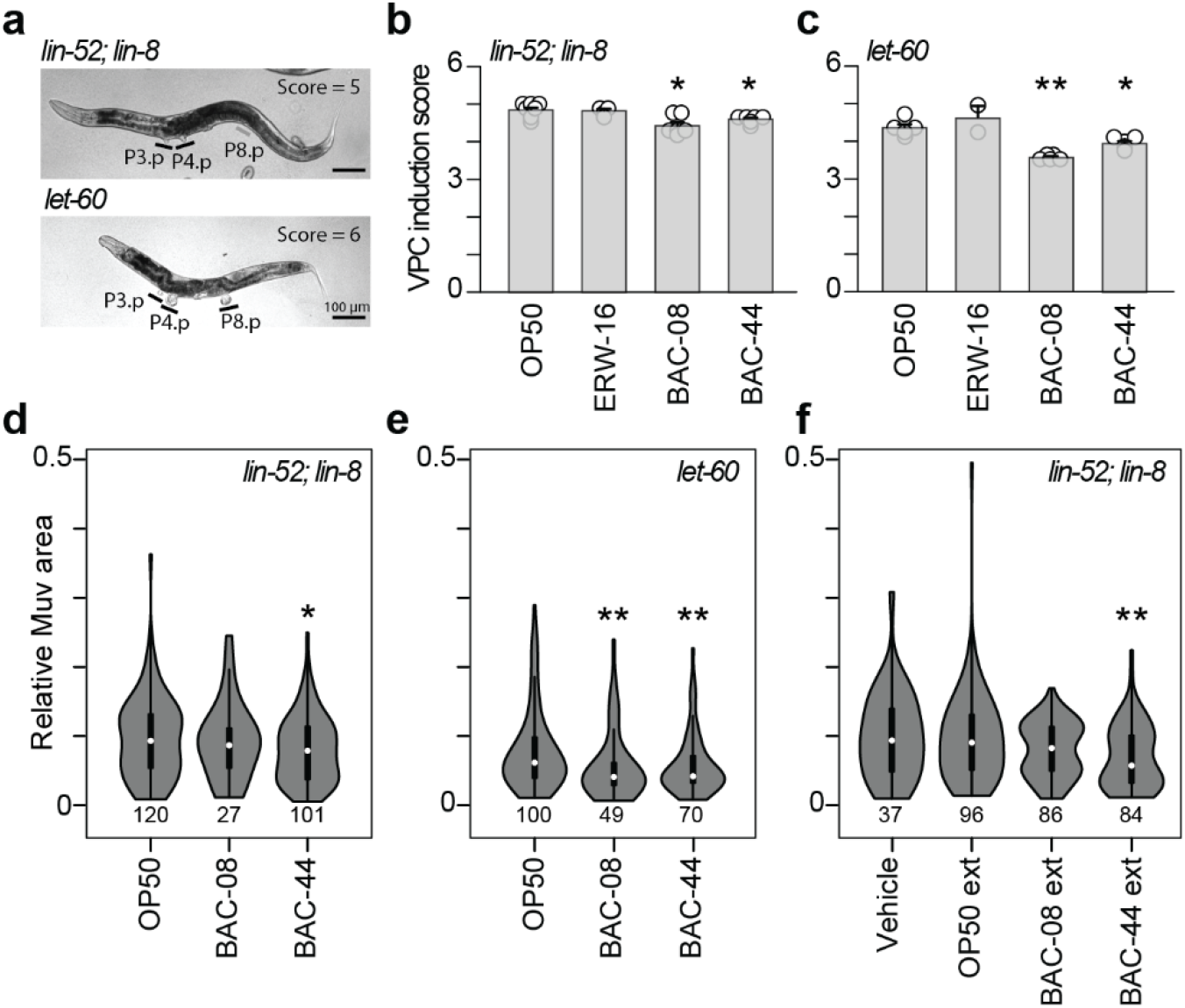
BAC-08 and BAC-44 suppress vulval induction and growth in *C. elegans* multivulva strains. (a) Representative DIC micrographs of live *lin-52; lin-8* (top) and *let-60* (bottom) adult hermaphrodites fed OP50, with induced vulval precursor cells (VPCs) labeled with a black bar and uninduced VPCs labeled with a grey bar. Score indicates the number of induced VPCs in the representative worms shown. Scale bar = 100 µm. b-c) VPC induction scores for *lin-52; lin-8* (b) and *let-60* (c) fed OP50, ERW-16, BAC-08, or BAC-44. Error bars represent mean ± SEM and the individual data points indicate the average for each biological replicate experiment. (d-e) Relative multivulva (Muv) area of individual pseudovulva in *lin-52; lin-8* (d) and *let-60* (e) fed live OP50, BAC-08, or BAC-44. (f) Relative Muv area of individual pseudovulva in *lin-52; lin-8* treated with bacteria cell-free extracts (ext) or vehicle control and fed OP50. Violin plots show the full data distribution, with a white dot indicating the median and black bar the interquartile range. Total number of individual pseudovulva measured (n) for each condition is shown below each violin plot. Statistical significance was determined by a Wilcoxon-Mann-Whitney test with * p < 0.05 and ** p < 0.01 compared to OP50 (b-e) or Vehicle (f) controls.

Using the VPC induction score, we fed *lin-52; lin-8* SynMuv and *let-60* Muv worms OP50, ERW-16, BAC-08, and BAC-44 (Fig. 4a-c). The average of VPC induction scores across 5-6 biological replicate populations revealed that both BAC-08 and BAC-44 significantly suppress VPC induction in both *lin-52; lin-8* and *let-60* worms, as compared to 6-7 biological replicate populations fed OP50 (Fig. 4b-c). We did not observe VPC induction suppression in populations fed ERW-16 assessing 2-4 biological replicate populations (Fig. 4b-c). The decrease in the VPC induction score was driven by a distinct population shift toward fewer ectopic inductions. Specifically, while OP50 and ERW-16 fed populations mostly contained individuals with 2-3 ectopic pseudovulvae, populations fed BAC-08 and BAC-44 exhibited a marked increase in the proportion of animals with only 0-1 ectopic pseudovulvae (Supplementary Fig. 5). Altogether, our results indicate that BAC-08 or BAC-44 diets reproducibly inhibit the frequency of ectopic VPC induction.

In addition to reducing the frequency of VPC induction events, we observed that the ectopic pseudovulva that developed appeared visibly smaller compared to those observed on OP50. To quantify these potential morphological differences independent of overall body size, we measured the area of each ectopic pseudovulva and normalized to the square of the worm’s width at the induction site. Using this metric, we assessed the impact of the live BAC-08 and BAC-44 diet on pseudovulva growth. In the *lin-52; lin-8* SynMuv strain, only a BAC-44 diet significantly reduced the size of ectopic pseudovulva compared to OP50 (Fig. 4e). However, in the *let-60* Muv strain, both BAC-08 and BAC-44 significantly reduced the pseudovulva size compared to OP50 (Fig. 4f). We speculate that the BAC-08 and BAC-44 diet may better counteract hyperactive Ras signaling with regards to pseudovulva growth, whereas their inhibitory effect from BAC-08 may be insufficient to overcome the ectopic LIN-3 overexpression in SynMuv strains if VPC induction was triggered.

To determine if the suppression of the Muv phenotype is mediated by a bioactive secondary metabolite, we performed an intracellular metabolite extraction of OP50, BAC-08, and BAC-44 using 80% methanol. Extracts were dried onto standard OP50 NGM plates to isolate the intracellular chemical effects from the effects from the live bacterial diet. We first evaluated the VPC induction score but observed that the methanol vehicle significantly suppressed induction in the *let-60* Muv strain (Supplementary Fig. 6), indicating that the methanol-induced toxicity or stress masks Ras/MAPK signaling overactivation. We also did not observe suppression of VPC induction in the *lin-52; lin-8* SynMuv strain (Supplementary Fig. 6), suggesting that intracellular metabolites isolated from this extraction method are insufficiently concentrated to inhibit VPC induction events in the developing larvae. Nevertheless, we observed that treatment with the BAC-44 extracts reduced pseudovulva size in *lin-52; lin-8* adults (Fig. 4f). The bacteria cell-free extracts also preserved growth inhibition of the adult worms previously observed in the live bacterial diets, although to a lesser extent (Supplementary Fig. 7), indicating that intracellular metabolites mediate general growth inhibition and not secondary physiological effects like nutritional deficiency. Altogether, our results demonstrate that the WormFood CURE is a viable experimental pipeline to identify wild bacterial isolates that harbor natural products capable of modulating conserved signaling pathways.

Lastly, we were interested in leveraging our bacterial isolate genomic data to characterize the biosynthetic potential of BAC-08 and BAC-44 relative to non-suppressing isolates. We aimed to evaluate whether the phenotypic suppression correlated with distinct biosynthetic gene clusters (BGCs), so we included in our analysis all species-level reference genomes and *E. coli* OP50 in our analysis. Across the 21 bacterial genomes, we identified a total of 235 BGCs, of which 145 were identified within our 12 wild bacterial isolate genomes (Fig. 5a). Within our wild isolates, the clusters comprised 32 terpene, 38 non-ribosomal peptide synthetase (NRPS), 40 ribosomally synthesized and post-translationally modified peptide (RiPP), 7 polyketide synthase (PKS), and 28 other BGCs (Fig. 5a). Compared to the non-suppressing *Bacillus* isolates, the BAC-08 and BAC-44 genomes exhibited notably diverse biosynthetic profiles, harboring 14 and 13 total BGCs, respectively (Fig. 5b).

**Fig. 5.**
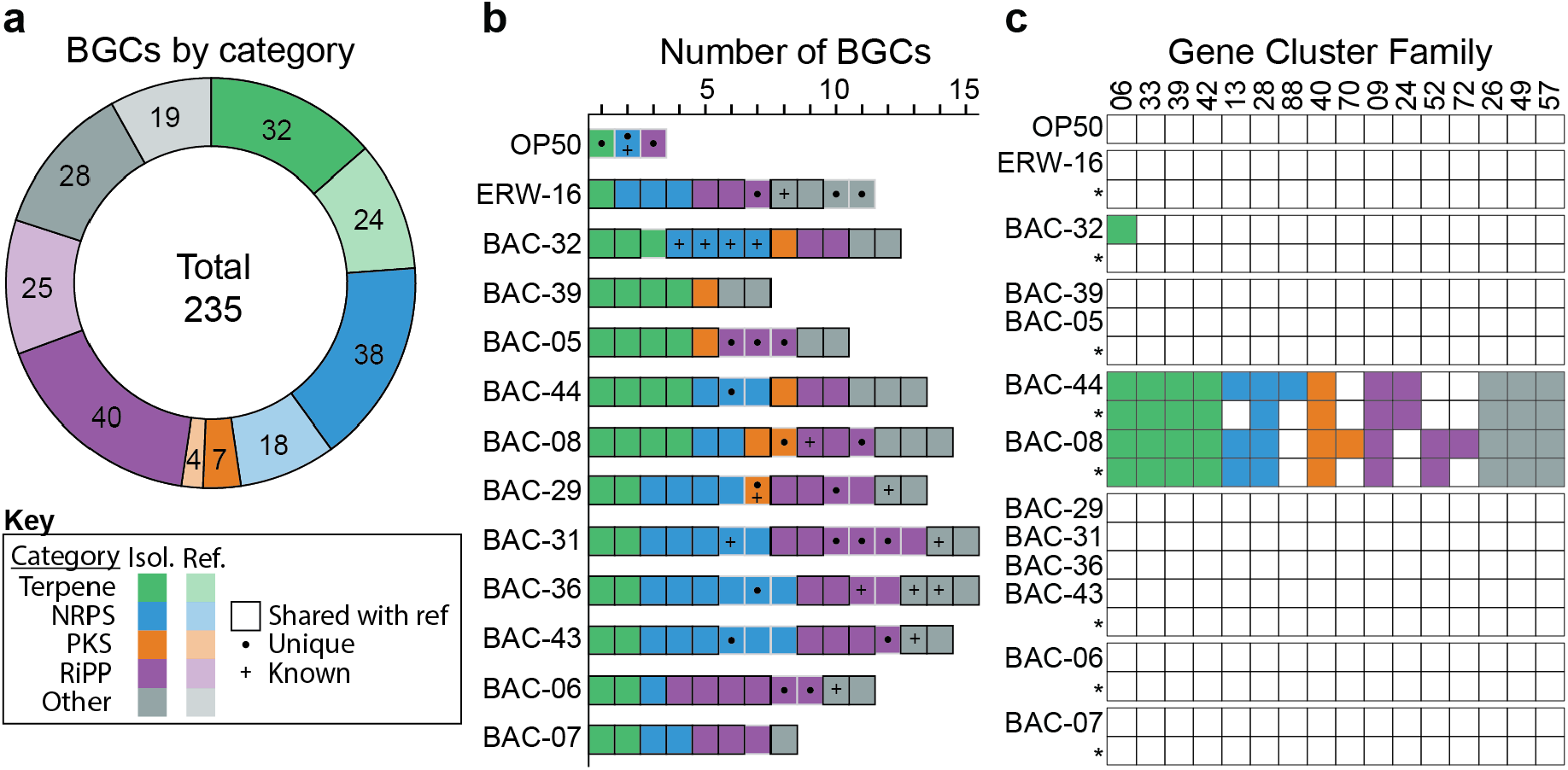
Biosynthetic gene cluster (BGC) diversity within bacterial isolates and reference genomes. (a) Distribution of 235 BGCs across five biosynthetic categories for isolate (Isol.) and reference (ref.) genomes. (b) Number of BGCs per genome colored by biosynthetic category. BGCs belonging to Gene Cluster Families (GCFs) shared with reference genomes are indicated by a black outline. Unique GCFs are marked with a filled circle and BGCs matching known MIBiG entries are marked with a plus. (c) Presence/absence heatmap of GCFs observed in *B. safensis* BAC-08 and *B. altitudinis* BAC-44, with GCF identifiers shown on the x-axis. Colored tiles indicate the BGC category present and white tiles indicate absence. Reference genomes are indicated underneath each isolate with an asterisk (*).

To further investigate genomic divergence, we evaluated which BGCs were unique to BAC-08 and BAC-44 relative to the rest of our wild isolate collection. By grouping BGCs into Gene Cluster Families (GCFs), we identified BGCs specific to these two suppressor isolates (Fig. 5c). Although representing different species, BAC-08 and BAC-44 uniquely share 10 GCFs, comprising terpenes, NRPs, RiPPs, PKSs, and other BGCs (Fig. 5c). The complete distribution of all identified GCFs across the other isolates is provided in Supplementary Fig 8 and Supplemental Table S4. Notably, the BAC-08 and BAC-44 genomes uniquely harbor BGCs predicted to synthesize secondary metabolites that are likely methanol soluble and often implicated in modulating eukaryotic signaling pathways [38,39]. These genomic signatures demonstrate that BAC-08 and BAC-44 possess unique biosynthetic capabilities compared to non-suppressing *Bacillus* isolates, providing a clear genetic basis for further investigation into their specialized metabolic profiles moving forward.

## Discussion

Our study establishes the WormFood CURE as an accessible and scalable platform for the discovery of bacterial secondary metabolites that modulate conserved eukaryotic signaling pathways. By leveraging the Multiullva (Muv) *C. elegans* phenotype, we identified two wild *Bacillus* isolates, BAC-08 and BAC-44, which reproducibly suppress ectopic vulval inductions and pseudovulva growth. While standard high-throughput screens rely on pre-extracted chemical libraries, the WormFood CURE model enables the screen of live environmental bacterial isolates for activities-of-interest. Our approach preserves the ecological context and interactions of the *C. elegans* species’ natural microbiome while simultaneously providing a more direct readout of the active biosynthetic potential of each wild isolate’s genome.

Our study outlined two general challenges to overcome to effectively establish a *C. elegans* dietary screen aimed towards identifying potential bioactive secondary metabolites. First, we demonstrated that a counter screen to eliminate toxic isolates is a necessary step to remove virulent or inedible isolates. Second, we demonstrated that careful consideration is required to ensure that indirect physiological effects do not produce false positives. For evaluating Muv phenotype suppression, larval developmental delay created initial difficulties during the WormFood CURE activities because of course logistics and student difficulty discerning scorable developmental stages. Such difficulties were easily corrected in the lab. Altogether, our pilot screen demonstrates that careful consideration of potential confounding physiological variables is essential in the design of an effective adaptation of the WormFood CURE.

Nutritional deficiency and the general worm starvation response posed a unique challenge for our pilot WormFood Muv suppressor screen. Specifically, L2 starvation antagonizes VPC induction [40,41]. In our study, several lines of evidence suggested that the Muv suppression effects on BAC-08 and BAC-44 were not due to the starvation response: (1) genomic profiling confirmed that both isolates possessed complete metabolic pathways for essential amino acids and key nutrients on par with non-suppressor *Bacillus* strains and OP50 *E. coli*, (2) the reduction in body size was strictly proportional on live bacteria and on OP50 with BAC-08 or BAC-44 intracellular metabolite extracts, and (3) larval developmental delay on both isolates was not as severe as other, non-suppressing *Bacillus* isolates. Altogether, our data indicate that at least BAC-44 produces an intracellular metabolite that mediates both Muv and a general growth suppression, which is an effect in line with an antagonist of the Ras/MAPK growth factor pathway.

The recovery of bioactive compounds from environmental sources requires extraction methods that preserve molecular integrity without introducing host toxicity. Methanol is a widely utilized medium for the recovery of secondary metabolites [42,43]. Unfortunately, our results demonstrated that methanol alone significantly suppressed VPC induction in the *let-60* Muv strain. Our findings suggest that methanol stress or toxicity effectively masks the overactivation of the Ras/MAPK pathway. However, while the methanol vehicle influenced the frequency of induction events in *let-60*, we observed that BAC-44 extracts specifically and significantly reduced pseudovulva size in *lin-52; lin-8* adults. Moreover, identification of multiple shared biosynthetic gene cluster families in BAC-08 and BAC-44 compared to all other non-suppressive *Bacillus* isolates indicates that Muv suppression is driven by a stable, methanol-soluble bacterial product. Altogether, our results provide a clear chemical starting point for future bioassay-guided fractionation and structural characterization.

Thousands of bacteria exist in a single gram of soil, potentially containing another thousand or more uncharacterized secondary metabolites [44,45]. A single microbe harbors the potential to produce as many as 50 secondary metabolites [46]. Method standardization within the field caused a trend towards the isolation of similar species and metabolite recoveries, slowing the recovery rate of novel compounds [47]. Moreover, the more recent employment of molecular and *in silico* methods have revealed that a significant portion of a microorganisms biosynthetic gene clusters are not expressed under many standard laboratory conditions, overlooking compound potential in even known and cultured bacterial species [48,49]. Ecologically, *C. elegans* does not interact with its microbial surroundings passively; but navigates a chemical network in which bacteria are not only deploying metabolites as defense to encroachment and communication with other bacteria, but also as a defense to direct predation [50]. While our study intended to isolate bacteria from known *C. elegans* habitats, the pilot WormFood CURE occurred during the winter, so the students selected more easily attainable environmental samples like ice and snow. Altogether, our results indicate that the environmental sample source is forgiving for identifying isolates-of-interest. Ultimately, while our study was not limited to the natural *C. elegans* microbiome, exploring the broader ecological *C. elegans* niche may yield a richer reservoir of potential bioactive compounds and novel bacterial-nematode genetic interactions.

With over 80% of the C. elegans proteome having homology to human genes [51,52], the WormFood CURE serves as a simple and scalable platform for revealing host-microbiome interactions relevant to human health Previous research on bacterially supplied B12, nitric oxide, folate, and methylglyoxal has functionally solidified the use of *C. elegans* to screen for bacterial products that modulate signaling pathways central to human disease [27,51,53,54]. Our findings expand the precedent by establishing a biological and methodological framework for identifying environmental bacterial isolates in Ras/MAPK-suppressor activity. By integrating bacterial metabolic profiling with phenotypic outcome, we outlined a productive strategy for prioritizing candidate natural products with inhibitory potential. Ultimately, the WormFood CURE demonstrates that student-driven screening effectively bridges the gap between environmental microbiology and the discovery of novel compounds for the targeted modulation of oncogenic signaling. In the future, we expect a similar strategy may be adapted for other conserved signaling pathways will enable the systematic exploration and screening of the environmental metabolome for novel chemical modulators of eukaryotic development.

## Methods

### Worm strains

All worm strains were maintained using standard methods at 20°C on Nematode Growth Medium (NGM) agarose plates with *Escherichia coli* OP50, unless otherwise noted. Some strains were provided by the Caenorhabditis Genetics Center (CGC), which is funded by the NIH Office of Research Infrastructure Programs (P40 OD010440). We used the following strains for this study: Wild-type (WT) N2 (Bristol), SS1408 *lin-8(n2731); lin-52(bn151)*, MT2124 *let-60(n1046)* IV, MT309 *lin-15AB(n309)* X, CB1322 *lin-8(n111)* II; *lin-9(n112)* III, MT1006 *lin-1(n431)* IV.

### Bacterial strain isolation and identification

Each participant in the WormFood CURE isolated one bacterial colony from an environmental sample (snow, ice, soil, or other) collected in a sterile 15 mL conical tube from the local area. Samples from human or animal sources were strictly prohibited. We diluted samples 1:1 with LB media, incubated them at 37°C for 2 hours, and then plated 25 µL of broth onto one LB agar plate and one 5% sheep blood agar plate. Each isolate was characterized by visual colony morphology and gram staining. Bacterial isolates were taxonomically identified by PCR amplification of the 16S rRNA gene using universal primers 27F (5’-AGAGTTTGATMTGGCTCAG-3’) and 1492R (5’-TACGGYTACCTTGTTACGACTT-3’), followed by Sanger sequencing of the amplicons by Eton Bioscience (San Diego, CA).

Forward and reverse Sanger sequencing reads of 16S rRNA gene amplicons were assembled by overlapping sequences anchored to conserved internal regions (C2: 5′-CTACGGGAGGCAGCAG-3′; C3: 5′-GTGCCAGCAGCCGCGGTAA-3′; C4: 5′-AACAGGATTAGATACCCTGGTAGTC-3′). High-quality sequence upstream of the C2 region was retained from forward reads, while high-quality sequence downstream of the C4 region was retained from reverse reads. The 5’ and 3’ ends of each assembled sequence were trimmed based on quality alignment to the conserved (C) regions in the *Escherichia coli* 16S gene using Clustal Omega [55]. The assembled sequences were used for BLAST-based taxonomic identification of each bacterial isolate [56]. All isolate information, including sample source, colony morphology, taxonomic identification, and the variable regions (V1–V9) represented in each final sequence are listed in Supplementary Table 1. The final assembled sequence for each isolate is also provided in Supplementary Information. The phylogenetic tree of the wild isolate 16S rRNA V3-V4 region sequences was constructed using Clustal Omega and visualized with the Interactive Tree of Life (iTOL) [57].

### WormFood CURE phenotype screening and vulval induction scoring

NGM agarose plates were seeded with wild isolates grown in LB media for 24 – 72 hours at 30°C. For the WormFood CURE phenotype screening assays, we performed a timed egg-laying experiment with 5 gravid adults for four hours, which resulted in *E. coli* OP50 contamination. For follow-up experiments in the lab, we performed the spot bleaching synchronization procedure on 10-20 gravid adults to remove residual OP50. For the toxicity assay, we counted the number of live adult worms per plate following a 4-day incubation at 20°C. For multivulval (Muv) phenotype scoring, we performed timed egg-laying experiments for the WormFood CURE and spot bleaching synchronization for follow-up experiments. Progeny were incubated at 20°C, then scored for the absence or presence of pseudovulva.

Vulval induction was scored young adult hermaphrodites using a dissecting microscope by examining the number of pseudovulva, which correspond to the ectopic induction of vulval precursor cells (VPCs). A vulval induction score of 3 represents a wild-type worm, indicating the normal induction of P5.p-P5.p VPCs. Scores greater than 3 indicate a Muv phenotype where tertiary VPCs P3.p, P4.p, or P8.p were ectopically induced to adopt vulval fates [37]. For each genotype and treatment, the vulval induction scores for all individuals on a single plate were averaged to represent one biological replicate.

### Microscopy and image analysis of wild bacterial isolates and *Caenorhabditis elegans* strains

Gram staining of wild bacterial isolates was performed using BD BBL Gram Stain Kit (BD Diagnostic Systems) and slides were imaged using a Leica DM750 and 100x Plan oil objective. Bacterial cell diameter and length measurements were measured using Image J [58]. Bacterial cell volume was calculated based on cell length and width measurements using the equation for a cylinder with hemispherical ends. All measurements were performed on at least 20 individual cells per isolate.

Light micrographs of live adult worms fed wild bacterial isolates were acquired using a Zeiss Axioskop Plus 2 microscope using a Nikon Plan Neofluar 10x air objective with a PCO Panda 4.2 sCMOS camera (Excelitas) using μManager software [59]. Wild-type adults were mounted on 2-5% agarose pads and immobilized using 20 mM levamisole in 1x S-basal under a cover slip and imaged. To prevent cuticle rupture and preserve the integrity of the pseudovulvae, Muv adults were immobilized using 20 mM levamisole in 1x S-basal within a watch glass.

To assess the physiological impact of bacterial diets on worm growth, total body length was measured from head to tail along the midline of the animal using ImageJ. The body width was measured at the vulval region, and the width-to-length ratio was calculated to assess proportional size reduction. To quantify pseudovulva morphological changes independent of variations in overall body size, we developed a relative area index for ectopic pseudovulva. For each adult, the two-dimensional surface of every ventral protrusion was manually traced using ImageJ to acquire the Muv area. Each Muv area was normalized to the square of the width of the worm at each induction site.

### Bacterial isolate genomic sequencing, assembly, and analysis

Genomic DNA extraction, library preparation, and genomic sequencing was performed by SeqCenter (Pittsburgh, PA) according to their Illumina Whole Genome Sequencing service. Briefly, genomic DNA was extracted using the ZymoBIOMICS DNA Miniprep kit (Zymo Research), following manufacturer’s instructions. Illumina sequencing libraries were prepared using the Illumina DNA Prep Kit (Illumina) with a target insert size of 320 bp. Libraries were sequenced on an Illumina NovaSeq 6000 sequencer, producing 2 × 151 bp paired-end reads.

Raw sequencing reads we *de novo* assembled using SPAdes v3.15.5 [60]. The quality and completeness of the resulting assemblies were assessed using CheckM2 v1.1.0 [61], and the Average Nucleotide Identity (ANI) comparing isolates to reference genomes was calculated using skani [62]. Taxonomic classification of bacterial isolate genomes was performed using the Genome Taxonomy Database Toolkit (GTDB-Tk) v2.6.1 using the r226 database [63]. To ensure consistency in downstream genome profiling, functional annotation of both isolate and reference genomes was conducted using Bakta v1.11.4 with the full database v6.0 [64]. For metabolic pathway reconstruction, protein sequences were screened against the Kofam database using KofamScan [65], and the resulting KO assignments were processed with KEGGDecoder [66] to visualize pathway completeness. To supplement these analyses, we used KEGGMapper [67] to investigate specific metabolic networks not covered by KEGGDecoder. The highest completion score for the following KEGG modules from KEGGMapper were included in our analysis: Heme biosynthesis (M00926 / M00121), B3 NAD biosynthesis (M00115 / M00912), B5 pantothenate biosynthesis (M00119 / M00913), B6 pyridoxal phosphate biosynthesis (M00916 / M00124), B7 biotin biosynthesis (M00123 / M00950 / M00573 / M00577), and B9 tetrahydrofolate biosynthesis (M00126 / M00840 / M00841). We predicted Biosynthetic gene clusters (BGCs) using antiSMASH v8.0.4 [68] on the Bakta annotation files with flags –fullhmmer, --asf, --cb-general, --cb-subclusters, and --cb- knownclusters enabled. Predicted BGCs were grouped into Gene Cluster Families (GCFs) alongside similar known BGCs from MIBig 4.0 [69] using the BiG-SCAPE v2.0.0 [70] cluster command with -m 4.0 --mix, --classify category, --include-singletons, and the default --gcf- cutoffs 0.3 flags enabled. All bacterial genome assembly statistics are provided in Supplemental Table S2, and genome annotation statistics are provided in Supplemental Table S3. All BiG-SCAPE 2.0 results are provided in Supplemental Table S4.

### Intracellular bacterial metabolite extraction

To isolate secondary metabolites, we harvested 100 mL overnight bacterial cultures, acquiring a high-density microbial culture of each isolate to an OD600 of approximately 1.7. Bacterial pellets were flash-frozen on dry ice prior to extraction. Bacterial pellets were resuspended in chilled 80% methanol extraction mixture and sonicated at 50% amplitude using a Qsonica Q125 Sonicator for 5 minutes (30 s on / 30 s off cycles). Following sonication, the extraction mixture was centrifuged at 4°C to remove cellular debris. The extraction process was repeated three times in total, recovering a total volume of 5 mL of clarified intracellular metabolite extracts per isolate. We overlaid 350 µL of the prepared extract on standard OP50 NGM plates, allowing the extracts to dry completely before the addition of synchronized *C. elegans* embryos via bleach synchronization for experimental analyses.

## Acknowledgements

We thank the students of the Spring 2021 WormFood CURE cohort at Michigan Technological University for their essential contributions to environmental bacteria strain isolations, characterization, and identification and *C. elegans* phenotype scoring.

## Author contributions

Conceptualization: EW, PDG. Computational analysis: EW, PDG. Data generation: EW, EM, RP, CH, MG, AB, PDG, PDG. Data curation: PDG. Funding acquisition: PDG. Animal maintenance: EW, PDG. Supervision: PDG. Visualization: EW, PDG. Writing (original draft): EW, PDG. Writing (review and editing): EW, EM, RP, CH, MG, AB, PDG.

## Data availability

All data supporting the findings of this study are available within the paper and its Supplementary Information. All *C. elegans* strains and wild bacterial isolates are available upon reques. Genomic data were deposited at NCBI under BioProject accession PRJNA1473339. We used the following references for genomic comparisons: *E. coli* OP50 (GCA_004355015.1), *E. aphidicola* (GCA_037149315.1), *B. glycinifermentans* (GCA_004103615.1), *P. megaterium* (GCA_006094495.1), *B. altitudinis* (GCA_001191605.1), *B. safensis* (GCA_039619585.1), *B. cereus* (GCA_006094295.1), *B. toyonensis* (GCA_016605985.1), and *B. paranthracis* (GCA_024296885.1)

