## Supplementary Information for "The WormFood CURE: Screening for bioactive metabolites that antagonize the *Caenorhabditis elegans* Ras signaling pathway"

### Supplementary Figures

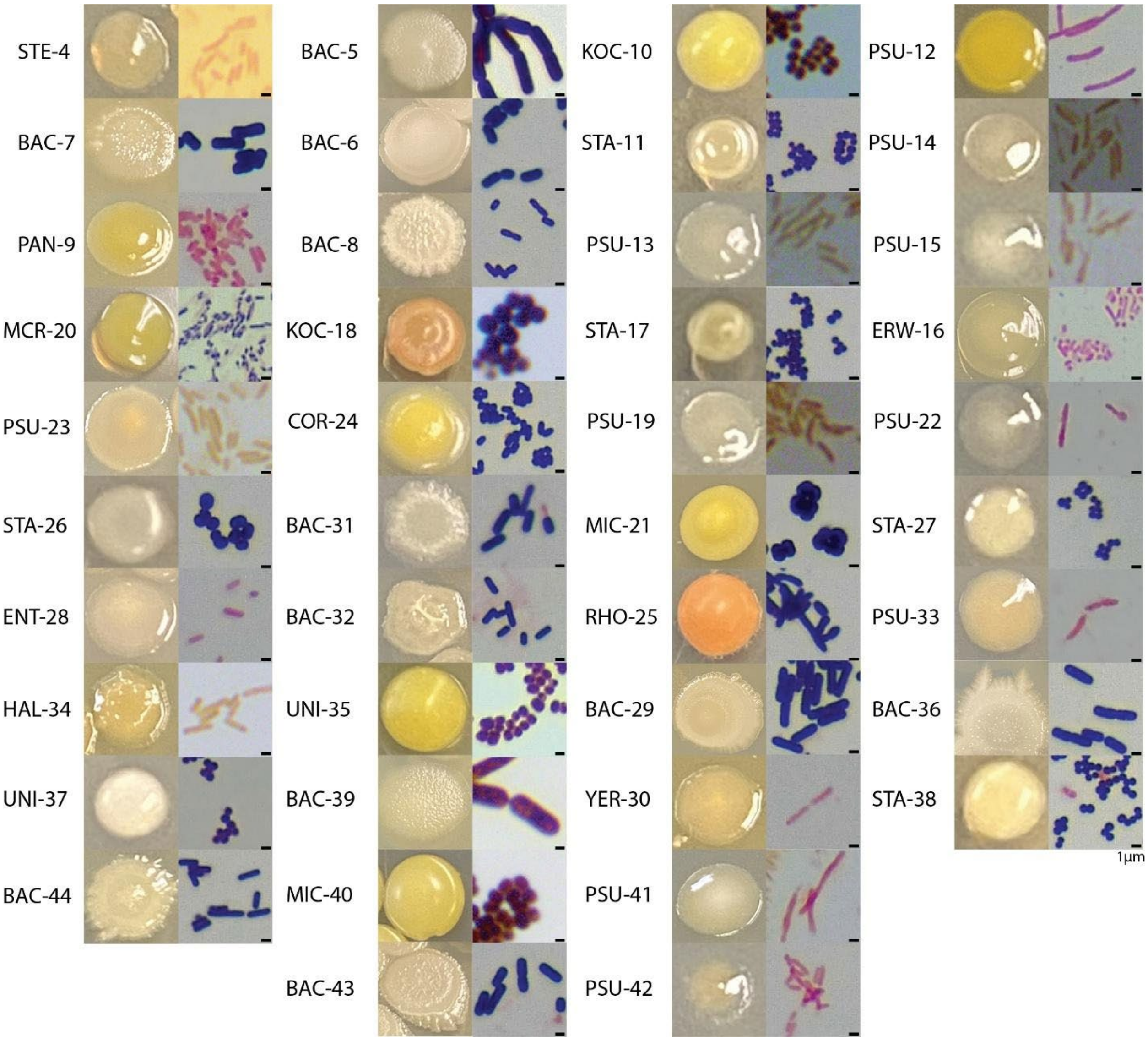


**Supplementary Fig. 1. Morphological and microscopic characterization of bacterial isolates.** The left of each panel displays representative images of wild bacterial isolate colonies on LB agar plates. The right of each panel displays the gram staining results for each wild isolate. Scale bar for microscopy images represents 1 μm

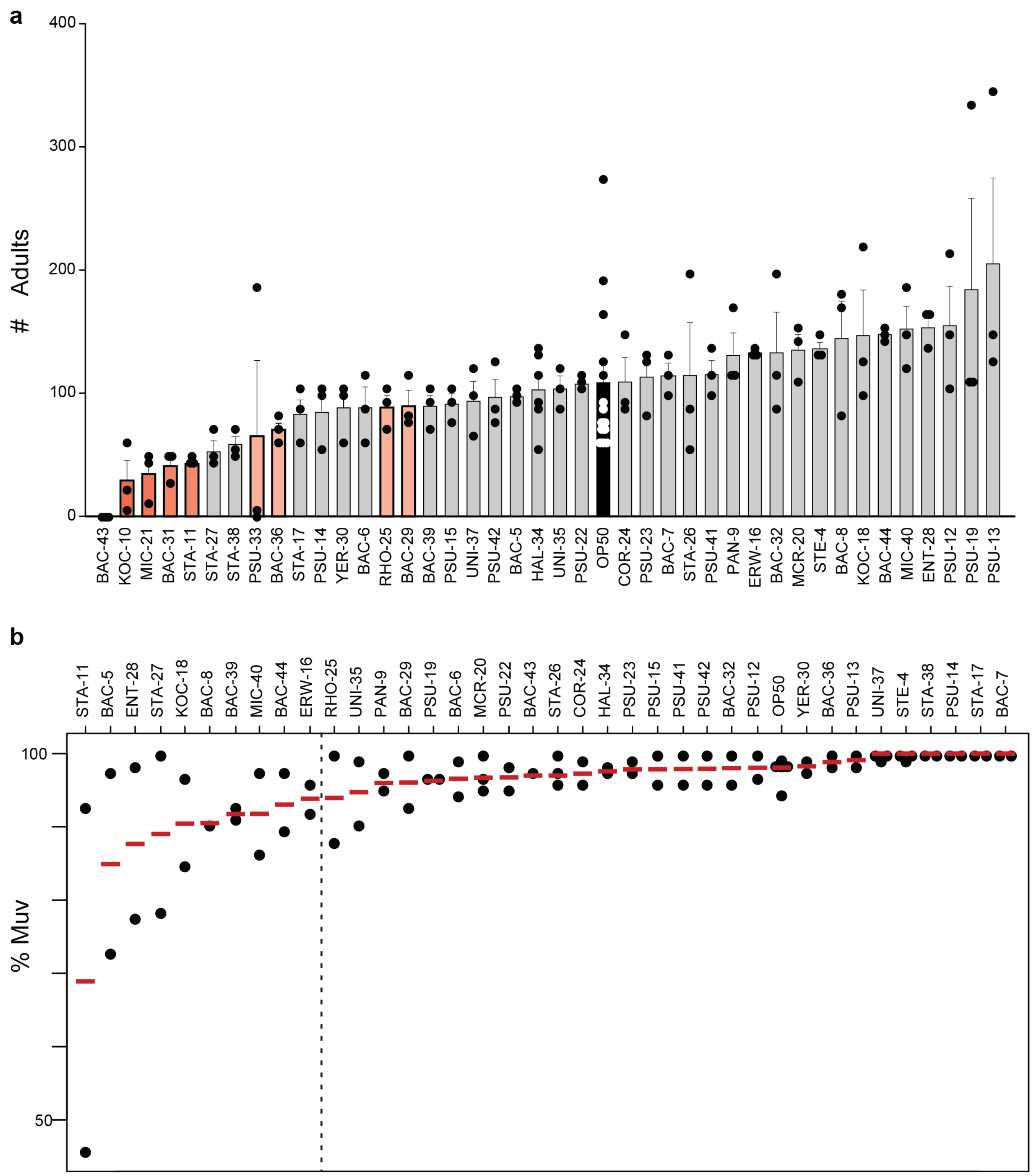


**Supplementary Fig. 2. WormFood CURE *C. elegans* phenotype screening results.** (a) Quantification of adult population size following a timed limited egg-lay and 4-day incubation at 20°C on the indicated wild bacterial isolate. Bars represent the mean number of adults and error bars indicate the standard error of the mean, with individual replicates shown as black or white dots. The standard laboratory food source, *E. coli* OP50, is highlighted in black for comparison.

Isolates that were specifically followed up in the research lab are highlighted in red. (b) Percentage of Muv worms observed in *lin-52; lin-8* mutants fed the indicated isolate. Dashed vertical line separates isolates that were evaluated further in the research lab (left) from candidates showing no effect (right). Red horizontal bars indicate the median and the black points are individual data points for each biological replicate.

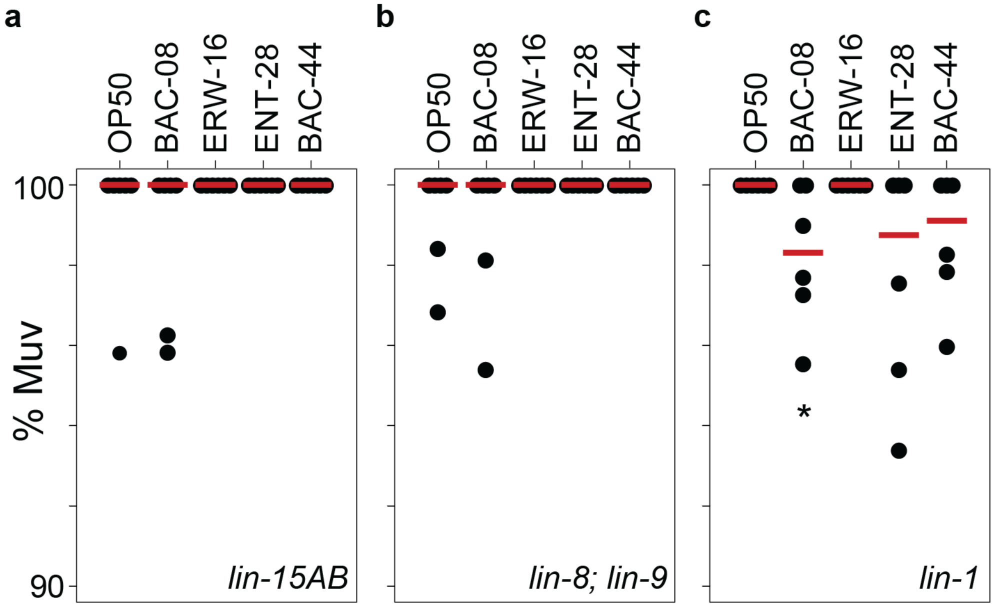


**Supplementary Fig. 3. Impact of wild bacterial isolate diet on Muv phenotype across different *C. elegans* synthetic multivulva (SynMuv) and multivulva (Muv) strains.** Percentage of Muv worms observed in (a) *lin-15AB,* (b) *lin-8; lin-9,* and (c) *lin-1* fed the indicated isolate. Red horizontal bars indicate the median and the black points are individual data points for each biological replicate. Statistical significance was determined by a Wilcoxon-Mann-Whitney test with * $p < 0.05$ compared to OP50 control.

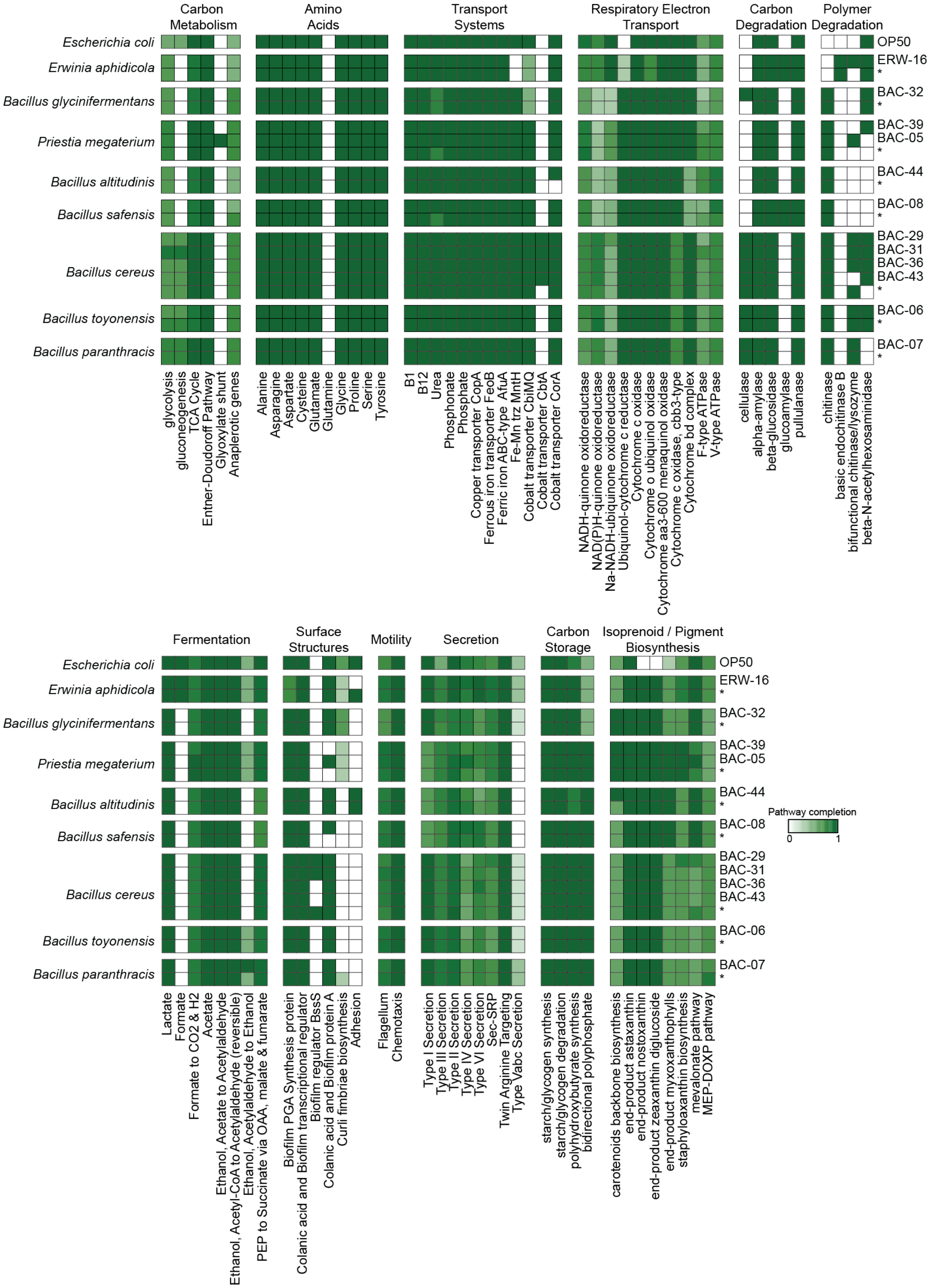


**Supplementary Fig. 4. Expanded pathway reconstruction of bacterial isolates.** Heatmap analysis showing the predicted biosynthetic pathway completion (gradient from white = 0 to dark green = 1) for carbon metabolism, additional amino acids, transport systems, respiratory electron transport, carbon degradation, polymer degradation, fermentation, surface structures, motility, secretion, carbon storage, and isoprenoid / pigment biosynthesis from KEGGdecoder analysis of the genome assembly of each wild bacterial isolate. An asterisk (*) indicate analysis of the reference genome for the indicated bacterial species identified by GTDB-Tk.

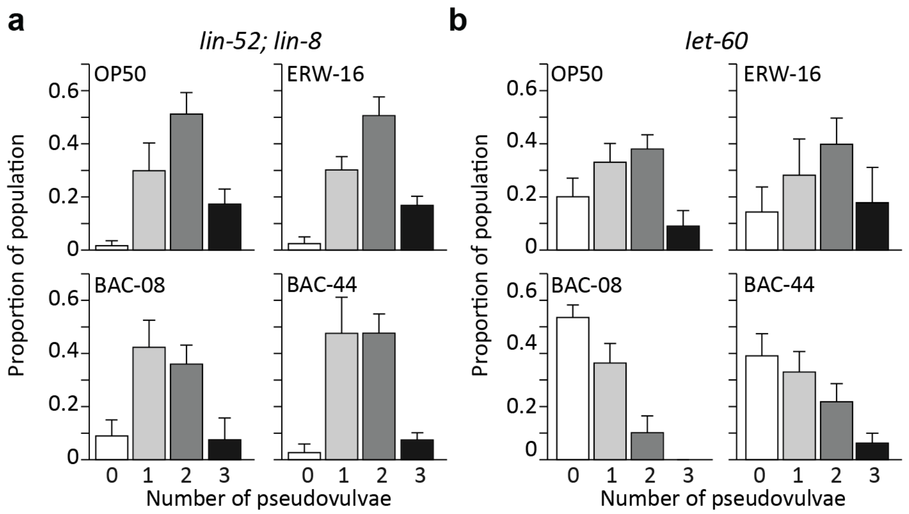


**Supplementary Fig. 5. Multivulva (Muv) phenotype expressivity across different bacterial diets.** Histograms representing the number of pseudovulvae induced on average across biological replicate populations in (a) *lin-52; lin-8* and (b) *let-60* genetic backgrounds. Worms were fed three selected wild bacterial isolates, ERW-16, BAC-08, and BAC-44, and compared to *E.coli* OP50. Data represent the average proportion of the population displaying zero (white), one (light gray), two (dark gray), or three (black) pseudovulvae across biological replicates. Error bars indicate the standard deviation.

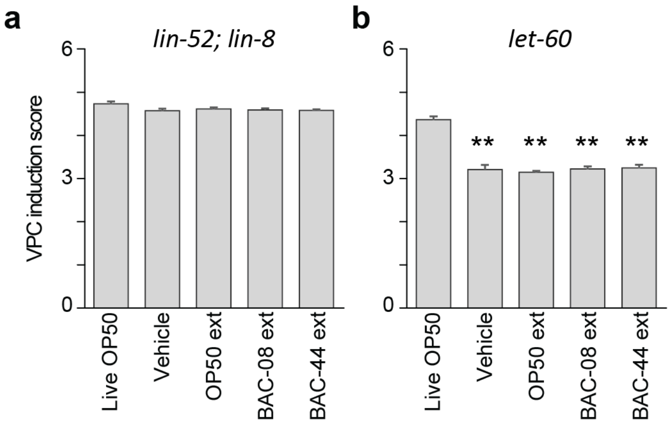


**Supplementary Fig. 6. Methanol obscures Vulval Precursor Cell (VPC) induction.** VPC induction scores for (a) *lin-52; lin-8* and (b) *let-60* fed live OP50 alone, live OP50 with 80% methanol (vehicle), or live OP50 with cell-free methanol extracts (ext) from OP50, BAC-08, or BAC-44, Bars represent the mean of the average VPC induction scores across biological replicate, with error bars indicating the standard error of the mean. Statistical significance was determined by a Wilcoxon-Mann-Whitney test with ** $p < 0.01$ compared to live OP50.

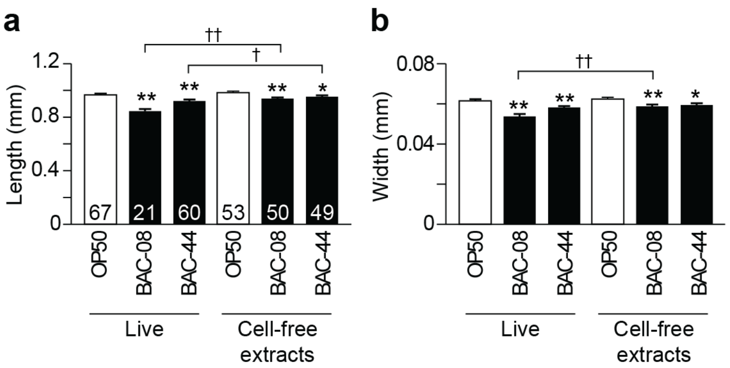


**Supplementary Fig. 7. Bacterial cell free extracts preserve BAC-08 and BAC-44 worm growth inhibition.** (a-b) Bar graphs of (a) body length and (b) body width measurements of *lin-52; lin-8* adult hermaphrodites fed live BAC-08 or BAC-44 (left) or treated with bacteria cell-free extracts and fed OP50 (right). Total number of individuals measured (n) for each condition is shown within each length bar. Error bars represent the standard error of the mean (SEM). Statistical significance was determined by a student's T-test with * $p < 0.05$ and ** $p < 0.01$ compared to live OP50 or OP50 extract controls and † $p < 0.05$ and †† $p < 0.01$ comparing cell-free extracts to live bacteria for BAC-08 or BAC-44.

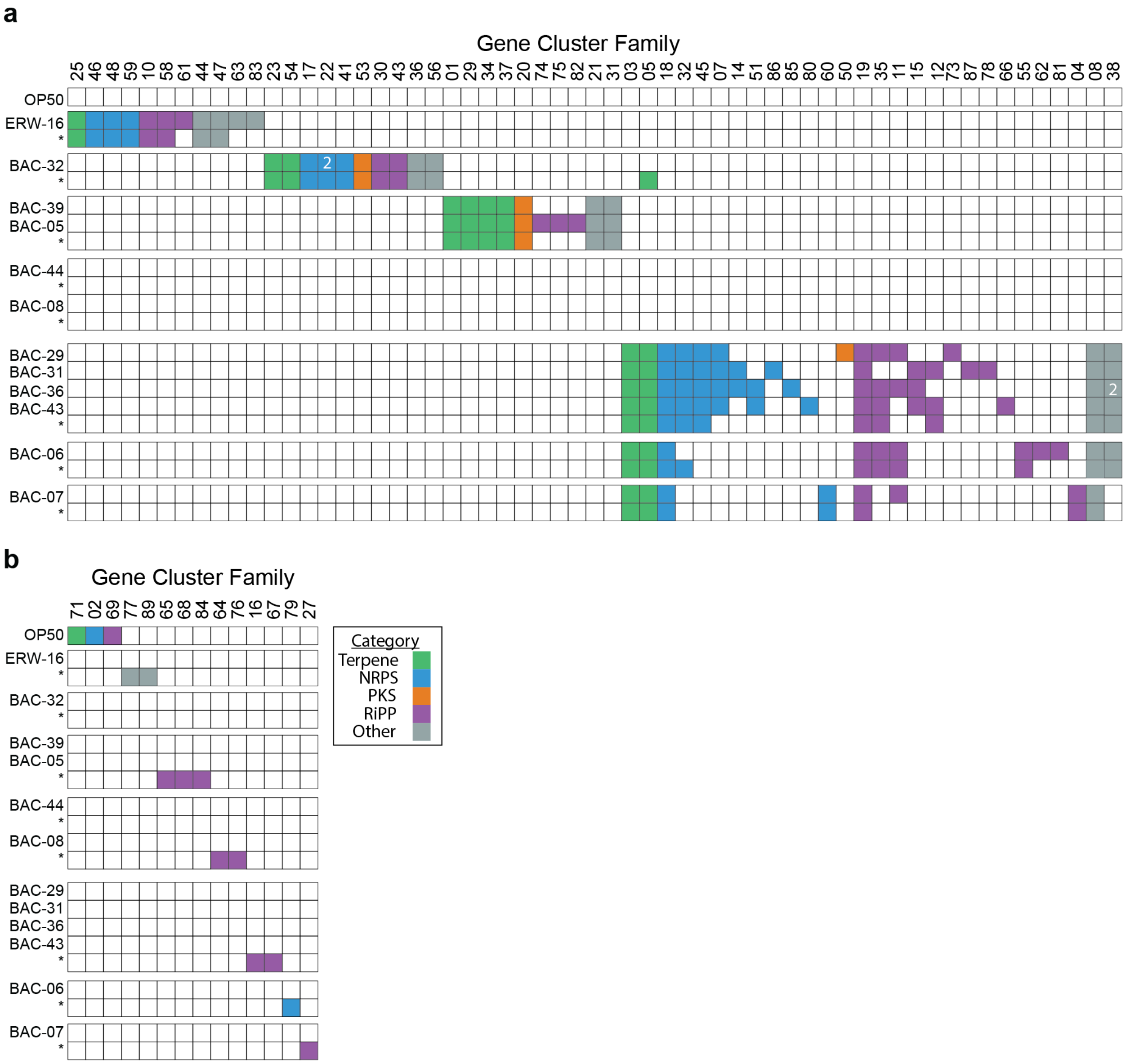


**Supplementary Fig. 8. Extended distribution of Biosynthetic Gene Cluster families across bacterial isolates.** Presence/absence heatmap of GCFs observed in all other wild isolate strains (a) or exclusive to reference genomes (b), with GCF identifiers shown on the x-axis. Colored tiles indicate the BGC category present and white tiles indicate absence. Reference genomes are indicated underneath each isolate with an asterisk (*).

**Supplementary Table S1. Source, characteristics, and 16S rRNA gene sequence results for wild bacterial isolates**

| Isolate ID | Sample Type | Colony Morphology | Gram Stain | 16S Consensus Sequence (length bp) | Consensus ID (Percent ID) |
|---|---|---|---|---|---|
| **STE-04** | Soil | tan, round, wet, medium | GNR | V1-V9 (1419) | *Stenotrophomonas* (99.93%) |
| **BAC-05** | Snow | grey, flat, non-hemolytic, round | GPR | V2-V9 (1354) | *Bacillus* (99.04%) |
| **BAC-06** | Other/ Kombucha | Off-white, round, large, dry | GPR | V2-V4 (709) | *Bacillus* (96.88%) |
| **BAC-07** | Snow | med, grey-white, flat, round, smooth | GPR | V3-V8 (1078) | *Bacillus* (100%) |
| **BAC-08** | Soil | Large, brown, round, dry, raised | GPR | V2-V8 (1260) | *Bacillus* (100%) |
| **PAN-09** | Snow | grayish-white, round, wet, medium | GNR | V2-V6 (1018) | *Pantoea* (99.51%) |
| **KOC-10** | Brine | bright yellow, round, smooth | GPC | V3-V8 (994) | *Kocuria* (98.20%) |
| **STA-11** | Snow | white, round, wet, small | GPC | V3-V7 (980) | *Staphylococcus* (99.29%) |
| **PSU-12** | Snow | yellow, round, wet, medium | GNR | V3-V8 (1107) | *Pseudomonas* (99.73%) |
| **PSU-13** | Soil | gray, round, smooth, medium, flat | GNR | V3-V8 (1032) | *Pseudomonas* (95.27%) |
| **PSU-14** | Snow | moist, moderate, yellow-white | GNR | V2-V8 (1268) | *Pseudomonas* (100%) |
| **PSU-15** | Snow | yellow, flat, sticky | GNR | V2-V8 (1269) | *Pseudomonas* (100%) |
| **ERW-16** | Fruit | small, grey, round, wet | GNR | V1-V9 (1390) | *Erwinia* (99.28%) |
| **STA-17** | Other/ Simple Syrup | medium, sticky, bright yellow, wet | GPC | V2-V8 (1262) | *Staphylococcus* (100%) |
| **KOC-18** | Snow | pink, round, convex, wet | GPC | V2-V8 (1239) | *Kocuria* (100%) |
| **PSU-19** | Soil | Grey-black,wet, beta homolysis,smooth,round | GNR | V2-V8 (1252) | *Pseudomonas* (99.84%) |

| | | | | | |
|---|---|---|---|---|---|
| **MCR-20** | Soil | wet, round, bright yellow. small | GPR | V3-V8 (1149) | *Microbacterium* (99.91%) |
| **MIC-21** | Snow | Black, round, small, dry, convex | GPC | V3-V8 (1096) | *Micrococcus* (99.91%) |
| **PSU-22** | Snow | flat, round, wet, light yellow | GNR | V2-V9 (1335) | *Pseudomonas* (100%) |
| **PSU-23** | Water | yellow,round,wet/sticky,medium | GNR | V3-V9 (1084) | *Pseudomonas* (98.54) |
| **COR-24** | Compost Soil | off-white, round, small, wet | GPR | V2-V8 (1229) | *Corynebacterium* (93.87%) |
| **RHO-25** | Compost Soil | off-white, round, small, dry | GPR | V2-V8 (1233) | *Rhodococcus* (99.92%) |
| **STA-26** | Snow | off-white, round, small, sticky, flat | GPC | V2-V8 (1322) | *Staphylococcus* (99.85%) |
| **STA-27** | Snow | off-white, round, small, wet | GPC | V3-V5 (649) | *Staphylococcus* (99.69%) |
| **ENT-28** | Snow | small, yellow, wet | GNR | V1-V9 (1399) | *Enterobacter* (99.57%) |
| **BAC-29** | Water | large, round, long, flat | GPR | - | - |
| **YER-30** | Fruit | yellow, round, small, raised | GNR | V1-V9 (1426) | *Rahnella* (99.09%) |
| **BAC-31** | Snow | Large, round, flat, rough, off white | GPR | - | - |
| **BAC-32** | Snow | Large, brown, mucoid, flat, round, entire | GPR | V2-V8 (1277) | *Bacillus* (100%) |
| **PSU-33** | Water | small, wet, round, mucus | GNR | V2-V9 (1355) | *Pseudomonas* (99.63%) |
| **HAL-34** | Snow | off-white, medium, irregular, dry, raised | GNR | V2-V9 (1352) | *Halomonas* (99.63%) |
| **UNI-35** | Snow | off white/yellow, medium, mucoid, umbonate, round, entire | GPC | V2-V3 (709) | *Bacillus* (96.92%) |
| **BAC-36** | Soil | off-white, medium, dry, flat, round, serrate | GPR | V3-V7 (1000) | *Bacillus* (97.40%) |
| **UNI-37** | Water | off-white, wet, round, medium | GPC | V2-V8 (1268) | *Fictibacillus* (100%) |

| | | | | | |
|---|---|---|---|---|---|
| **STA-38** | Snow | cream, med, smooth | GPC | V3-V8 (1090) | *Staphylococcus* (99.27%) |
| **BAC-39** | Ice | large, grey, flat/raised, round | GPR | V3-V5 (707) | *Bacillus* (86.33%) |
| **MIC-40** | Soil | Mid-sized, yellow, round, wet | GPC | V3-V7 (873) | *Micrococcus* (96.71%) |
| **PSU-41** | Snow | medium, wet, round , translucent | GNR | V2-V9 (1357) | *Pseudomonas* (99.34%) |
| **PSU-42** | Compost Soil | small, yellow, "fried egg" | GNR | V2-V8 (1257) | *Pseudomonas* (98.97%) |
| **BAC-43** | Snow | small, off white, round, dry | GPR | V2-V9 (1360) | *Bacillus* (100%) |
| **BAC-44** | Water | small, brown, raised, wet | GPR | V2-V9 (1358) | *Bacillus* (100%) |

**Note:** GNR, Gram-negative rod. GPR, Gram-positive rod. GPC, Gram positive cocci. bp, base pairs. V1-V9 denote the variable regions of the 16S rRNA gene included in the final assembled consensus sequence. Percent identify (%) indicates the top BLASTn hit. Isolate IDs with “-“ indicate samples where 16S sequencing failed.

**Supplementary Table S2. Wild bacterial isolate genome assembly statistics and quality metrics compared to reference genomes**

| Strain | Species ID (Accession) | Total Read Pairs | ANI vs. ref | Genome Size (bp) | Contigs | GC% | Contig N50 | Comp. (%) | Cont. (%) |
|---|---|---|---|---|---|---|---|---|---|
| **OP50** | *Escherichia coli* (GCA_004355015.1) | - | - | 4,563,825 | 74 | 50.7 | 115,119 | 100 | 0.15 |
| **ERW-16** | *Erwinia aphidicola* | 3,287,677 | 98.92 | 4,943,726 | 31 | 56.7 | 510,411 | 100 | 0.09 |
| **-** | *Erwinia aphidicola* ref (GCA_037149315.1) | - | - | 4,837,878 | 54 | 56.7 | 575,153 | 100 | 0.05 |
| **BAC-32** | *Bacillus glycinifermentans* | 3,942,783 | 99.9 | 4,471,938 | 106 | 46.2 | 103,394 | 100 | 0.81 |
| **-** | *Bacillus glycinifermentans* SRCM103574 (GCA_004103615.1) | - | - | 4,810,226 | 2 | 46 | 4,744,953 | 100 | 1 |
| **BAC-05** | *Priestia megaterium* | 4,030,919 | 96.06 | 5,925,444 | 35 | 37.5 | 2,978,497 | 100 | 2.99 |
| **BAC-39** | *Priestia megaterium* | 4,253,024 | 97.89 | 5,868,208 | 70 | 37.5 | 1,099,977 | 100 | 2.73 |
| **-** | *Priestia megaterium* ATCC 14581 (GCA_006094495.1) | - | - | 5,746,548 | 7 | 37.8 | 5,343,009 | 99.99 | 0.33 |
| **BAC-44** | *Bacillus altitudinis* | 4,059,276 | 98.04 | 3,731,020 | 18 | 41.3 | 534,178 | 100 | 0 |
| **-** | *Bacillus altitudinis* GR-8 (GCA_001191605.1) | - | - | 3,681,784 | 2 | 41.4 | 3,674,849 | 100 | 0 |
| **BAC-08** | *Bacillus safensis* | 2,726,591 | 98.86 | 3,649,452 | 15 | 41.7 | 947,972 | 100 | 0.08 |
| **-** | *Bacillus safensis* ref (GCA_039619585.1) | - | - | 3,703,645 | 2 | 41.7 | 3,697,266 | 100 | 0.07 |
| **BAC-29** | *Bacillus cereus* | 3,155,211 | 97.37 | 5,984,915 | 91 | 34.9 | 220,833 | 100 | 1.84 |
| **BAC-31** | *Bacillus cereus* | 4,060,472 | 97.25 | 6,254,551 | 104 | 34.7 | 149,624 | 100 | 0.07 |
| **BAC-36** | *Bacillus cereus* | 3,889,447 | 97.06 | 6,483,551 | 265 | 34.7 | 96,107 | 100 | 0.01 |
| **BAC-43** | *Bacillus cereus* | 4,786,152 | 97.26 | 5,815,293 | 83 | 34.8 | 547,666 | 100 | 0.02 |
| **-** | *Bacillus cereus* ATCC 14579 (GCA_006094295.1) | - | - | 5,431,377 | 2 | 35.3 | 5,416,249 | 100 | 0.61 |
| **BAC-06** | *Bacillus toyonensis* | 4,166,001 | 98.36 | 5,651,472 | 47 | 35.2 | 494,161 | 100 | 0.2 |
| **-** | *Bacillus toyonensis* ref (GCA_016605985.1) | - | - | 5,787,787 | 2 | 35.2 | 5,250,895 | 100 | 0.13 |
| **BAC-07** | *Bacillus paranthracis* | 4,111,942 | 98.8 | 5,404,450 | 59 | 35.3 | 231,477 | 100 | 0.13 |
| **-** | *Bacillus paranthracis* BtC4 (GCA_024296885.1) | - | - | 5,537,675 | 2 | 35.4 | 5,245,328 | 100 | 0.1 |

**Note:** ANI, Average Nucleotide Identity. bp, base pairs. N50, the shortest contig length such that 50% of the genome assembly is contained in contigs of this length or longer. Comp, Genome completeness. Cont, Genome contamination. Genome completeness and contamination were estimated using CheckM2. Reference accessions (GCA preflix) were retrieved from the NCBI Assembly database. ANI vs. ref denotes the percentage identity of the isolate genome compared to the reference genome listed immediately below it, as calculated by skani.

**Supplementary Table S3. Wild bacterial isolate functional genomic annotations compared to reference genomes**

| Strain | Species ID (Accession) | CDSs | rRNA | tRNA | tmRNA |
|---|---|---|---|---|---|
| **OP50** | *Escherichia coli*<br>(GCA_004355015.1) | 4233 | 11 | 77 | 1 |
| **ERW-16** | *Erwinia aphidicola* | 4444 | 9 | 82 | 1 |
| **-** | *Erwinia aphidicola* ref<br>(GCA_037149315.1) | 4434 | 9 | 79 | 1 |
| **BAC-32** | *Bacillus glycinifermentans* | 4605 | 9 | 71 | 1 |
| **-** | *Bacillus glycinifermentans*<br>SRCM103574<br>(GCA_004103615.1) | 5052 | 25 | 82 | 1 |
| **BAC-05** | *Priestia megaterium* | 6090 | 14 | 109 | 1 |
| **BAC-39** | *Priestia megaterium* | 6040 | 15 | 108 | 1 |
| **-** | *Priestia megaterium*<br>ATCC 14581<br>(GCA_006094495.1) | 5854 | 41 | 125 | 1 |
| **BAC-44** | *Bacillus altitudinis* | 3762 | 9 | 70 | 1 |
| **-** | *Bacillus altitudinis*<br>GR-8<br>(GCA_001191605.1) | 3691 | 24 | 81 | 1 |
| **BAC-08** | *Bacillus safensis* | 3670 | 9 | 67 | 1 |
| **-** | *Bacillus safensis* ref<br>(GCA_039619585.1) | 3682 | 27 | 90 | 1 |
| **BAC-29** | *Bacillus cereus* | 6024 | 15 | 102 | 1 |
| **BAC-31** | *Bacillus cereus* | 6145 | 17 | 101 | 1 |
| **BAC-36** | *Bacillus cereus* | 6592 | 15 | 104 | 1 |
| **BAC-43** | *Bacillus cereus* | 5743 | 21 | 107 | 1 |
| **-** | *Bacillus cereus*<br>ATCC 14579<br>(GCA_006094295.1) | 5421 | 39 | 108 | 1 |
| **BAC-06** | *Bacillus toyonensis* | 5661 | 16 | 90 | 1 |
| **-** | *Bacillus toyonensis* ref<br>(GCA_016605985.1) | 5666 | 42 | 106 | 1 |
| **BAC-07** | *Bacillus paranthracis* | 5498 | 13 | 86 | 1 |
| **-** | *Bacillus paranthracis*<br>BtC4<br>(GCA_024296885.1) | 5551 | 39 | 106 | 1 |

**Note:** CDSs, Coding DNA sequences. rRNA, ribosomal RNA. tRNA, transfer RNA, tmRNA, transfer-messenger RNA. All functional annotations for wild bacterial isolates and reference genomes were performed using Bakta v1.11.4 to ensure consistent feature identification.

**Supplementary Table S4. Composition and classification of Gene Cluster Families (GCFs) across wild bacterial isolates and reference genomes**

| Family | Category | Known MIBiG accession | Known compound | Wild Isolate | Species Reference |
|---|---|---|---|---|---|
| **GCF_01** | terpene | - | - | BAC-05, BAC-39 | *P. megaterium* |
| **GCF_02** | NRPS | - | - | - | *E. coli* |
| **GCF_03** | terpene | - | - | BAC-29, BAC-31, BAC-36, BAC-43, BAC-06, BAC-07 | *B. cereus, B. paranthracis, B. toyonensis* |
| **GCF_04** | RiPP | - | - | BAC-07 | *B. paranthracis* |
| **GCF_05** | terpene | - | - | BAC-29, BAC-31, BAC-36, BAC-43, BAC-06, BAC-07 | *B. cereus, B. glycinifermentans, B. paranthracis, B. toyonensis* |
| **GCF_06** | terpene | - | - | BAC-32, BAC-08, BAC-44 | *B. glycinifermentans, B. altitudinis, B. safensis* |
| **GCF_07** | NRPS | - | - | BAC-29, BAC-31, BAC-36, BAC-43 | - |
| **GCF_08** | other | - | - | BAC-29, BAC-31, BAC-36, BAC-43, BAC-06, BAC-07 | *B. cereus, B. paranthracis, B. toyonensis* |
| **GCF_09** | RiPP | - | - | BAC-08, BAC-44 | *B. altitudinis, B. safensis* |
| **GCF_10** | RiPP | - | - | ERW-16 | *E. aphidicola* |
| **GCF_11** | RiPP | - | - | BAC-06, BAC-07, BAC-29, BAC-36 | *B. toyonensis* |
| **GCF_12** | RiPP | - | - | BAC-31, BAC-43 | *B. cereus* |
| **GCF_13** | NRPS | - | - | BAC-08, BAC-44 | *B. safensis* |

| | | | | | |
|---|---|---|---|---|---|
| **GCF_14** | NRPS.PKS.RiPP | BGC0000626 / BGC0001059 | thuricin / zwittermicin A | BAC-31, BAC-36 | - |
| **GCF_15** | RiPP | - | | BAC-31, BAC-36, BAC-43 | - |
| **GCF_16** | RiPP | - | | - | *B. cereus* |
| **GCF_17** | NRPS | BGC0001185 / BGC0000401 / BGC0000309 / BGC0002695 | bacillibactin / paenibactin | BAC-32 | *B. glycinifermentans* |
| **GCF_18** | NRPS | - | | BAC-06, BAC-07, BAC-29, BAC-31, BAC-36, BAC-43 | *B. cereus, B. paranthracis, B. toyonensis* |
| **GCF_19** | RiPP | - | | BAC-06, BAC-07, BAC-29, BAC-31, BAC-36, BAC-43 | *B. cereus, B. paranthracis, B. toyonensis* |
| **GCF_20** | PKS | - | | BAC-05, BAC-39 | *P. megaterium* |
| **GCF_21** | other | - | | BAC-05, BAC-39 | *P. megaterium* |
| **GCF_22** | NRPS | BGC0000310 | bacitracin | BAC-32 x2 | *B. glycinifermentans* |
| **GCF_23** | terpene | - | | BAC-32 | *B. glycinifermentans* |
| **GCF_24** | RiPP | - | | BAC-44 | *B. altitudinis* |
| **GCF_25** | terpene | - | | ERW-16 | *E. aphidicola* |
| **GCF_26** | other | - | | BAC-08, BAC-44 | *B. altitudinis, B. safensis* |
| **GCF_27** | RiPP.terpene | BGC0000502 | cerecidin | - | *B. paranthracis* |
| **GCF_28** | NRPS | - | | BAC-08, BAC-44 | *B. altitudinis, B. safensis* |
| **GCF_29** | terpene | - | | BAC-05, BAC-39 | *P. megaterium* |
| **GCF_30** | RiPP | - | | BAC-32 | *B. glycinifermentans* |
| **GCF_31** | other | - | | BAC-05, BAC-39 | *P. megaterium* |
| **GCF_32** | NRPS | - | | BAC-29, BAC-31, | *B. cereus, B. toyonensis* |

| | | | | | |
|---|---|---|---|---|---|
| | | | | BAC-36, BAC-43 | |
| **GCF_33** | terpene | - | | BAC-08, BAC-44 | *B. altitudinis, B. safensis* |
| **GCF_34** | terpene | - | | BAC-05, BAC-39 | *P. megaterium* |
| **GCF_35** | RiPP | - | | BAC-06, BAC-29, BAC-36, BAC-43 | *B. cereus, B. toyonensis* |
| **GCF_36** | other | - | | BAC-32 | *B. glycinifermentans* |
| **GCF_37** | terpene | - | | BAC-05, BAC-39 | *P. megaterium* |
| **GCF_38** | other | BGC0000942 | petrobactin | BAC-06, BAC-29, BAC-31, BAC-36 x2, BAC-43 | *B. cereus, B. toyonensis* |
| **GCF_39** | terpene | - | | BAC-08, BAC-44 | *B. altitudinis, B. safensis* |
| **GCF_40** | PKS | - | | BAC-08, BAC-44 | *B. altitudinis, B. safensis* |
| **GCF_41** | NRPS | BGC0000381 | lichenysin | BAC-32 | *B. glycinifermentans* |
| **GCF_42** | other.terpene | - | | BAC-08, BAC-44 | *B. altitudinis, B. safensis* |
| **GCF_43** | RiPP | - | | BAC-32 | *B. glycinifermentans* |
| **GCF_44** | other | BGC0001572 | desferrioxamine E | ERW-16 | *E. aphidicola* |
| **GCF_45** | NRPS | - | | BAC-29, BAC-31, BAC-36, BAC-43 | *B. cereus* |
| **GCF_46** | NRPS | - | | ERW-16 | *E. aphidicola* |
| **GCF_47** | other | - | | ERW-16 | *E. aphidicola* |
| **GCF_48** | NRPS | - | | ERW-16 | *E. aphidicola* |
| **GCF_49** | other | - | | BAC-08, BAC-44 | *B. altitudinis, B. safensis* |
| **GCF_50** | PKS.RiPP | BGC0001887 | huazacin | BAC-29 | - |
| **GCF_51** | NRPS | - | | BAC-36, BAC-43 | - |
| **GCF_52** | RiPP | BGC0001173 | plantazolicin | BAC-08 | *B. safensis* |
| **GCF_53** | PKS | - | | BAC-32 | *B. glycinifermentans* |

| | | | | | |
|---|---|---|---|---|---|
| **GCF_54** | terpene | - | | BAC-32 | *B. glycinifermentans* |
| **GCF_55** | RiPP | - | | BAC-06 | *B. toyonensis* |
| **GCF_56** | other | - | | BAC-32 | *B. glycinifermentans* |
| **GCF_57** | other | - | | BAC-08, BAC-44 | *B. altitudinis, B. safensis* |
| **GCF_58** | RiPP | - | | ERW-16 | *E. aphidicola* |
| **GCF_59** | NRPS | - | | ERW-16 | *E. aphidicola* |
| **GCF_60** | NRPS | - | | BAC-07 | *B. paranthracis* |
| **GCF_61** | RiPP | - | | ERW-16 | - |
| **GCF_62** | RiPP | - | | BAC-06 | - |
| **GCF_63** | other | - | | ERW-16 | - |
| **GCF_64** | RiPP | - | | - | *B. safensis* |
| **GCF_65** | RiPP | - | | - | *P. megaterium* |
| **GCF_66** | RiPP | - | | BAC-43 | - |
| **GCF_67** | RiPP | - | | - | *B. cereus* |
| **GCF_68** | RiPP | - | | - | *P. megaterium* |
| **GCF_69** | RiPP | - | | - | *E. coli* |
| **GCF_70** | PKS | - | | BAC-08 | - |
| **GCF_71** | terpene | - | | - | *E. coli* |
| **GCF_72** | RiPP | - | | BAC-08 | - |
| **GCF_73** | RiPP | - | | BAC-29 | - |
| **GCF_74** | RiPP | - | | BAC-05 | - |
| **GCF_75** | RiPP | - | | BAC-05 | - |
| **GCF_76** | RiPP | - | | - | *B. safensis* |
| **GCF_77** | other | - | | - | *E. aphidicola* |
| **GCF_78** | RiPP | - | | BAC-31 | - |
| **GCF_79** | NRPS | - | | - | *B. toyonensis* |
| **GCF_80** | NRPS.PKS | - | | BAC-43 | - |
| **GCF_81** | RiPP | - | | BAC-06 | - |
| **GCF_82** | RiPP | - | | BAC-05 | - |
| **GCF_83** | other | - | | ERW-16 | - |
| **GCF_84** | RiPP | - | | - | *P. megaterium* |
| **GCF_85** | NRPS | - | | BAC-36 | - |
| **GCF_86** | RiPP | - | | BAC-31 | - |
| **GCF_87** | RiPP | - | | BAC-31 | - |
| **GCF_88** | NRPS | - | | BAC-44 | - |
| **GCF_89** | other | - | | - | *E. aphidicola* |

**Note:** GCF, Gene Cluster Family. NRPS, Non-Ribosomal Peptide Synthetase. PKS, Polyketide Synthase. RiPP, Ribosomally synthhesized and Post-translationally modified Peptide. MiBiG

Accession and Known Compound represent top hits against the MiBig database v4.0. Species reference indicate BGCs identified in reference genomes included in the BiG-SCAPE analysis. Multi-copy clusters within a single genome are indicated by “x2.”

### Assembled 16s rRNA gene sequences from wild bacterial isolates

The following section contains the final assembled 16S rRNA gene sequences for the 39 successfully sequenced isolates described in the study. Sequences were constructed using forward and reverse Sanger sequencing reads anchored to conserved internal regions C2, C3, and C4 and trimmed for quality as described in the Materials and Methods section. The strain ID, specific variable regions covered, and size of the sequence is provided in the header line. Additional details can be found in Supplementary Table S1.

```
>BAC-05 | V2-V9 | 1354 bp
GGCGGACGGGTGAGTAACACGTGGGCCACCTGCCTGTATGACTGGGATAACTTCGGGAAACCGATTCTAATACGGGATAGGATCTT
CTCCTTCATGGGAGATGATTGAAAGATGGTTTCGGCTATCACTTACAGATGGGCCCGCGGTGCATTAGCTAGTTGGTGATGTAACG
GCTCACCAAGGCAACGATGCATAGCCGACCTGAGAGGGTGATCGGCCACACTGGGACTGAGACACGGCCCAGACTCCTACGGGAGG
CAGCAGTAGGGATCTTCCGCCAATGGACGAAAGTCTGACGAGCACGCCGCGTGAGTGATGAAGGCTTTCGGGTCGTAAACTCTGTG
TTAGGGAAGAACAAGTACGAGAGTAACTGCTCGTACCTGACGGTACCTAACCAGAAAGCCACGGCTAACTACGTGCCAGCAGCCGC
GGTAATACGTAGGTGGCAAGCGTTATCCGGGAATTATTGGGCGTAAAGCGCGCGCAGGCGGTTTCTTAAGTCTGATGTGAAAGCCC
ACGGCTCAACCGTGGAGGGTCATTGGAAACTGGGGAACTTGAGTGCAGAAGAGAAAAGCGGAATTCCACGTGTAGCGGTGAAATGC
GTAGAGATGTGGAGGAACACCAGTGGCGAAGGCGGCTTTTTGGTCTGTAACTGACGCTGAGGCGCGAAAGCGTGGGGAGCAAACAG
GATTAGATACCCTGGTAGTCCACGCCGTAAACGATGAGTGCTAAGTGTTAGAGGGTTTCCGCCCTTTAGTGCTGCAGCTAACGCAT
TAAGCACTCCGCCTGGGGAGTACGGTCGCAAGACTGAAACTCAAAGGAATTGACGGGGGCCCGCACAAGCGGTGGAGCATGTGGTT
TAATTCGAAGCAACGCGAAGAACCTTACCAGGTCTTGACATCCTCTGACAACTCTAGAGATAGAGCGTTCCCCTTCGGGGGACAGA
GTGACAGGTGGTGCATGGTTGTCGTCAGCTCGTGTCGTGAGATGTTGGGTTAAGTCCCGCAACGAGCGCAACCCTTGATCTTAGTT
GCCAGCATTTAGTTGGGCACTCTAAGGTGACTGCCGGTGACAAACCGGAGGAAGGTGGGGATGACGTCAAATCATCATGCCCCTTA
TGACCTGGGCTACACACGTGCTACAATGGATGGTACAAAGGGCTGCAAGACCGCGAGGTCAAGCCAATCCCATAAAACCATTCTCA
GTTCGGATTGTAGGCTGCAACTCGCCTACATGAAGCTGGAATCGCTAGTAATCGCGGATCAGCATGCCGCGGTGAATACGTTCCCG
GGCCTTGTACACACCGCCCGTCACACCACGAGAGTTTGTAACACCCGAAGTCGGTGGAGTAACC
```

```
>BAC-06 | V2-V4 | 709 bp
GGCGGACGGGTGAGTAACACGTGGGTAACCTGCCCATAAGACTGGGATAACTCCGGGAAACCGGGGCTAATACCGGATAATATTTT
GAACTGCATGGTTCGAAATTGAAAGGCGGCTTCGGCTGTCACTTATGGATGGACCCGCGTCGCATTAGCTAGTTGGTGAGGTAACG
GCTCACCAAGGCAACGATGCGTAGCCGACCTGAGAGGGTGATCGGCCACACTGGGACTGAGACACGGCCCAGACTCCTACGGGAGG
CAGCAGTAGGGAATCTTCCGCAATGGACGAAAGTCTGACGGAGCAACGCCGCGTGAGTGATGAAGGCTTTCGGGTCGTAAAACTCT
GTTGTTAGGGAAGAACAAGTGCTCCTTGAATAAGCTGGCCCCTTGACGGTACCTACCCAGAAAGCCACGGCTAAAAAGGTGCTAGC
AGCCGCGGTAATACTTGGGTGTCAAGCGTTCTCAGGAATTTGGGGGCGTGCGCGGGGGGGGGGCGCTCAATTCCTTTGAGTTTCAA
CCTTGCGGGCGGACACCCCGAGCGGAGTGCGGAATGAGGTAACTTCTGCTCTGAAGAGGGAACCCCACTACACTCACGAGTCCACC
ATCTTTCGTCATGGACGAGAAAAAGGGCGGTAACTGATTGGTTACTCGCGCACTCGCGCGCGCGCCACCGCGACTGACAAAAAGCC
GGGGTCGCCACTATGTTCGTG
```

```
>BAC-07 | V3-V8 | 1078 bp
CTACGGGAGGCAGCAGTAGGGAATCTTCCGCAATGGACGAAAGTCTGACGGAGCAACGCCGCGTGAGTGATGAAGGCTTTCGGGTC
GTAAAACTCTGTTGTTAGGGAAGAACAAGTACAAGAGTAACTGCTTGTACCTTGACGGTACCTAACCAGAAAGCCACGGCTAACTA
CGTGCCAGCAGCCGCGGTAATACGTAGGTGGCAAGCGTTATCCGGAATTATTGGGCGTAAAGCGCGCGCAGGCGGTTTCTTAAGTC
TGATGTGAAAGCCCACGGCTCAACCGTGGAGGGTCATTGGAAACTGGGGAACTTGAGTGCAGAAGAGAAAAGCGGAATTCCACGTG
TAGCGGTGAAATGCGTAGAGATGTGGAGGAACACCAGTGGCGAAGGCGGCTTTTTGGTCTGTAACTGACGCTGAGGCGCGAAAGCG
TGGGGAGCAAACAGGATTAGATACCCTGGTAGTCCACGCCGTAAACGATGAGTGCTAAGTGTTAGAGGGTTTCCGCCCTTTAGTGC
```

TGCAGCTAACGCATTAAGCACTCCGCCTGGGGAGTACGGTCGCAAGACTGAAACTCAAAGGAATTGACGGGGGCCCGCACAAGCGG
TGGAGCATGTGGTTTAATTCGAAGCAACGCGAAGAACCTTACCAGGTCTTGACATCCTCTGACAACTCTAGAGATAGAGCGTTCCC
CTTCGGGGGACAGAGTGACAGGTGGTGCATGGTTGTCGTCAGCTCGTGTCGTGAGATGTTGGGTTAAGTCCCGCAACGAGCGCAAC
CCTTGATCTTAGTTGCCAGCATTTAGTTGGGCACTCTAAGGTGACTGCCGGTGACAAACCGGAGGAAGGTGGGGATGACGTCAAAT
CATCATGCCCCTTATGACCTGGGCTACACACGTGCTACAATGGATGGTACAAAGGGCTGCAAGACCGCGAGGTCAAGCCAATCCCA
TAAAACCATTCTCAGTTCGGATTGTAGGCTGCAACTCGCCTACATGAAGCTGGAATCGCTAGTAATCGCGGATCAGCATGCCGCGG
TGAATACGTTCCCGGGCCTTGTACACACCGCCCGTCACACCA
>BAC-08 | V2-V8 | 1260 bp
GGCGGACGGGTGAGTAACACGTGGGTAACCTGCCTGTAAGACTGGGATAACTCCGGGAAACCGGAGCTAATACCGGATAGTTCCTT
GAACCGCATGGTTCAAGGATGAAAGACGGTTTCGGCTGTCACTTACAGATGGACCCGCGGCGCATTAGCTAGTTGGTGGGGTAATG
GCTCACCAAGGCGACGATGCGTAGCCGACCTGAGAGGGTGATCGGCCACACTGGGACTGAGACACGGCCCAGACTCCTACGGGAGG
CAGCAGTAGGGAATCTTCCGCAATGGACGAAAGTCTGACGGAGCAACGCCGCGTGAGTGATGAAGGTTTTCGGATCGTAAAGCTCT
GTTGTTAGGGAAGAACAAGTGCGAGAGTAACTGCTCGCACCTTGACGGTACCTAACCAGAAAGCCACGGCTAACTACGTGCCAGCA
GCCGCGGTAATACGTAGGTGGCAAGCGTTGTCCGGAATTATTGGGCGTAAAGGGCTCGCAGGCGGTTTCTTAAGTCTGATGTGAAA
GCCCCCGGCTCAACCGGGGAGGGTCATTGGAAACTGGGAAACTTGAGTGCAGAAGAGGAGAGTGGAATTCCACGTGTAGCGGTGAA
ATGCGTAGAGATGTGGAGGAACACCAGTGGCGAAGGCGACTCTCTGGTCTGTAACTGACGCTGAGGAGCGAAAGCGTGGGGAGCGA
ACAGGATTAGATACCCTGGTAGTCCACGCCGTAAACGATGAGTGCTAAGTGTTAGGGGGTTTCCGCCCCTTAGTGCTGCAGCTAAC
GCATTAAGCACTCCGCCTGGGGAGTACGGTCGCAAGACTGAAACTCAAAGGAATTGACGGGGGCCCGCACAAGCGGTGGAGCATGT
GGTTTAATTCGAAGCAACGCGAAGAACCTTACCAGGTCTTGACATCCTCTGACAACCCTAGAGATAGGGCTTTCCCTTCGGGGACA
GAGTGACAGGTGGTGCATGGTTGTCGTCAGCTCGTGTCGTGAGATGTTGGGTTAAGTCCCGCAACGAGCGCAACCCTTGATCTTAG
TTGCCAGCATTCAGTTGGGCACTCTAAGGTGACTGCCGGTGACAAACCGGAGGAAGGTGGGGATGACGTCAAATCATCATGCCCCT
TATGACCTGGGCTACACACGTGCTACAATGGACAGAACAAAGGGCTGCAAGACCGCAAGGTTTAGCCAATCCCATAAATCTGTTCT
CAGTTCGGATCGCAGTCTGCAACTCGACTGCGTGAAGCTGGAATCGCTAGTAATCG
>PAN-09 | V2-V6 | 1018 bp
AGTGGCGGACGGGTGAGTAATGTCTGGGGATCTGCCCGATAGAGGGGGATAACCACTGGAAACGGTGGCTAATACCGCATAACGTC
GCAAGACCAAAGAGGGGGACCTTCGGGCCTCTCACTATCGGATGAACCCAGATGGGATTAGCTAGTAGGCGGGGTAATGGCCCACC
TAGGCGACGATCCCTAGCTGGTCTGAGAGGATGACCAGCCACACTGGAACTGAGACACGGTCCAGACTCCTACGGGAGGCAGCAGT
GGGGAATATTGCACAATGGGCGCAAGCCTGATGCAGCCATGCCGCGTGTATGAAGAAGGCCTTCGGGTTGTAAAGTACTTTCAGCG
GGGAGGAAGGCGATGCGGTTAATAACCGCGTCGATTGACGTTACCCGCAGAAGAAGCACCGGCTAACTCCGTGCCAGCAGCCGCGG
TAATACGGAGGGTGCAAGCGTTAATCGGAATTACTGGGCGTAAAGCGCACGCAGGCGGTCTGTTAAGTCAGATGTGAAATCCCCGG
GCTTAACCTGGGAACTGCATTTGAAACTGGCAGGCTTGAGTCTTGTAGAGGGGGGTAGAATTCCAGGTGTAGCGGTGAAATGCGTA
GAGATCTGGAGGAATACCGGTGGCGAAGGCGGCCCCCTGGACAAAGACTGACGCTCAGGTGCGAAAGCGTGGGGAGCAAACAGGAT
TAGATACCCTGGTAGTCCACGCCGTAAACGATGTCGACTTGGAGGTTGTTCCCTTGAGGAGTGGCTTCCGGAGCTAACGCGTTAAG
TCGACCGCCTGGGGAGTACGGCCGCAAGGTTAAAACTCAAATGAATTGACGGGGGCCCGCACAAGCGGTGGAGCATGTGGTTTAAT
TCGATGCAACGCGAAGAACCTTACCTACTCTTGACATCCAGCGGACTTTCCAGAGATGGATTGGTGCCTTCCGGGAACGCTGAGAC
AGTGCTGCATGGCTGTCGTCAGCTCGTGTTGTGAAATGTTGGGTTAAGTCCCGCAACGAgCGCAACCCTTA
>KOC-10 | V3-V8 | 994 bp
CTACGGGAGGCAGCAGTGGGGAATATTGCACAATGGGCGCAAGCTGATGCAGCGACGCCGCTGAGGGATGACGGCTTCGGTGTAAA
CCTCTTTCAGCACGGAAGAAGCGAAAGTGACGGTACGTGCAGAAGAAGCGCTGGCTAACTACGTGCCAGCAGCCGCGGTAATACGT
AGGGCGCAGCGTGTCCGGAATTATTGGGCGTAAAGAGCTCGTAGGCGGTTTGTCGCGTCTGCTGTGAAAGCCCGGGGCTTAACCCC
GGGTGTGCAGTGGGTACGGGCAGACTTGAGTGCAGTAGGGGAGACTGGAACTCCTGGTGTAGCGGTGAAATGCGCAGATATCAGGA
AGAACACCGATGGCGAAGGCAGGTCTCTGGGCTGTTACTGACGCTGAGGAGCGAAAGCATGGGGAGCGAACAGGATTAGATACCCT

GGTAGTCCATGCCGTAAACGTTGGGCACTAGGTGTGGGGGACATTCCACGTTTTCCGCGCCGTAGCTAACGCATTAAGTGCCCCGC
CTGGGGAGTACGGCCGCAAGGCTAAAACTCAAAGGAATTGACGGGGGCCCGCACAAGCGGCGGAGCATGCGGATTAATTCGATGCA
ACGCGAAGAACCTTACCAAGGCTTGCAAGAGCCTTCCCGGGATGGCTCAGAGATGGGTTTTCCTCCTTGTGGGGCTGGTGTACAGG
TGGTGCATGGTTGTCGTCAGCTCGTGTCGTGAGATGTTGGGTTAAGTCCCGCAACGAGCGCAACCCTCGTTCTATGTTGCCAGCAC
GTGATGGTGGGGACTCATAGGAGACTGCCGGGGTCAACTCGGAGGAAGGTGGGGATGACGTCAAATCATCATGCCCCTTATGTCTT
GGGCTTCACGCATGCTACAATGGCCAGTACAATGGGTTGCGATACCGTGAGGTGGAGCTAATCCCAAAAAGCTGGTCTCAGTTCGG
ATCGTGGTCTGCAACTCGACCACGTGAAGTCGGAGTCGCTAGTAATCG
>STA-11 | V3-V7 | 980 bp
GGTAACGGCTTACCAAGGCAACGATGCGTAGCCGACCTGAGAGGGTGATCGGCCACACTGGAACTGAGACACGGTCCAGACTCCTA
CGGGAGGCAGCAGTAGGGAATCTCGCAATGGGCGAAAGCCTGACGAGCATCGCCGCGTGAGTGATGAAGGTCTTCGGATCGTAAAA
CTCTGTTATTAGGGAGAACAAATGTGTAAGTAACTATGCACGTCTTGACGGTACTAATCAGAAAGCCACGGCTAACTACGTGCCAG
CAGCCGCGGTAATACGTAGGTGGCAAGCGTTATCTGGAATTATTGGGCGTAAAGCGCGCGTAGGCGGTTTTTTAAGTCTGATGTGA
AAGCCCACGGCTCAACCGTGGAGGGTCATTGGAAACTGGAAAACTTGAGTGCAGAAGAGGAAAGTGGAATTCCATGTGTAGCGGTG
AAATGCGCAGAGATATGGAGGAACACCAGTGGCGAAGGCGACTTCTGGTCTGTAACTGACGCTGATGTGCGAAAGCGTGGGGATCA
AACAGGATTAGATACCCTGGTAGTCCACGCCGTAAACGATGAGTGCTAAGTGTTAGGGGGTTTCCGCCCCTTAGTGCTGCAGCTAA
CGCATTAAGCACTCCGCCTGGGGAGTACGACCGCAAGGTTGAAACTCAAAGGAATTGACGGGGACCCGCACAAGCGGTGGAGCATG
TGGTTTAATTCGAAGCAACGCGAAGAACCTTACCAAATCTTGACATCCTCTGACCCCTCTAGAGATAGAGTTTTCCCCTTCGGGGG
ACAGAGTGACAGGTGGTGCATGGTTGTCGTCAGCTCGTGTCGTGAGATGTTGGGTTAAGTCCCGCAACGAGCGCAACCCTTAAGCT
TAGTTGCCATCATTAAGTTGGGCACTCTAAGTTGACTGCCGGTGACAAACCGGAGGAAGGTGGGGATGACGTCAAATCATCATGCC
CCTTATGATTTGGGCTACACACGTGCTACAATGG
>PSU-12 | V3-V8 | 1107 bp
TTAGCTAGTTGGTGGGGTAATGGCTCACCAAGGCGACGATCCGTAACTGGTCTGAGAGGATGATCAGTCACACTGGAACTGAGACA
CGGTCCAGACTCCTACGGGAGGCAGCAGTGGGGAATATTGGACAATGGGCGAAAGCCTGATCCAGCCATGCCGCGTGTGTGAAGAA
GGTCTTCGGATTGTAAAGCACTTTAAGTTGGGAGGAAGGGCAGTAAGCTAATACCTTGCTGTTTTGACTTTACCGACACAATAAGC
ACCGGCTAACTCTGTGCCAGCAGCCGCGGTAATACAGAGGGTGCAAGCGTTAATCGCAATTACTGGGCGTAAAGCGCGCGTAGGTG
GTTTGTTAAGTTGGATGTGAAAGCCCCGGGCTCAACCTGGGAACTGCATCCAAAACTGGCAAGCTAGAGTACGGTAGAGGGTGGTG
GAATTTCCTGTGTAGCGGTGAAATGCGTAGATATAGGAAGGAACACCAGTGGCGAAGGCGACCACCTGGACTGATACTGACACTGA
GGTGCGAAAGCGTGGGGAGCAAACAGGATTAGATACCCTGGTAGTCCACGCCGTAAACGATGTCAACTAGCCGTTGGAATCCTTGA
GATTTTAGTGGCGCAGCTAACGCATTAAGTTGACCGCCTGGGGAGTACGGCCGCAAGGTTAAAACTCAAATGAATTGACGGGGGCC
CGCACAAGCGGTGGAGCATGTGGTTTAATTCGAAGCAACGCGAAGAACCTTACCAGGCCTTGACATGCAGAGAACTTTCCAGAGAT
GGATTGGTGCCTTCGGGAACTCTGACACAGGTGCTGCATGGCTGTCGTCAGCTCGTGTCGTGAGATGTTGGGTTAAGTCCCGTAAC
GAGCGCAACCCTTGTCCTTAGTTACCAGCACGTTATGGTGGGCACTCTAAGGAGACTGCCGGTGACAAACCGGAGGAAGGTGGGGA
TGACGTCAAGTCATCATGGCCCTTACGGCCTGGGCTACACACGTGCTACAATGGTCGGTACAGAGGGTTGCCAAGCCGCGAGGTGG
AGCTAATCTCACAAAACCGATCGTAGTCCGGATCGCAGTCTGCAACTCGACTGCGTGAAGTCGGAATCGCTAGTA
>PSU-13 | V3-V8 | 1032 bp
CCAGACTCCTACGGGAGGCAGCAGTGGGGAATATTGGACAATGGGCGAAAGCTGATCCCAGCCATGCCGCGTGTGTGAAGAAGGTC
TCGGATTGTAAAGCACTTTAAGTTGGGAGGAAGAGCAGTTACTAATACGTGATTGTTTTGACGTTACCGACAGAATAAGCACCGGC
TAACTCTGTGCCAGCAGCCGCGGTAATACAGAGGGTGCAAGCGTTAATCGGAATTACTGGGCGTAAAGCGCGCGTAGGTGGTTTGT
TAAGTTGGATGTGAAATCCCCGGGCTCAACCTGGGAACTGCATTCAAAACTGACTGACTAGAGTATGGTAGAGGGTGGTGGAATTT
CCTGTGTAGCGGTGAAATGCGTAGATATAGGAAGGAACACCAGTGGCGAAGGCGACCACCTGGACTAATACTGACACTGAGGTGCG
AAAGCGTGGGGAGCAAACAGGATTAGATACCCTGGTAGTCCACGCCGTAAACGATGTCAACTAGCCGTTGGAAGCCTTGAGCTTTT
AGTGGCGCAGCTAACGCATTAAGTTGACCGCCTGGGGAGTACGGCCGCAAGGTTAAAACTCAAATGAATTGACGGGGGCCCGCACA

AGCGGTGGAGCATGTGGTTTAATTCGAAGCAACGCGAAGAACCTTACCAGGCCTTGACATCCAATGAACTTTCTAGAGATAGATTG
GTGCCTTCGGGAACATTGAGACAGGTGCTGCATGGCTGTCGTCAGCTCGTGTCGTGAGATGTTGGGTTACTATCCTGTAAATAGCG
CTCCCCTTGTCGTTAAACACCAGCAGCAACAGCCGGGCAATCTAAAGAGACGGACGGAGACAATCGGGAGGAAGGTGGGGATGATG
CCAAGTCTTCATGGCCCTTTCGACCGGGGATACACACCTGCTCCAATGGTCGGTCCAGAGGGTAGCCAACCCGAGAGGTGGAGGTA
ATCCCACAAAACCGATCGTAGTCCGGATCGCAGTAAGCAACTCGACTGCGTGAAGTTGGAATCCATAGTAATCGCGAATCAGAATG
>PSU-14 | V2-V8 | 1268 bp
GGCGGACGGGTGAGTAATGCCTAGGAATCTGCCTGGTAGTGGGGGATAACGTTCGGAAACGGACGCTAATACCGCATACGTCCTAC
GGGAGAAAGCAGGGGACCTTCGGGCCTTGCGCTATCAGATGAGCCTAGGTCGGATTAGCTAGTTGGTGGGGTAATGGCTCACCAAG
GCGACGATCCGTAACTGGTCTGAGAGGATGATCAGTCACACTGGAACTGAGACACGGTCCAGACTCCTACGGGAGGCAGCAGTGGG
GAATATTGGACAATGGGCGAAAGCCTGATCCAGCCATGCCGCGTGTGTGAAGAAGGTCTTCGGATTGTAAAGCACTTTAAGTTGGG
AGGAAGAGCAGTTACCTAATACGTGATTGTTTTGACGTTACCGACAGAATAAGCACCGGCTAACTCTGTGCCAGCAGCCGCGGTAA
TACAGAGGGTGCAAGCGTTAATCGGAATTACTGGGCGTAAAGCGCGCGTAGGTGGTTTGTTAAGTTGGATGTGAAATCCCCGGGCT
CAACTGGGAACTGCATTCAAAACTGACTGACTAGAGTATGGTAGAGGGTGGTGGAATTTCCTGTGTAGCGGTGAAATGCGTAGATA
TAGGAAGGAACACCAGTGGCGAAGGCGACCACCTGGACTAATACTGACACTGAGGTGCGAAAGCGTGGGGAGCAAACAGGATTAGA
TACCCTGGTAGTCCACGCCGTAAACGATGTCAACTAGCCGTTGGAAGCCTTGAGCTTTTAGTGGCGCAGCTAACGCATTAAGTTGA
CCGCCTGGGGAGTACGGCCGCAAGGTTAAAACTCAAATGAATTGACGGGGGCCCGCACAAGCGGTGGAGCATGTGGTTTAATTCGA
AGCAACGCGAAGAACCTTACCAGGCCTTGACATCCAATGAACTTTCTAGAGATAGATTGGTGCCTTCGGGAACATTGAGACAGGTG
CTGCATGGCTGTCGTCAGCTCGTGTCGTGAGATGTTGGGTTAAGTCCCGTAACGAGCGCAACCCTTGTCCTTAGTTACCAGCACGT
AATGGTGGGCACTCTAAGGAGACTGCCGGTGACAAACCGGAGGAAGGTGGGGATGACGTCAAGTCATCATGGCCCTTACGGCCTGG
GCTACACACGTGCTACAATGGTCGGTACAGAGGGTTGCCAAGCCGCGAGGTGGAGCTAATCCCACAAAACCGATCGTAGTCCGGAT
CGCAGTCTGCAACTCGACTGCGTGAAGTCGGAATCGCTAGTAATCGCGAATCAGAATGTCGCGG
>PSU-15 | V2-V8 | 1269 bp
GGCGGACGGGTGAGTAATGCCTAGGAATCTGCCTGGTAGTGGGGGATAACGTTCGGAAACGGACGCTAATACCGCATACGTCCTAC
GGGAGAAAGCAGGGGACCTTCGGGCCTTGCGCTATCAGATGAGCCTAGGTCGGATTAGCTAGTTGGTGGGGTAATGGCTCACCAAG
GCGACGATCCGTAACTGGTCTGAGAGGATGATCAGTCACACTGGAACTGAGACACGGTCCAGACTCCTACGGGAGGCAGCAGTGGG
GAATATTGGACAATGGGCGAAAGCCTGATCCAGCCATGCCGCGTGTGTGAAGAAGGTCTTCGGATTGTAAAGCACTTTAAGTTGGG
AGGAAGAGCAGTTACCTAATACGTGATTGTTTTGACGTTACCGACAGAATAAGCACCGGCTAACTCTGTGCCAGCAGCCGCGGTAA
TACAGAGGGTGCAAGCGTTAATCGGAATTACTGGGCGTAAAGCGCGCGTAGGTGGTTTGTTAAGTTGGATGTGAAATCCCCGGGCT
CAACCTGGGAACTGCATTCAAAACTGACTGACTAGAGTATGGTAGAGGGTGGTGGAATTTCCTGTGTAGCGGTGAAATGCGTAGAT
ATAGGAAGGAACACCAGTGGCGAAGGCGACCACCTGGACTAATACTGACACTGAGGTGCGAAAGCGTGGGGAGCAAACAGGATTAG
ATACCCTGGTAGTCCACGCCGTAAACGATGTCAACTAGCCGTTGGAAGCCTTGAGCTTTTAGTGGCGCAGCTAACGCATTAAGTTG
ACCGCCTGGGGAGTACGGCCGCAAGGTTAAAACTCAAATGAATTGACGGGGGCCCGCACAAGCGGTGGAGCATGTGGTTTAATTCG
AAGCAACGCGAAGAACCTTACCAGGCCTTGACATCCAATGAACTTTCTAGAGATAGATTGGTGCCTTCGGGAACATTGAGACAGGT
GCTGCATGGCTGTCGTCAGCTCGTGTCGTGAGATGTTGGGTTAAGTCCCGTAACGAGCGCAACCCTTGTCCTTAGTTACCAGCACG
TAATGGTGGGCACTCTAAGGAGACTGCCGGTGACAAACCGGAGGAAGGTGGGGATGACGTCAAGTCATCATGGCCCTTACGGCCTG
GGCTACACACGTGCTACAATGGTCGGTACAGAGGGTTGCCAAGCCGCGAGGTGGAGCTAATCCCACAAAACCGATCGTAGTCCGGA
TCGCAGTCTGCAACTCGACTGCGTGAAGTCGGAATCGCTAGTAATCGCGAATCAGAATGTCGCGG
>ERW-16 | V1-V9 | 1390 bp
ACACATGCAAGTCGTACCAGTGAGTCACACGTTCGCTTGCTCCTTGGGTGACGAGTGGCGGACGGGTGAGTAATGTCTGGGAAACT
GCCCGATGGAGGGGGATAACTACTGGAAACGGTAGCTAATACCGCATAACGTCTTCGGACCAAAGTGGGGGACCTTCGGGCCTCAC
ACCATCGGATGTGCCCAGATGGGATTAGCTAGTAGGTGGGGTAACGGCTCACCTAGGCGACGATCCCTAGCTGGTCTGAGAGGATG
ACCAGCCACACTGGAACTGAGACACGGTCCAGACTCCTACGGGAGGCAGCAGTGGGGAATATTGCACAATGGGCGCAAGCCTGATG

CAGCCATGCCGCGTGTATGAAGAAGGCCTTCGGGTTGTAAAGTACTTTCAGTGGGGAGGAAGGCGAAGAGGTTAATAACCTTTTCG
ATTGACGTTACCCGCAGAAGAAGCACCGGCTAACTCCGTGCCAGCAGCCGCGGTAATACGGAGGGTGCAAGCGTTAATCGGAATTA
CTGGGCGTAAAGCGCACGCAGGCGGTCTGTCAAGTCGGATGTGAAATCCCCGGGCTCAACCTGGGAACTGCATTCGAAACTGGCAG
GCTAGAGTCTTGTAGAGGGGGGTAGAATTCCAGGTGTAGCGGTGAAATGCGTAGAGATCTGGAGGAATACCGGTGGCGAAGGCGGC
CCCCTGGACAAAGACTGACGCTCAGGTGCGAAAGCGTGGGGAGCAAACAGGATTAGATACCCTGGTAGTCCACGCCGTAAACGATG
TCGACTTGGAGGTTGTGCCCTTGAGGCGTGGCTTCCGGAGCTAACGCGTTAAGTCGACCGCCTGGGGAGTACGGCCGCAAGGTTAA
AACTCAAATGAATTGACGGGGGCCCGCACAAGCGGTGGAGCATGTGGTTTAATTCGATGCAACGCGAAGAACCTTACCTGGCCTTG
ACATCCACGGAATTTGGCAGAGATGCCTTAGTGCCTTCGGGAACCGTGAGACAGGTGCTGCATGGCTGTCGTCAGCTCGTGTTGTG
AAATGTTGGGTTAAGTCCCGCAACGAGCGCAACCCTTATCCTTTGTTGCCAGCACGTAATGGTGGGAACTCAAAGGAGACTGCCGG
TGATAAACCGGAGGAAGGTGGGGATGACGTCAAGTCATCATGGCCCTTACGGCCAGGGCTACACACGTGCTACAATGGCGCATACA
AAGAGAAGCGACCTCGCGAGAGCAAGCGGACCTCATAAAGTGCGTCGTAGTCCGGATCGGAGTCTGCAACTCGACTCCGTGAAGTC
GGAATCGCTAGTAATCGTAGATCAGAATGCTACGGTGAATACGTTCCCGGGCCTTGTACACACCGCCCGTCACACCATGGGAGTGG
GTTGCAAAAGAAGT
>STA-17 | V2-V8 | 1262 bp
GGCGGACGGGTGAGTAACACGTGGATAACCTACCTATAAGACTGGGATAACTTCGGGAAACCGGAGCTAATACCGGATAAGATTTT
GAACCGCATGGTTCAATAGTGAAAGACGGCCTTGCTGTCACTTATAGATGGATCCGCGCCGTATTAGCTAGTTGGTAAGGTAACGG
CTTACCAAGGCAACGATACGTAGCCGACCTGAGAGGGTGATCGGCCACACTGGAACTGAGACACGGTCCAGACTCCTACGGGAGGC
AGCAGTAGGGAATCTTCCGCAATGGGCGAAAGCCTGACGGAGCAACGCCGCGTGAGTGATGAAGGTCTTCGGATCGTAAAACTCTG
TTATCAGGGAAGAACAAACGTGTAAGTAACTGTGCACGTCTTGACGGTACCTGATCAGAAAGCCACGGCTAACTACGTGCCAGCAG
CCGCGGTAATACGTAGGTGGCAAGCGTTATCCGGAATTATTGGGCGTAAAGCGCGCGTAGGCGGTTTTTTAAGTCTGATGTGAAAG
CCCACGGCTCAACCGTGGAGGGTCATTGGAAACTGGAAAACTTGAGTGCAGAAGAGGAAAGTGGAATTCCATGTGTAGCGGTGAAA
TGCGCAGAGATATGGAGGAACACCAGTGGCGAAGGCGACTTTCTGGTCTGTAACTGACGCTGATGTGCGAAAGCGTGGGGATCAAA
CAGGATTAGATACCCTGGTAGTCCACGCCGTAAACGATGAGTGCTAAGTGTTAGGGGGTTTCCGCCCCTTAGTGCTGCAGCTAACG
CATTAAGCACTCCGCCTGGGGAGTACGACCGCAAGGTTGAAACTCAAAGGAATTGACGGGGACCCGCACAAGCGGTGGAGCATGTG
GTTTAATTCGAAGCAACGCGAAGAACCTTACCAAATCTTGACATCCTTTGACCGCTCTAGAGATAGAGTTTTCCCCTTCGGGGGAC
AAAGTGACAGGTGGTGCATGGTTGTCGTCAGCTCGTGTCGTGAGATGTTGGGTTAAGTCCCGCAACGAGCGCAACCCTTAAGCTTA
GTTGCCATCATTAAGTTGGGCACTCTAAGTTGACTGCCGGTGACAAACCGGAGGAAGGTGGGGATGACGTCAAATCATCATGCCCC
TTATGATTTGGGCTACACACGTGCTACAATGGACAATACAAAGGGCAGCTAAACCGCGAGGTCAAGCAAATCCCATAAAGTTGTTC
TCAGTTCGGATTGTAGTCTGCAACTCGACTACATGAAGCTGGAATCGCTAGTAATCGT >KOC-18 | V2-V8 | 1239 bp
AGTGGCGAACGGGTGAGTAATACGTGAGTAACCTGCCCTTGACTCTGGGATAAGCCTGGGAAACTGGGTCTAATACTGGATACTAC
CTCTTACCGCATGGTGGGTGGTGGAAAGGGTTTTACTGGTTTTGGATGGGCTCACGGCCTATCAGCTTGTTGGTGGGGTAATGGCT
CACCAAGGCGACGACGGGTAGCCGGCCTGAGAGGGTGACCGGCCACACTGGGACTGAGACACGGCCCAGACTCCTACGGGAGGCAG
CAGTGGGGAATATTGCACAATGGGCGGAAGCCTGATGCAGCGACGCCGCGTGAGGGATGACGGCCTTCGGGTTGTAAACCTCTTTC
AGTAGGGAAGAAGCGAGAGTGACGGTACCTGCAGAAGAAGCGCCGGCTAACTACGTGCCAGCAGCCGCGGTAATACGTAGGGCGCA
AGCGTTGTCCGGAATTATTGGGCGTAAAGAGCTCGTAGGCGGTTTGTCGCGTCTGCTGTGAAAGCCCGGGGCTCAACCCCGGGTCT
GCAGTGGGTACGGGCAGACTAGAGTGCAGTAGGGGAGACTGGAATTCCTGGTGTAGCGGTGAAATGCGCAGATATCAGGAGGAACA
CCGATGGCGAAGGCAGGTCTCTGGGCTGTTACTGACGCTGAGGAGCGAAAGCATGGGGAGCGAACAGGATTAGATACCCTGGTAGT
CCATGCCGTAAACGTTGGGCACTAGGTGTGGGGGACATTCCACGTTTTCCGCGCCGTAGCTAACGCATTAAGTGCCCCGCCTGGGG
AGTACGGCCGCAAGGCTAAAACTCAAAGGAATTGACGGGGGCCCGCACAAGCGGCGGAGCATGCGGATTAATTCGATGCAACGCGA
AGAACCTTACCAAGGCTTGACATTCACCGGACCGCCCCAGAGATGGGGTTTCCCTTCGGGGCTGGTGGACAGGTGGTGCATGGTTG
TCGTCAGCTCGTGTCGTGAGATGTTGGGTTAAGTCCCGCAACGAGCGCAACCCTCGTTCTATGTTGCCAGCACGTGATGGTGGGGA
CTCATAGGAGACTGCCGGGGTCAACTCGGAGGAAGGTGGGGATGACGTCAAATCATCATGCCCCTTATGTCTTGGGCTTCACGCAT

GCTACAATGGCCGGTACAAAGGGTTGCGATACTGTGAGGTGGAGCTAATCCCAAAAAGCCGGTCTCAGTTCGGATTGAGGTCTGCA
ACTCGACCTCATGAAGTCGGAGTCGCTAGTAATCG
>PSU-19 | B2-V8 | 1252 bp
GGCGGACGGGAGAGTAATGCCTAGGAATCTGCCTGGTAGTGGGGGATAACGTTCGGAAACGGACGCTAATACCGCATACGTCCTAC
GGGAGAAAGCAGGGGACCTTCGGGCCTTGCGCTATCAGATGAGCCTAGGTCGGATTAGCTAGTTGGTGGGGTAATGGCTCACCAAG
GCGACGATCCGTAACTGGTCTGAGAGGATGATCAGTCACACTGGAACTGAGACACGGTCCAGACTCCTACGGGAGGCAGCAGTGGG
GAATATTGGACAATGGGCGAAAGCCTGATCCAGCCATGCCGCGTGTGTGAAGAAGGTCTTCGGATTGTAAAGCACTTTAAGTTGGG
AGGAAGAGCAGTTACCTAATACGTGATTGTTTTGACGTTACCGACAGAATAAGCACCGGCTAACTCTGTGCCAGCAGCCGCGGTAA
TACAGAGGGTGCAAGCGTTAATCGGAATTACTGGGCGTAAAGCGCGCGTAGGTGGTTTGTTAAGTTGGATGTGAAATCCCCGGGCT
CAACCTGGGAACTGCATTCAAAACTGACTGACTAGAGTATGGTAGAGGGTGGTGGAATTTCCTGTGTAGCGGTGAAATGCGTAGAT
ATAGGAAGGAACACCAGTGGCGAAGGCGACCACCTGGACTAATACTGACACTGAGGTGCGAAAGCGTGGGGAGCAAACAGGATTAG
ATACCCTGGTAGTCCACGCCGTAAACGATGTCAACTAGCCGTTGGAAGCCTTGAGCTTTTAGTGGCGCAGCTAACGCATTAAGTTG
ACCGCCTGGGGAGTACGGCCGCAAGGTTAAAACTCAAATGAATTGACGGGGGCCCGCACAAGCGGTGGAGCATGTGGTTTAATTCG
AAGCAACGCGAAGAACCTTACCAGGCCTTGACATCCAATGAACTTTCTAGAGATAGATTGGTGCCTTCGGGAACATTGAGACAGGT
GCTGCATGGCTGTCGTCAGCTCGTGTCGTGAGATGTTGGGTTAAGTCCCGTAACGAGCGCAACCCTTGTCCTTAGTTACCAGCACG
TAATGGTGGGCACTCTAAGGAGACTGCCGGTGACAAACCGGAGGAAGGTGGGGATGACGTCAAGTCATCATGGCCCTTACGGCCTG
GGCTACACACGTGCTACAATGGTCGGTACAGAGGGTTGCCAAGCCGCGAGGTGGAGCTAATCCCACAAAACCGATCGTAGTCCGGA
TCGCAGTCTGCAACTCGACTGCGTGAAGTCGGAATCGCTAGTAATCG
>MCR-20 | V3-V8 | 1149 bp
TCAGCTTGTTGGTGAGGTAATGGCTCACCAAGGCGTCGACGGGTAGCCGGCCTGAGAGGGTGACCGGCCACACTGGGACTGAGACA
CGGCCCAGACTCCTACGGGAGGCAGCAGTGGGGAATATTGCACAATGGGCGCAAGCCTGATGCAGCAACGCCGCGTGAGGGATGAC
GGCCTTCGGGTTGTAAACCTCTTTTAGCAGGGAAGAAGCGAAAGTGACGGTACCTGCAGAAAAAGCACCGGCTAACTACGTGCCAG
CAGCCGCGGTAATACGTAGGGTGCAAGCGTTATCCGGAATTATTGGGCGTAAAGAGCTCGTAGGCGGTTTGTCGCGTCTGCTGTGA
AATTCCGAGGCTCAACCTCGGGCTTGCAGTGGGTACGGGCAGACTAGAGTGCGGTAGGGGAGATTGGAATTCCTGGTGTAGCGGTG
GAATGCGCAGATATCAGGAGGAACACCGATGGCGAAGGCAGATCTCTGGGCCGTAACTGACGCTGAGGAGCGAAAGGGTGGGGAGC
AAACAGGCTTAGATACCCTGGTAGTCCACCCCGTAAACGTTGGGAACTAGTTGTGGGGTCCTTTCCACGGATTCCGTGACGCAGCT
AACGCATTAAGTTCCCCGCCTGGGGAGTACGGCCGCAAAGCTAAAACTCAAAGGAATTGACGGGGACCCGCACAAGCGGCGGAGCA
TGCGGATTAATTCGATGCAACGCGAAGAACCTTACCAAGGCTTGACATACACGAGAACGGGCCAGAAATGGTCAACTCTTTGGACA
CTCGTGAACAGGTGGTGCATGGTTGTCGTCAGCTCGTGTCGTGAGATGTTGGGTTAAGTCCCGCAACGAGCGCAACCCTCGTTCTA
TGTTGCCAGCACGTAATGGTGGGAACTCATGGGATACTGCCGGGGTCAACTCGGAGGAAGGTGGGGATGACGTCAAATCATCATGC
CCCTTATGTCTTGGGCTTCACGCATGCTACAATGGCCGGTACAATGGGCTGCGATACCGTAAGGTGGAGCGAATCCCAAAAAGCCG
GTCCCAGTTCGGATTGAGGTCTGCAACTCGACCTCATGAAGTCGGAGTCGCTAGTAATCGCAGATCAGCAACGCTGCGGTGAATAC
GTTCCCGGGTCTTGTACACACCGCCCGTCA
>MIC-21 | V3-V8 | 1096 bp
ATCAGCTTGTTGGTGAGGTAATGGCTCACCAAGGCGACGACGGGTAGCCGGCCTGAGAGGGTGACCGGCCACACTGGGACTGAGAC
ACGGCCCAGACTCCTACGGGAGGCAGCAGTGGGGAATATTGCACAATGGGCGCAAGCCTGATGCAGCGACGCCGCGTGAGGGATGA
CGGCCTTCGGGTTGTAAACCTCTTTCAGTAGGGAAGAAGCGAAAGTGACGGTACCTGCAGAAGAAGCACCGGCTAACTACGTGCCA
GCAGCCGCGGTAATACGTAGGGTGCGAGCGTTATCCGGGAATTATTGGGCGTAAAGAGCTCGTAGGCGGTTTGTCGCGTCTGTCGT
GAAAGTCCGGGGCTTAACCCCGGGATCTGCGGTGGGTACGGGCAGACTAGAGTGCAGTAGGGGAGACTGGAATTCCTGGTGTAGCG
GTGGAATGCGCAGATATCAGGAGGAACACCGATGGCGAAGGCAGGTCTCTGGGCTGTAACTGACGCTGAGGAGCGAAAGCATGGGG
AGCGAACAGGATTAGATACCCTGGTAGTCCATGCCGTAAACGTTGGGCACTAGGTGTGGGGACCATTCCACGGTTTCCGCGCCGCA
GCTAACGCATTAAGTGCCCCGCCTGGGGAGTACGGCCGCAAGGCTAAAACTCAAAGGAATTGACGGGGGCCCGCACAAGCGGCGGA

GCATGCGGATTAATTCGATGCAACGCGAAGAACCTTACCAAGGCTTGACATGTTCTCGATCGCCGTAGAGATACGGTTTCCCCTTT
GGGGCGGGTTCACAGGTGGTGCATGGTTGTCGTCAGCTCGTGTCGTGAGATGTTGGGTTAAGTCCCGCAACGAGCGCAACCCTCGT
TCCATGTTGCCAGCACGTAATGGTGGGGACTCATGGGAGACTGCCGGGGTCAACTCGGAGGAAGGTGAGGACGACGTCAAATCATC
ATGCCCCTTATGTCTTGGGCTTCACGCATGCTACAATGGCCGGTACAATGGGTTGCGATACTGTGAGGTGGAGCTAATCCCAAAAA
GCCGGTCTCAGTTCGGATTGGGGTCTGCAACTCGACCCCATGAAGTCGGAGTCGCTAGTAATCG
>PSU-22 | V2-V9 | 1335 bp
GGCGGACGGGTGAGTAATGCCTAGGAATCTGCCTGGTAGTGGGGGATAACGTTCGGAAACGGACGCTAATACCGCATACGTCCTAC
GGGAGAAAGCAGGGGACCTTCGGGCCTTGCGCTATCAGATGAGCCTAGGTCGGATTAGCTAGTTGGTGGGGTAATGGCTCACCAAG
GCGACGATCCGTAACTGGTCTGAGAGGATGATCAGTCACACTGGAACTGAGACACGGTCCAGACTCCTACGGGAGGCAGCAGTGGG
GAATATTGGACAATGGGCGAAAGCCTGATCCAGCCATGCCGCGTGTGTGAAGAAGGTCTTCGGATTGTAAAGCACTTTAAGTTGGG
AGGAAGAGCAGTTACCTAATACGTGATTGTTTTGACGTTACCGACAGAATAAGCACCGGCTAACTCTGTGCCAGCAGCCGCGGTAA
TACAGAGGGTGCAAGCGTTAATCGGAATTACTGGGCGTAAAGCGCGCGTAGGTGGTTTGTTAAGTTGGATGTGAAATCCCCGGGCT
CAACCTGGGAACTGCATTCAAAACTGACTGACTAGAGTATGGTAGAGGGTGGTGGAATTTCCTGTGTAGCGGTGAAATGCGTAGAT
ATAGGAAGGAACACCAGTGGCGAAGGCGACCACCTGGACTAATACTGACACTGAGGTGCGAAAGCGTGGGGAGCAAACAGGATTAG
ATACCCTGGTAGTCCACGCCGTAAACGATGTCAACTAGCCGTTGGAAGCCTTGAGCTTTTAGTGGCGCAGCTAACGCATTAAGTTG
ACCGCCTGGGGAGTACGGCCGCAAGGTTAAAACTCAAATGAATTGACGGGGGCCCGCACAAGCGGTGGAGCATGTGGTTTAATTCG
AAGCAACGCGAAGAACCTTACCAGGCCTTGACATCCAATGAACTTTCTAGAGATAGATTGGTGCCTTCGGGAACATTGAGACAGGT
GCTGCATGGCTGTCGTCAGCTCGTGTCGTGAGATGTTGGGTTAAGTCCCGTAACGAGCGCAACCCTTGTCCTTAGTTACCAGCACG
TAATGGTGGGCACTCTAAGGAGACTGCCGGTGACAAACCGGAGGAAGGTGGGGATGACGTCAAGTCATCATGGCCCTTACGGCCTG
GGCTACACACGTGCTACAATGGTCGGTACAGAGGGTTGCCAAGCCGCGAGGTGGAGCTAATCCCACAAAACCGATCGTAGTCCGGA
TCGCAGTCTGCAACTCGACTGCGTGAAGTCGGAATCGCTAGTAATCGCGAATCAGAATGTCGCGGTGAATACGTTCCCGGGCCTTG
TACACACCGCCCGTCACACCATGGGAGTGGGTTGCACCAGAAGTA
>PSU-23 | V3-V9 | 1089 bp
CTACGGGAGGCAGCAGTGGGGAATATTGGACATGGGCGAAAGCTGATCCAGCATGCGCGTGTGTGAGAGTCTCGATGTAAAGCACT
TTAAGTTGGGAGAAGGGTAGTACTTATACGTTGCTACTTTGACGTTACCGACAGAATAGCACGGCTAACTTCGTGCCAGCAGCCGC
GGTAATACGAAGGGTGCAAGCGTTAATCGGAATTACTGGGCGTAAAGCGCGCGTAGGTGGTTCAGTAAGTTGGAAGTGAAATCCCC
GGGCTCAACCTGGGAACTGCTTTCAAAACTGCTGAGCTAGAGTACGGTAGAGGGTGGTGGAATTTCCTGTGTAGCGGTGAAATGCG
TAGATATAGGAAGGAACACCAGTGGCGAAGGCGACCACCTGGACTGATACTGACACTGAGGTGCGAAAGCGTGGGGAGCAAACAGG
ATTAGATACCCTGGTAGTCCACGCCGTAAACGATGTCAACTAGCCGTTGGGAGTCTTGAACTTTTAGTGGCGCAGCTAACGCATTA
AGTTGACCGCCTGGGGAGTACGGCCGCAAGGTTAAAACTCAAATGAATTGACGGGGGCCCGCACAAGCGGTGGAGCATGTGGTTTA
ATTCGAAGCAACGCGAAGAACCTTACCTGGCCTTGACATGCTGAGAACTTTCTAGAGATAGATTGGTGCCTTCGGGAACTCAGACA
CAGGTGCTGCATGGCTGTCGTCAGCTCGTGTCGTGAGATGTTGGGTTAAGTCCCGTAACGAGCGCAACCCTTGTCCTTAGTTACCA
GCACGTTATGGTGGGAACTCTAAGGAGACTGCCGGTGACAAACCGGAGGAAGGTGGGGATGACGTCAAGTCATCATGGCCCTTACG
GCCAGGGCTACACACGTGCTACAATGGTCGGTACAAAGGGTTGCCAAGCCGCGAGGTGGAGCTAATCCCATAAAACCGATCGTAGT
CCGGATCGCAGTCTGCAACTCGACTGCGTGAAGTCGGAATCGCTAGTAATCGTGAATCAGAATGTCACGGTGAATACGTTCCCGGG
CCTTGTACACACCGCCCGTCACACCATGGGAGTGGGTTGCACCAGAAGTAGC
>COR-24 | V2-V8 | 1229 bp
GGCGAACGGTGGAGTAACTGCTGGGTGAGCTGCCCTACACTTTGGGATAAGCCTGGGAAACTGGGTCTAATACCGAATATTCCCAC
CAATGTAGGGGTGGTGTGGAAAGCCTGGATCTGTGTGGGATGACCCTGCCGCATCTCCTCTTGTTGGTGGGGCCTGGCCTTTCAT
TGCCTCTAGGGTCAGCCTGCCTGAAGGTTGTACTAACACATTGGGACTGAGACACGGCCCAGACTCCTACGGGAGGCAGCAGTGGG
GAATATTGCACAATGGGCGCAAGCCTGATGCAGCGACGCCGCGTGGGGGATGACGGCCTTCGGGTTGTAAACTCCTTTCGCTAGGG
ACGAAGCCTTTTTGGTGACGGTACCTGGAGAAGAAGCACCGGCTAACTACGTGCCAGCAGCCGCGGTAATACGTAGGGTGCGAGCG

TTGTCCGGAATTACTGGGCGTAAAGAGCTCGTAGGTGGTTTGTCGCGTCGTCTGTGAAATCCCGGGGCTTAACTTCGGGCGTGCAG
GCGATACGGGCATAACTTGAGTGCTGTAGGGGAGACTGGAATTCCTGGTGTAGCGGTGAAATGCGCAGATATCAGGAGGAACACCA
ATGGCGAAGGCAGGTCTCTGGGCAGTAACTGACGCTGAGGAGCGAAAGCATGGGTAGCGAACAGGATTAGATACCCTGGTAGTCCA
TGCCGTAAACGGTGGGCGCTAGGTGTAGGGGTCTTCCACGACTTCTGTGCCGCAGCTAACGCATTAAGCGCCCCGCCTGGGGAGTA
CGGCCGCAAGGCTAAAACTCAAAGGAATTGACGGGGGCCCGCACAAGCGGCGGAGCATGTGGATTAATTCGATGCAACGCGAAGAA
CCTTACCTGGGCTTGACATGGACCGGATCGGCGTAGAGATACGTTTTCCCTTGTGGTCGGTTCACAGGTGGTGCATGGTTGTCGTC
AGCTCGTGTCGTGAGATGTTGGGTTAAGTCCCGCAACGAGCGCAACCCTTGTCTTATGTTGCCAGCACATTATGGTGGGTACTCAT
GAGAGACTGCCGGGGTTAACTCGGAGGAAGGTGGGGATGACGTCAAATCATCATGCCCCTTATAGACTGGGATCCTGGCTCAGCGC
TAGGTTCAGCACAGGGAGCTGCCTCACGACAGGTGCAGGAGCTTCCTCGAAGCGCGCCTCAGTTCGGCAGGGGTCTGCAACTCGAC
CCCTTGAAGTGCATCTCGCTAGGA
>RHO-25 | V2-V8 | 1233 bp
GGCGAACGGGAGAGTAACACGTGGGTGATCTGCCCTGCACTCTGGGATAAGCCTGGGAAACTGGGTCTAATACCGGATATGACCTC
TTGCTGCATGGCGAGGGGTGGAAAGTTTTTCGGTGCAGGATGAGCCCGCGGCCTATCAGCTTGTTGGTGGGGTAATGGCCTACCAA
GGCGACGACGGGTAGCCGGCCTGAGAGGGCGACCGGCCACACTGGGACTGAGACACGGCCCAGACTCCTACGGGAGGCAGCAGTGG
GGAATATTGCACAATGGGCGAAAGCCTGATGCAGCGACGCCGCGTGAGGGATGACGGCCTTCGGGTTGTAAACCTCTTTCAGCAGG
GACGAAGCGAAAGTGACGGTACCTGCAGAAGAAGCACCGGCCAACTACGTGCCAGCAGCCGCGGTAATACGTAGGGTGCGAGCGTT
GTCCGGAATTACTGGGCGTAAAGAGCTCGTAGGCGGTTTGTCGCGTCGTCTGTGAAATCCCGCAGCTCAACTGCGGGCTTGCAGGC
GATACGGGCAGACTCGAGTACTGCAGGGGAGACTGGAATTCCTGGTGTAGCGGTGAAATGCGCAGATATCAGGAGGAACACCGGTG
GCGAAGGCGGGTCTCTGGGCAGTAACTGACGCTGAGGAGCGAAAGCGTGGGTAGCGAACAGGATTAGATACCCTGGTAGTCCACGC
CGTAAACGGTGGGCGCTAGGTGTGGGTTTCCTTCCACGGGATCCGTGCCGTAGCCAACGCATTAAGCGCCCCGCCTGGGGAGTACG
GCCGCAAGGCTAAAACTCAAAGGAATTGACGGGGGCCCGCACAAGCGGCGGAGCATGTGGATTAATTCGATGCAACGCGAAGAACC
TTACCTGGGTTTGACATGTACCGGACGACTGCAGAGATGTGGTTTCCCTTGTGGCCGGTAGACAGGTGGTGCATGGCTGTCGTCAG
CTCGTGTCGTGAGATGTTGGGTTAAGTCCCGCAACGAGCGCAACCCTTGTCCTGTGTTGCCAGCACGTAATGGTGGGGACTCGCAG
GAGACTGCCGGGGTCAACTCGGAGGAAGGTGGGGACGACGTCAAGTCATCATGCCCCTTATGTCCAGGGCTTCACACATGCTACAA
TGGTCGGTACAGAGGGCTGCGATACCGTGAGGTGGAGCGAATCCCTTAAAGCCGGTCTCAGTTCGGATCGGGGTCTGCAACTCGAC
CCCGTGAAGTCGGAGTCGCTAGTAATCG
>STA-26 | V2-V8 | 1322 bp
AGCGGCGGACGGGTGAGTAACACGTAGGTAACCTACCTATAAGACTGGGATAACTTCGGGAAACCGGAGCTAATACCGGATAATAT
TTCGAACCGCATGGTTCGATAGTGAAAGATGGCTTTGCTATCACTTATAGATGGACCTGCGCCGTATTAGCTAGTTGGTAAGGTAA
CGGCTTACCAAGGCAACGATACGTAGCCGACCTGAGAGGGTGATCGGCCACACTGGAACTGAGACACGGTCCAGACTCCTACGGGA
GGCAGCAGTAGGGAATCTTCCGCAATGGGCGAAAGCCTGACGGAGCAACGCCGCGTGAGTGATGAAGGTCTTCGGATCGTAAAACT
CTGTTATTAGGGAAGAACAAACGTGTAAGTAACTGTGCACGTCTTGACGGTACCTAATCAGAAAGCCACGGCTAACTACGTGCCAG
CAGCCGCGGTAATACGTAGGTGGCAAGCGTTATCCGGAATTATTGGGCGTAAAGCGCGCGTAGGCGGTTTTTTAAGTCTGATGTGA
AAGCCCACGGCTCAACCGTGGAGGGTCATTGGAAACTGGAAAACTTGAGTGCAGAAGAGGAAAGTGGAATTCCATGTGTAGCGGTG
AAATGCGCAGAGATATGGAGGAACACCAGTGGCGAAGGCGACTTTCTGGTCTGTAACTGACGCTGATGTGCGAAAGCGTGGGGATC
AAACAGGATTAGATACCCTGGTAGTCCACGCCGTAAACGATGAGTGCTAAGTGTTAGGGGGTTTCCGCCCCTTAGTGCTGCAGCTA
ACGCATTAAGCACTCCGCCTGGGGAGTACGACCGCAAGGTTGAAACTCAAGGAATTGACGGGACCCGCACAAGCGGTGGAGCATGT
GGTTTAATTCGAAGCAACGCGAAGAACCTTACCAAATCTTGACATCCTTTGACCCTTCTAGAGATAGAAGTTTCCCCTTCGGGGGA
CAAAGTGACAGGTGGTGCATGGTTGTCGTCAGCTCGTGTCGTGAGATGTTGGGTTAAGTCCCGCAACGAGCGCAACCCTTAAGCTT
AGTTGCCATCATTAAGTTGGGCACTCTAAGTTGACTGCCGGTGACAAACCGGAGGAAGGTGGGGATGACGTCAAATCATCATGCCC
CTTATGATTTGGGCTACACACGTGCTACAATGGACAATACAAAGGGCAGCGAAACCGCGAGGTCAAGCAAATCCCATAAAGTTGTT

CTCAGTTCGGATTGTAGTCTGCAACTCGACTACATGAAGCTGGAATCGCTAGTAATCGTAGATCAGCATGCTACGGTGAATACGTT
CCCGGGTCTTGTACACACCGCCCGTCACACCA
>STA-27 | V3-V5 | 649 bp
ATTAGCTAGTTGGTAAGGTAACGGCTTACCAAGGCAACGATACGTAGCCGACCTGAGAGGGTGATCGGCCACACTGGAACTGAGAC
ACGGTCCAGACTCCTACGGGAGGCAGCAGTAGGGAATCTTCCGCAATGGGCGAAAGCCTGACGGAGCAACGCCGCGTGAGTGATGA
AGGTCTTCGGATCGTAAAACTCTGTTATCAGGGAAGAACAAACGTGTAAGTAACTGTGCACGTCTTGACGGTACCTGATCAGAAAG
CCACGGCTAACTACGTGCCAGCAGCCGCGGTAATACGTAGGTGGCAAGCGTTATCCGGAATTATTGGGCGTAAAGCGCGCGTAGGC
GGTTTTTTAAGTCTGATGTGAAAGCCCACGGCTCAACCGTGGAGGGTCATTGGAAACTGGAAAACTTGAGTGCAGAAGAGGAAAGT
GGAATTCCATGTGTAGCGGTGAAATGCGCAGAGATATGGAGGAGCACCAGTGGCGAAGGCGACTTTCTGGTCTGTAACTGACGCTG
ATGTGCGAAAGCGTGGGGATCAAACAGGATTAGATACCCTGGTAGTCCACGCCGTAAACGATGAGTGCTAAGTGTTAGGGGGTTTC
CGCCCCTTAGTGCTGCCGCTAACGCATTAAGCACTCCGCCTGGGGAG
>ENT-28 | V1-V9 | 1399 bp
ACGGTAACAGGAAGCAGCTTGCTGCTTCGCTGACGAGTGGCGGACGGGTGAGTAATGTCTGGGAAACTGCCTGATGGAGGGGGATA
ACTACTGGAAACGGTAGCTAATACCGCATAACGTCGCAAGACCAAAGAGGGGGACCTTCGGGCCTCTTGCCATCGGATGTGCCCAG
ATGGGATTAGCTTGTTGGTGGGGTAACGGCTCACCAAGGCGACGATCCCTAGCTGGTCTGAGAGGATGACCAGCCACACTGGAACT
GAGACACGGTCCAGACTCCTACGGGAGGCAGCAGTGGGGAATATTGCACAATGGGCGCAAGCCTGATGCAGCCATGCCGCGTGTAT
GAAGAAGGCCTTCGGGTTGTAAAGTACTTTCAGCGGGGAGGAAGGCGGTACGGTTAATAACCGTGCCGATTGACGTTACCCGCAGA
AGAAGCACCGGCTAACTCCGTGCCAGCAGCCGCGGTAATACGGAGGGTGCAAGCGTTAATCGGAATTACTGGGCGTAAAGCGCACG
CAGGCGGTCTGTCAAGTCGGATGTGAAATCCCCGGGCTCAACCTGGGAACTGCATCCGAAACTGGCAGGCTAGAGTCTTGTAGAGG
GGGGTAGAATTCCAGGTGTAGCGGTGAAATGCGTAGAGATCTGGAGGAATACCGGTGGCGAAGGCGGCCCCCTGGACAAAGACTGA
CGCTCAGGTGCGAAAGCGTGGGGAGCAAACAGGATTAGATACCCTGGTAGTCCACGCCGTAAACGATGTCGACTTGGAGGTTGTGC
CCTTGAGGCGTGGCTTCCGGAGCTAACGCGTTAAGTCGACCGCCTGGGGAGTACGGCCGCAAGGTTAAAACTCAAATGAATTGACG
GGGGCCCGCACAAGCGGTGGAGCATGTGGTTTAATTCGATGCAACGCGAAGAACCTTACCTACTCTTGACATCCACAGAACTTGGC
AGAGATGCTTTGGTGCCTTCGGGAACTGTGAGACAGGTGCTGCATGGCTGTCGTCAGCTCGTGTTGTGAAATGTTGGGTTAAGTCC
CGCAACGAGCGCAACCCTTATCCTTTGTTGCCAGCGAGTAATGTCGGGAACTCAAAGGAGACTGCCAGTGATAAACTGGAGGAAGG
TGGGGATGACGTCAAGTCATCATGGCCCTTACGAGTAGGGCTACACACGTGCTACAATGGCGCATACAAAGAGAAGCGACCTCGCG
AGAGCAAGCGGACCTCATAAAGTGCGTCGTAGTCCGGATCGGAGTCTGCAACTCGACTCCGTGAAGTCGGAATCGCTAGTAATCGT
GGATCAGAATGCCACGGTGAATACGTTCCCGGGCCTTGTACACACCGCCCGTCACACCATGGGAGTGGGTTGCAAAAGAAGTAGGT
AGCTTAACCTCCGGGAGGGCGCT
>YER-30 | V1-V9 | 1426 bp
GGCCTAACACATGGCAAGTCGAGCGGCAGCGGGAAGTAGCTTGCTACTTTGCCGGCGAGCGGCGGACGGGTGAGTAATGTCTGGGA
AACTGCCTGATGGAGGGGGATAACTACTGGAAACGGTAGCTAATACCGCATGACCTCGCAAGAGCAAAGTGGGGGACCTTCGGGCC
TCACGCCATCGGATGTGCCCAGATGGGATTAGCTAGTAGGTGGGGTAATGGCTCACCTAGGCGACGATCCCTAGCTGGTCTGAGAG
GATGACCAGCCACACTGGAACTGAGACACGGTCCAGACTCCTACGGGAGGCAGCAGTGGGGAAATATTGCACCAATGGCGCAAGCC
TGATGCAGCCATGCCGCGGTGTGTGAAGAAGCTTAGGGTTGTAAAGCACTTCAGCGAGGAGGAAGGGTTCAGTGTAATAGCACTGT
ACATGACGTACTCGCAGAAGAAGCACGGCTAACTCCGTGCCAGCAGCCGCGGTAATACGGAGGGTGCAAGCGTTAATCGGAATTAC
TGGGCGTAAAGCGCACGCAGGCGGTTTGTTAAGTCAGATGTGAAATCCCCGAGCTTAACTTGGGAACTGCATTTGAAACTGGCAAG
CTAGAGTCTTGTAGAGGGGGGTAGAATTCCAGGTGTAGCGGTGAAATGCGTAGAGATCTGGAGGAATACCGGTGGCGAAGGCGGCC
CCCTGGACAAAGACTGACGCTCAGGTGCGAAAGCGTGGGGAGCAAACAGGATTAGATACCCTGGTAGTCCACGCTGTAAACGATGT
CGACTTGGAGGTTGTGCCCTTGAGGCGTGGCTTCCGGAGCTAACGCGTTAAGTCGACCGCCTGGGGAGTACGGCCGCAAGGTTAAA
ACTCAAATGAATTGACGGGGGCCCGCACAAGCGGTGGAGCATGTGGTTTAATTCGATGCAACGCGAAGAACCTTACCTACTCTTGA
CATCCAGAGAATTCGCTAGAGATAGCTTAGTGCCTTCGGGAACTCTGAGACAGGTGCTGCATGGCTGTCGTCAGCTCGTGTTGTGA

AATGTTGGGTTAAGTCCCGCAACGAGCGCAACCCTTATCCTTTGTTGCCAGCGAGTAATGTCGGGAACTCAAAGGAGACTGCCGGT
GATAAACCGGAGGAAGGTGGGGATGACGTCAAGTCATCATGGCCCTTACGAGTAGGGCTACACACGTGCTACAATGGCATATACAA
AGAGAAGCGAACTCGCGAGAGCAAGCGGACCTCATAAAGTATGTCGTAGTCCGGATTGGAGTCTGCAACTCGACTCCATGAAGTCG
GAATCGCTAGTAATCGTAGATCAGAATGCTACGGTGAATACGTTCCCGGGCCTTGTACACACCGCCCGTCACACCATGGGAGTGGG
TTGCAAAAGAAGTAGTGTAGCTTAACCTTCGGGAGGGCGCTACCACTTTG
>BAC-32 | V2-V8 | 1277 bp
GGCGGACGGGTGAGTAACACGTGGGTAACCTGCCTGTAAGACTGGGATAACTCCGGGAAACCGGGGCTAATACCGGATGCTTGTTT
GAACCGCATGGTTCAAACATAAAAGGTGGCTTTTCGCTACCACTTACAGATGGACCCGCGGCGCATTAGCTAGTTGGTGGGGTAAC
GGCTCACCAAGGCGACGATGCGTAGCCGACCTGAGAGGGTGATCGGCCACACTGGGACTGAGACACGGCCCAGACTCCTACGGGAG
GCAGCAGTAGGGAATCTTCCGCAATGGACGAAAGTCTGACGGAGCAACGCCGCGTGAGTGATGAAGGTTTTCGGATCGTAAAACTC
TGTTGTTAGGGAAGAACAAGTACCGTTCGAACAGGGCGGTACCTTGACGGTACCTAACCAGAAAGCCACGGCTAACTACGTGCCAG
CAGCCGCGGTAATACGTAGGTGGCAAGCGTTGTCCGGAATTATTGGGCGTAAAGCGCGCGCAGGCGGTTTTTTAAGTCTGATGTGA
AAGCCCCCGGCTCAACCGGGGAGGGTCATTGGAAACTGGGGAACTTGAGTGCAGAAGAGGAGAGTGGAATTCCACGTGTAGCGGTG
AAATGCGTAGAGATGTGGAGGAACACCAGTGGCGAAGGCGACTCTCTGGTCTGTAACTGACGCTGAGGCGCGAAAGCGTGGGGAGC
GAACAGGATTAGATACCCTGGTAGTCCACGCCGTAAACGATGAGTGCTAAGTGTTAGAGGGTTTCCGCCCTTTAGTGCTGCAGCAA
ACGCATTAAGCACTCCGCCTGGGGAGTACGGTCGCAAGACTGAAACTCAAAGGAATTGACGGGGGCCCGCACAAGCGGTGGAGCAT
GTGGTTTAATTCGAAGCAACGCGAAGAACCTTACCAGGTCTTGACATCCTCTGACAACCCTAGAGATAGGGCTTCCCCTTCGGGGG
CAGAGTGACAGGTGGTGCATGGTTGTCGTCAGCTCGTGTCGTGAGATGTTGGGTTAAGTCCCGCAACGAGCGCAACCCTTGATCTT
AGTTGCCAGCATTCAGTTGGGCACTCTAAGGTGACTGCCGGTGACAAACCGGAGGAAGGTGGGGATGACGTCAAATCATCATGCCC
CTTATGACCTGGGCTACACACGTGCTACAATGGGCAGAACAAAGGGCAGCGAAGCCGCGAGGCTAAGCCAATCCCACAAATCTGTT
CTCAGTTCGGATCGCAGTCTGCAACTCGACTGCGTGAAGCTGGAATCGCTAGTAATCGCGGATCAGCATGCC
>PSU-33 | V2-V9 | 1355 bp
GGCGGACGGGTGAGTAATGCCTAGGAATCTGCCTGGTAGTGGGGGATAACGTTCGGAAACGGAAGCTAATACCGCATACGTCCTAC
GGGAGAAAGCAGGGGACCTTCGGGCCTTGCGCTATCAGATGAGCCTAGGTCGGATTAGCTAGTTGGTGAGGTAATGGCTCACCAAG
GCGACGATCCGTAACTGGTCTGAGAGGATGATCAGTCACACTGGAACTGAGACACGGTCCAGACTCCTACGGGAGGCAGCAGTGGG
GAATATTGGACAATGGGCGAAAGCCTGATCCAGCCATGCCGCGTGTGTGAAGAAGGTCTTCGGATTGTAAAGCACTTTAAGTTGGG
AGGAAGGGCAGTTACCTAATACGTGATTGTTTTGACGTTACCGACAGAATAAGCACCGGCTAACTCTGTGCCAGCAGCCGCGGTAA
TACAGAGGGTGCAAGCGTTAATCGGAATTACTGGGCGTAAAGCGCGCGTAGGTGGTTAGTTAAGTTGGATGTGAAATCCCCGGGCT
CAACCTGGGAACTGCATTCAAAACTGACTGACTAGAGTATGGTAGAGGGTGGTGGAATTTCCTGTGTAGCGGTGAAATGCGTAGAT
ATAGGAAGGAACACCAGTGGCGAAGGCGACCACCTGGACTGATACTGACACTGAGGTGCGAAAGCGTGGGGAGCAAACAGGATTAG
ATACCCTGGTAGTCCACGCCGTAAACGATGTCAACTAGCCGTTGGGAGCCTTGAGCTCTTAGTGGCGCAGCTAACGCATTAAGTTG
ACCGCCTGGGGAGTACGGCCGCAAGGTTAAAACTCAAATGAATTGACGGGGGCCCGCACAAGCGGTGGAGCATGTGGTTTAATTCG
AAGCAACGCGAAGAACCTTACCAGGCCTTGACATCCAATGAACTTTCCAGAGATGGATTGGTGCCTTCGGGAACATTGAGACAGGT
GCTGCATGGCTGTCGTCAGCTCGTGTCGTGAGATGTTGGGTTAAGTCCCGTAACGAGCGCAACCCTTGTCCTTAGTTACCAGCACG
TAATGGTGGGCACTCTAAGGAGACTGCCGGTGACAAACCGGAGGAAGGTGGGGATGACGTCAAGTCATCATGGCCCTTACGGCCTG
GGCTACACACGTGCTACAATGGTCGGTACAGAGGGTTGCCAAGCCGCGAGGTGGAGCTAATCCCAGAAAACCGATCGTAGTCCGGA
TCGCAGTCTGCAACTCGACTGCGTGAAGTCGGAATCGCTAGTAATCGCGAATCAGAATGTCGCGGTGAATACGTTCCCGGGCCTTG
TACACACCGCCCGTCACACCCTGGGAGTGGGTTGCACCAGAAAGTAGTTTAGTCTAACCTTCGGG
>HAL-34 | V2-V9 | 1352 bp
GGCGGACGGGTGAGTAATGCATAGGAATCTGCCCGGTAGTGGGGGATAACCTGGGGAAACCCAGGCTAATACCGCATACGTCCTAC
GGGAGAAAGGGGGCTTCGGCTCCCGCTATTGGATGAGCCTATGTCGGATTAGCTAGTTGGTGAGGTAAAGGCTCACCAAGGCTGCG
ATCCGTAGCTGGTCTGAGAGGATGATCAGCCACATCGGGACTGAGACACGGCCCGAACTCCTACGGGAGGCAGCAGTGGGGAATAT

TGGACAATGGGGGCAACCCTGATCCAGCCATGCCGCGTGTGTGAAGAAGGCCCTCGGGTTGTAAAGCACTTTCAGCGAGGAAGAAC
GCCTAGCGGTTAATACCCGTTAGGAAAGACATCACTCGCAGAAGAAGCACCGGCTAACTCCGTGCCAGCAGCCGCGGTAATACGGA
GGGTGCAAGCGTTAATCGGAATTACTGGGCGTAAAGCGCGCGTAGGTGGCTTGATAAGCCGGTTGTGAAAGCCCCGGGCTCAACCT
GGGAACGGCATCCGGAACTGTCAGGCTAGAGTGCAGGAGAGGAAGGTAGAATTCCCGGTGTAGCGGTGAAATGCGTAGAGATCGGG
AGGAATACCAGTGGCGAAGGCGGCCTTCTGGACTGACACTGACACTGAGGTGCGAAAGCGTGGGTAGCAAACAGGATTAGATACCC
TGGTAGTCCACGCCGTAAACGATGTCGACCAGCCGTTGGGTGCCTAGCGCACTTTGTGGCGAAGTTAACGCGATAAGTCGACCGCC
TGGGGAGTACGGCCGCAAGGTTAAAACTCAAATGAATTGACGGGGGCCCGCACAAGCGGTGGAGCATGTGGTTTAATTCGATGCAA
CGCGAAGAACCTTACCTACCCTTGACATCTACAGAAGCCGGAAGAGATTCTGGTGTGCCTTCGGGAACTGTAAGACAGGTGCTGCA
TGGCTGTCGTCAGCTCGTGTTGTGAAATGTTGGGTTAAGTCCCGTAACGAGCGCAACCCTTGTCCTTATTTGCCAGCGAGTAATGT
CGGGAACTCTAAGGAGACTGCCGGTGACAAACCGGAGGAAGGTGGGGACGACTTCAAGTCATCATGGCCCTTACGGGTAGGGCTAC
ACACGTGCTACAATGGCCGGTACAAAGGGCTGCGAGCTCGCGAGAGTCAGCGAATCCCTTAAAGCCGGTCTCAGTCCGGATCGGAG
TCTGCAACTCGACTCCGTGAAGTCGGAATCGCTAGTAATCGTGAATCAGAATGTCACGGTGAATACGTTCCCGGGCCTTGTACACA
CCGCCCGTCACACCATGGGAGTGGACTGCACCAGAAGTGTTTTAGCTCTAACGCAAGAGGGC
>UNI-35 | V2-V3 | 709 bp
GGCGGAAGGGTGAGTAACAGGTGGGTAACCTGCCCATAAGACTGGGATAACTCCGGGAAACCGGGGCTAATACCGGATAACATTTT
GAACCGCATGGTTCGAAATTGAAAGGCGGCTTCGGCTGTCACTTATGGATGGACCCGCGTCGCATTAGCTAGTTGGTGAGGTAACG
GCTCACCAAGGCAACGATGCGTAGCCGACCTGAGAGGGTGATCGGCCACACTGGGACTGAGACACGGCCCAGACTCCTACGGGAGG
CAGCAGTAGGGAATCTTCCGCAATGGACGAAAGTCTGACGGAGCAACGCCGCGTGAGTGATGAAGGCTTTCGGGTCGTAAAACTCT
GTTGTTAGGGAAGAACAAGTGCTAGTTGAATAAGCTGGCACCTTGACGGTACCTAACCAAAAAGCCACGGCTAACTACGTGCCAGC
AGCCGCGGTAATACGTAGGTGGCAAGCGTTATCCGGAATTATTGGGCGTAAAGCGCGGGGCAGGTGGTTTCTTAATTCTGATGTGA
AACCCCACGGCTCAACCGTGGAGGGTCAGTGGAAACTGGGAGACTTGAGTGCAGAAGAGGAAAGTGCAATTCCATGTGTACCCATA
AGATGCGTGGAGATGTGGAGAAACACCCCGTGACCAAGGCGACTCTTCCCCCCTTTTGCCCCCCCATTAGGGACAAAGAGGAGGAA
AAAGAAGACGGTCTTGTTTT
>BAC-36 | V3-V7 | 1000 bp
ATTAGCTAGTTGGTGAGGTAACGGCTCACCAAGGCAACGATGCGTAGCCGACCTGAGAGGGTGATCGGCCACACTGGAACTGAAAC
CCCCCCCAGACTCCTACGGGAGGCACCAGTAGGGAATCTTCCGCAATGGACGAGGATCTGACGGAGCGACGCCGCGTGAGTGATAA
AAGCTTTCGGGTCGTAAAACTCTGTTGTTAGGGAAAAACAAGTGCTAGTTGAATAAGCTGGCTCCTTGAAGGTAGCTAACCCGAAT
CCCCCGGCTAACTACGTGCCAGCAGCCGCGGTAATAGGTAGTTGGCAAGCGTTATCCGGAACTATTGGGGGTAAAGCGCGCGCAGG
TGGTTTCTTAAGTTTGATGTGAAAGCGCACGGCTCAACCGTGGAGGGTCATTGGAAACTGGGAGACTTGAGTGCAGAAGAGGAAAG
TGGAATTCCATGTGTAGCGGTGAAATGCGTAGAGATATGGAGGAACACCAGTGGCGAAGGGACTTTCTGGTCTGTAACTGACACTG
AGGCGCGAAAGCGTGGGGAGCAAACAGGATTAGATACCCTGGTAGTCCACGCCGTAAACGATGAGTGCTAAGTGTTAGAGGGTTTC
CGCCCTTTAGTGCTGAAGTTAACGCATTAAGCACTCCGCCTGGGGAGTACGGCCGCAAGGCTGAAACTCAAAGGAATTGACGGGGG
CCCGCACAAGCGGTGGAGCATGTGGTTTAATTCGAAGCAACGCGAAGAACCTTACCAGGTCTTGACATCCTCTGAAAACCCTAGAG
ATAGGGCTTCTCCTTCGGGAGCAGAGTGACAGGTGGTGCATGGTTGTCGTCAGCTCGTGTCGTGAGATGTTGGGTTAAGTCCCGCA
ACGAGCGCAACCCTTGATCTTAGTTGCCATCATTAAGTTGGGCACTCTAAGGTGACTGCCGGTGACAAACCGGAGGAAGGTGGGGA
TGACGTCAAATCATCATGCCCCTTATGACCTGGGCTACACACGTGCTACAATGG
>UNI-37 | V2-V8 | 1268 bp
GGCGGACGGGTGAGTAACACGTGGGTAATCTGCCTGTAAGACGGGGATAACTCCGGGAAACCGGGGCTAATACCGGATAATAAGAG
AAGAAGCATTTCTTCTTTTTGAAAGTTGGTTTCGGCTGACACTTACAGATGAGCCCGCGGCGCATTAGCTAGTTGGTGAGGTAACG
GCTCACCAAGGCGACGATGCGTAGCCGACCTGAGAGGGTGATCGGCCACACTGGGACTGAGACACGGCCCAGACTCCTACGGGAGG
CAGCAGTAGGGAATCTTCGGCAATGGGCGAAAGCCTGACCGAGCAACGCCGCGTGAGCGATGAAGGCCTTCGGGTCGTAAAGCTCT
GTTGTTAGAGAAGAACAAGTACGAGAGTAACTGCTCGTACCTTGACGGTACCTAACCAGAAAGCCACGGCTAACTACGTGCCAGCA

GCCGCGGTAATACGTAGGTGGCAAGCGTTATCCGGAATTATTGGGCGTAAAGCGCGCGCAGGCGGTCTCTTAAGTCTGATGTGAAA
GCCCACGGCTCAACCGTGGAGGGTCATTGGAAACTGGGAGACTTGAGTGCAGGAGAGAAAAGTGGAATTCCACGTGTAGCGGTGAA
ATGCGTAGAGATGTGGAGGAACACCAGTGGCGAAGGCGGCTTTTTGGCCTGTAACTGACGCTGAGGCGCGAAAGCGTGGGGAGCAA
ACAGGATTAGATACCCTGGTAGTCCACGCCGTAAACGATGAGTGCTAGGTGTTGGGGGGTTCCACCCTCAGTGCTGAAGTTAACAC
ATTAAGCACTCCGCCTGGGGAGTACGACCGCAAGGTTGAAACTCAAAGGAATTGACGGGGGCCCGCACAAGCAGTGGAGCATGTGG
TTTAATTCGAAGCAACGCGAAGAACCTTACCAGGTCTTGACATCCTTTGACCACTCTAGAGATAGAGCTTTCCCCTTCGGGGGACA
AAGTGACAGGTGGTGCATGGTTGTCGTCAGCTCGTGTCGTGAGATGTTGGGTTAAGTCCCGCAACGAGCGCAACCCTTGACCTTAG
TTGCCAGCATTCAGTTGGGCACTCTAAGGTGACTGCCGGTGACAAACCGGAGGAAGGTGGGGATGACGTCAAATCATCATGCCCCT
TATGACCTGGGCTACACACGTGCTACAATGGATGATACAAAGGGTTGCGAAGCCGCGAGGCCAAGCCAATCCCAAAAAGTCATTCT
CAGTTCGGATTGTAGGCTGCAACTCGCCTACATGAAGCCGGAATTGCTAGTAATCGCGGATCAG
>STA-38 | V3-V8 | 1090 bp
GGTAACGGCTTACCAAGGCAACGATACGTATCCGACCTGAGAGGGTGATCGGCCACACTGGAACTGAGACACGGTCCAGACTCCTA
CGGGAGGCAGCAGTAGGGAATCTTCCGCAATGGGCGAAAGCCTGACGGAGCAACGCCGCGTGAGTGATGAAGGTCTTCGGATCGTA
AAACTCTGTTATCAGGGAAGAACAAACGTGTAAGTAACTGTGCACGTCTTGACGGTACCTGATCAGAAAGCCACGGCTAACTACGT
GCCAGCAGCCGCGGTAATACGTAGGTGGCAAGCGTCATCCGGAATTATTGGGCGTAAAGCGCGCGTAGGCGGCTTTTTTAAGTCTG
ATGTGAAAGCCCACGGCTCACCCGTGGAGGGTCATTGGAAACTGGAAAACTTGAGTGCAGAAGAGGAAAGTGGAATTCCATGTGTA
GCGGTGAAATGCGCAGAGATATGGAGGAACACCAGTGGCGAAGGCGACTTTCTGGTCTGTAACTGACGCTGATGTGCGAAAGCGTG
GGGATCAAACAGGATTAGATACCCTGGTAGTCCACGCCGTAAACGATGAGTGCTAAGTGTTAGGGGGTTTCCGCCCCTTAGTGCTG
CAGCTAACGCATTAAGCACTCCGCCTGGGGAGTACGACCGCAAGGTTGAAACTCAAAGGAATTGACGGGGACCCGCACAAGCGGTG
GAGCATGTGGTTTAATTCGAAGCAACGCGAAGAACCTTACCAAATCTTGACATCCTTTGACCGCTCTAGAGATAGAGTTTTCCCCT
TCGGGGGACAAAGTGACAGGTGGTGCATGGTTGTCGTCAGCTCGTGTCGTGAGATGTTGGGTTAAGTCCCGCAACGAGCGCAACCC
TTAAGCTTAGTTGCCATCATTAAGTTGGGCACTCTAAGTTGACTGCCGGTGACAAACCGGAGGAAGGTGGGGATGACGTCAAATCA
TCATGCCCCTTATGATTTGGGCTACACACGTGCTACAATGGACAATACAAAGGGCAGCTAAACCGCGAGGTCAAGCAAATCCCATA
AAGTTGTTCTCAGTTCGGATTGTAGTTTGCAACTCGACCACATGAAGTAGGAATCGCT
>BAC-39 | V3-V5 | 707 bp
TTAGCTAGTTGGTGAGGTAACTGCTCACCAAGGCAACAATGCAGAGCCAACCTGAGAGGGTGATCGGCCACACTGGGACTGAAACC
CCGCCCATACTCCTACGGGAGGGAGCAGTATGGAATCTTCCGCAATGGACGAAAGTCTGACGGAACAGGGGGGGTGTGTGATGAAG
GCTTTCCGTTCGTAAAACTCTGTTGTTAGGAAAAAACAAATGCGAAAGTAGCTGCTTGTAGCTTGACGGTACCTAACAGAATTTCC
ACGGCTAACTACGTGCCAGCAGCCGCGAAAATACGTAGGTGGCGAGCGTTATCCGGAATTATTGGGCGTAAAGCGCGCGGAGGGGG
CTTCATAAGTCTGATGTGAAAGCCCGCGGCCCAACCCTGCAGGGGCATTGGAAACTGGGGAGCTTGAGTGGCAAGAGAGAACCGGA
ATTCCACGTGTAGCAGTGAAATGCTTAAGGTGTGGAGGAACGCCGGTGGCTAAAGCGGTTTTTTCCTCTGTATCTCACCCTCAGCG
CGAAAGCATGGGGAAAGGCGGCATTAGACACCCTGGTTCTCCCCTCCTTAAGCGATTTCTGCGAACTGTTAGGGGGTTCCCCTTCT
TCTTTCGGGCACCTCACGCGCTAATCACTCCGGCCGGCCAGTGGGGGGGCCCGGGGGGGTTCACAAAGAGTTTAAAGGGGCCCCCA
CAAGCGCTTGAGCATGTG
>MIC-40 | V3-V7 | 873 bp
CTACGGACGGCAGCAGTTGGGAATATGCACAATGGCGCATGCTGATGCAGCGACGCGTGTGAGGGACGACGGCTTTGGGTGTAACC
TCTTCAGTAGGAAGAAGGGAAATGACGGTACTGCAGATGCAGCACCGGGTAACTACGTGCCAGCAGCCGCGGTAATATGTAGGGTG
CGAGCGTTATCTGGAATTATTGGGTGTAAAGAGCTGTAGGCGGTTGTCGCGTCTGTCGTGAATGTCCGGGGCTTAACCCTGGATCC
GCGGTGGGTACGGGCAGACTAGAGTGCAGTAGGGGAGAGTGGAATTCCTGGTGTAGCGGTGGAATGCGCAGATATCAGGAGGAACA
CCGATGGCGAAGGCAGGTCTCTGGGCTGTAACTGACGCTGAGGAGCGAAAGCATGGGGAGCGAACAGGATTAGATACCTTGGTAGT
CCATGCCGTAAACGTTGGGCACTAGGTGTGGGGACCATTCCACGGTTTCCGCGCCGCAGCTAACGCATTAAGTGCCCCGCCTGGGG
AGTACGGCCGCAAGGCTAAAACTCAAAGGAATTGACGGGGGCCCGCACAAGCGGCGGAGCATGCGGATTAATTCGATGCAACGCGA

AGAACCTTACCAAGGCTTGACATGTTCTCGATCGCCGTAGAGATACGGTTTCCCCTTTGGGGCGGGTTCACAGGTGGTGCATGGTT
GTCGTCAGCTCGTGTCGTGAGATGTTGGGTTAAGTCCCGCAACGAGCGCAACCCTCGTTCCATGTTGCCAGCACGTAATGGTGGGG
ACTCATGGGAGACTGCCGGGGTCAACTCGGAGGAAGGTGAGGACGACGTCAAATCATCATGCCCCTTATGTCTTGGGCTTCACGCA
TGCTACAATGGC
>PSU-41 | V2-V9 | 1357 bp
GGCGGACGGGTGAGTAATGCCTAGGAATCTGCCTGGTAGTGGGGGATAACGTTCGGAAACGGACGCTAATACCGCATACCTCCTAC
GGGAGAAAGCAGGGGACCTTCGGGCCTTGCGCTATCAGATGAGCCTAGGTCGGATTAGCTAGTTGGTGGGGTAATGGCTCACCAAG
GCGACGATCCGTAACTGGTCTGAGAGGATGATCAGTCACACTGGAACTGAGACACGGTCCAGACTCCTACGGGAGGCAGCAGTGGG
GAATATTGGACAATGGGCGAAAGCCTGATCCAGCCATGCCGCGTGTGTGAAGAAGGTCTTCGGATTGTAAAGCACTTTAAGTTGGG
AGGAAGAGCAGTTACCTAATACGTGATTGTTTTGACGTTACCGACAGAATAAGCACCGGCTAACTCTGTGCCAGCAGCCGCGGTAA
TACAGAGGGTGCAAGCGTTAATCGGAATTACTGGGCGTAAAGCGCGCGTAGGTGGTTTGTTAAGTTGGATGTGAAATCCCGGGGCC
TCAACCTGGGAACTGCATTCAAAACTGACTGACTAGAGTATGGTAGAGGGTGGTGGAATTTCCTGTGTAGCGGTGAAATGCGTAGA
TATAGGAAGGAACACCAGTGGCGAAGGCGACCACCTGGACTAATACTGACACTGAGGTGCGAAAGCGTGGGGAGCAAACAGGATTA
GATACCCTGGTAGTCCACGCCGTAAACGATGTCAACTAGCCGTTGGAAGCCTTGAGCTTTTAGTGGCGCAGCTAACGCATTAAGTT
GACCGCCTGGGGAGTACGGCCGCAAGGTTAAAACTCAAATGAATTGACGGGGGCCCGCACAAGCGGTGGAGCATGTGGTTTAATTC
GAAGCAACGCGAAGAACCTTACCAGGCCTTGACATCCAATGAACTTTCTAGAGATAGATTGGTGCCTTCGGGAACATTGAGACAGG
TGCTGCATGGCTGTCGTCAGCTCGTGTCGTGAGATGTTGGGTTAAGTCCCGTAACGAGCGCAACCCTTGTCCTTAGTTACCAGCAC
GTAATGGCGGGCACTCTAAGGAGACTGCCGGTGACAATCCGGAGGAAGGTGGGGATGATGTCAAGTCATCATGGCCCTTACGGCCT
GGGCTACACACGTGCTACAATGGTCGGTACAGAGGGTTGCCAAGCCGCGAGGTGGAGGTAATCCCACAAAACCGATCGTAGTCCGG
ATCGCAGTCTGCAACTCGACTGCGTGAAGTCGGAATCGCTAGTAATCGCGAATCAGAATGTCGCGGTGAATACGTTCCCGGGCCTT
GTACACACCGCCCGTCACACCATGGGAGTGGGTTGCACCAGAAGTAGTTAGTCTAACCTTCGGGAGG
>PSU-42 | V2-V8 | 1257 bp
GGCGGACGGGTGAGTAATGCCTAGGAATCTGCCTAGTAGTGGGGGATAACTTGCCGAAAGGTAAGCTAATACCGCAAACGTCCTAC
GGGAGAAAGGAGGGGACCTTCGGGCCTTTCGCTATTAGATGAGCCTAGGTCGGATTAGTTAGTTGGTGAGGTAATGGCTCACCAAG
ACCGCGATCCGTAACTGGTCTGAGAGGATGATCAGTCACACTGGAACTGAGACACGGTCCAGACTCCTACGGGAGGCAGCAGTGGG
GGAATATTGGACAATGGGGGAACCCTGATCCAGCATGCCGCGTGTGTGAAGAAGTCTAGGATGTAAAGCACTTTAAGTGGGAGGAAG
AGCAGTTAACTAATATTTGACTGTTTTGACGTTACCGACAGAATAAGCACGGCTAACTTCGTGCCAGCAGCCGCGGTAATACGAAG
GGTGCAGCGTTAATCGGAATTACTGGGCGTAAAGCGCGCGTAGGTGGTTCAGTAAGTTAGGAGTGAAAGCCCCGGGCTTAACCTGG
GAATTGCTTCTAAAACTGCTGAGCTAGAGTACGGTAGAGGGTGGTGGAATTTCCTGTGTAGCGGTGAAATGCGTAGATATAGGAAG
GAACATCAGTGGCGAAGGCGACCACCTGGACTGATACTGACACTGAGGTGCGAAAGCGTGGGGAGCAAACAGGATTAGATACCCTG
GTAGTCCACGCCGTAAACGATGTCAACTAGTTGTTGGGCTCCTTGAGGACTTAGTAACGCAGCTAACGCATTAAGTTGACCGCCTG
GGGAGTACGGCCGCAAGGTTAAAACTCAAATGAATTGACGGGGGCCCGCACAAGCGGTGGAGCATGTGGTTTAATTCGAAGCAACG
CGAAGAACCTTACCTGGCCTTGACATGCTGAGAACTTTCCAGAGATGGATTGGTGCCTTCGGGAACTCAGACACAGGTGCTGCATG
GCTGTCGTCAGCTCGTGTCGTGAGATGTTGGGTTAAGTCCCGTAACGAGCGCAACCCTTGTCCTTATTTACCAGCACGTAATGGTG
GGCACTCTAAGGAGACTGCCGGTGACAAACCGGAGGAAGGTGGGGATGACGTCAAGTCATCATGGCCCTTACGGCCAGGGCTACAC
ACGTGCTACAATGGTTGGTACAACGGGCTGCCAAGTCGCGAGACGGAGCTAATCCCATAAAACCAATCGTAGTCCGGATCGCAGTC
TGCAACTCGACTGCGTGAAGTCGGAATCGCTAGTAATCGTGGATCAGAATGCC
>BAC-43 | V2-V9 | 1360 bp
GGCGGACGGGTGAGTAACACGTGGGTAACCTGCCCATAAGACTGGGATAACTCCGGGAAACCGGGGCTAATACCGGATAACATTTT
GAACTGCATGGTTCGAAATTGAAAGGCGGCTTCGGCTGTCACTTATGGATGGACCCGCGTCGCATTAGCTAGTTGGTGAGGTAACG
GCTCACCAAGGCAACGATGCGTAGCCGACCTGAGAGGGTGATCGGCCACACTGGGACTGAGACACGGCCCAGACTCCTACGGGAGG
CAGCAGTAGGGAATCTTCCGCAATGGACGAAAGTCTGACGGAGCAACGCCGCGTGAGTGATGAAGGCTTTCGGGTCGTAAAACTCT

GTTGTTAGGGAAGAACAAGTGCTAGTTGAATAAGCTGGCACCTTGACGGTACCTAACCAGAAAGCCACGGCTAACTACGTGCCAGC
AGCCGCGGTAATACGTAGGTGGCAAGCGTTATCCGGAATTATTGGGCGTAAAGCGCGCGCAGGTGGTTTCTTAAGTCTGATGTGAA
AGCCCACGGCTCAACCGTGGAGGGTCATTGGAAACTGGGAGACTTGAGTGCAGAAGAGGAAAGTGGAATTCCATGTGTAGCGGTGA
AATGCGTAGAGATATGGAGGAACACCAGTGGCGAAGGCGACTTTCTGGTCTGTAACTGACACTGAGGCGCGAAAGCGTGGGGAGCA
AACAGGATTAGATACCCTGGTAGTCCACGCCGTAAACGATGAGTGCTAAGTGTTAGAGGGTTTCCGCCCTTTAGTGCTGAAGTTAA
CGCATTAAGCACTCCGCCTGGGGAGTACGGCCGCAAGGCTGAAACTCAAAGGAATTGACGGGGGCCCGCACAAGCGGTGGAGCATG
TGGTTTAATTCGAAGCAACGCGAAGAACCTTACCAGGTCTTGACATCCTCTGAAAACCCTAGAGATAGGGCTTCTCCTTCGGGAGC
AGAGTGACAGGTGGTGCATGGTTGTCGTCAGCTCGTGTCGTGAGATGTTGGGTTAAGTCCCGCAACGAGCGCAACCCTTGATCTTA
GTTGCCATCATTAAGTTGGGCACTCTAAGGTGACTGCCGGTGACAAACCGGAGGAAGGTGGGGATGACGTCAAATCATCATGCCCC
TTATGACCTGGGCTACACACGTGCTACAATGGACGGTACAAAGAGCTGCAAGACCGCGAGGTGGAGCTAATCTCATAAAACCGTTC
TCAGTTCGGATTGTAGGCTGCAACTCGCCTACATGAAGCTGGAATCGCTAGTAATCGCGGATCAGCATGCCGCGGTGAATACGTTC
CCGGGCCTTGTACACACCGCCCGTCACACCACGAGAGTTTGTAACACCCGAAGTCGGTGGGGTAACCTT
>BAC-44 | V2-V9 | 1358 bp
GGCGGACGGGTGAGTAACACGTGGGTAACCTGCCTGTAAGACTGGGATAACTCCGGGAAACCGGAGCTAATACCGGATAGTTCCTT
GAACCGCATGGTTCAAGGATGAAAGACGGTTTCGGCTGTCACTTACAGATGGACCCGCGGCGCATTAGCTAGTTGGTGAGGTAACG
GCTCACCAAGGCGACGATGCGTAGCCGACCTGAGAGGGTGATCGGCCACACTGGGACTGAGACACGGCCCAGACTCCTACGGGAGG
CAGCAGTAGGGAATCTTCCGCAATGGACGAAAGTCTGACGGAGCAACGCCGCGTGAGTGATGAAGGTTTTCGGATCGTAAAGCTCT
GTTGTTAGGGAAGAACAAGTGCAAGAGTAACTGCTTGCACCTTGACGGTACCTAACCAGAAAGCCACGGCTAACTACGTGCCAGCA
GCCGCGGTAATACGTAGGTGGCAAGCGTTGTCCGGAATTATTGGGCGTAAAGGGCTCGCAGGCGGTTTCTTAAGTCTGATGTGAAA
GCCCCCGGCTCAACCGGGGAGGGTCATTGGAAACTGGGAAACTTGAGTGCAGAAGAGGAGAGTGGAATTCCACGTGTAGCGGTGAA
ATGCGTAGAGATGTGGAGGAACACCAGTGGCGAAGGCGACTCTCTGGTCTGTAACTGACGCTGAGGAGCGAAAGCGTGGGGAGCGA
ACAGGATTAGATACCCTGGTAGTCCACGCCGTAAACGATGAGTGCTAAGTGTTAGGGGGTTTCCGCCCCTTAGTGCTGCAGCTAAC
GCATTAAGCACTCCGCCTGGGGAGTACGGTCGCAAGACTGAAACTCAAAGGAATTGACGGGGGCCCGCACAAGCGGTGGAGCATGT
GGTTTAATTCGAAGCAACGCGAAGAACCTTACCAGGTCTTGACATCCTCTGACAACCCTAGAGATAGGGCTTTCCCTTCGGGGACA
GAGTGACAGGTGGTGCATGGTTGTCGTCAGCTCGTGTCGTGAGATGTTGGGTTAAGTCCCGCAACGAGCGCAACCCTTGATCTTAG
TTGCCAGCATTCAGTTGGGCACTCTAAGGTGACTGCCGGTGACAAACCGGAGGAAGGTGGGGATGACGTCAAATCATCATGCCCCT
TATGACCTGGGCTACACACGTGCTACAATGGACAGAACAAAGGGCTGCGAGACCGCAAGGTTTAGCCAATCCCACAAATCTGTTCT
CAGTTCGGATCGCAGTCTGCAACTCGACTGCGTGAAGCTGGAATCGCTAGTAATCGCGGATCAGCATGCCGCGGTGAATACGTTCC
CGGGCCTTGTACACACCGCCCGTCACACCACGAGAGTTTGCAACACCCGAAGTCGGTGAGGTAACCTT
